# Economic hardship mediates differences in brain network dynamics associated with family history of substance use disorder in a sex-specific manner

**DOI:** 10.64898/2026.10.14.759285

**Authors:** Louisa Schilling, Marie Hédo, Keith Jamison, Christie Gillies, Ana Radanovic, Qingyu Zhao, Amy Kuceyeski

## Abstract

Family history of substance use disorder (SUD) is associated with brain differences before substance exposure, but the pathways linking familial risk to brain function may differ by sex. Using baseline data from the Adolescent Brain Cognitive Development (ABCD) Study, we tested whether environmental adversity mediates previously observed sex-specific associations between family history of SUD and transition energy, a network control theory-derived measure of the energetic cost of transitioning between brain activity states. We derived three environmental dimensions—economic hardship, family–school neglect, and parental mental illness—and fit sex-stratified mediation models at global, network, and regional scales. Environmental mediation was significant in females and was driven primarily by economic hardship, which mediated higher transition energy across default mode, visual, ventral attention, and somatomotor networks, with widespread positive regional effects. In males, mediation was weaker and more heterogeneous, with more negative transition-energy patterns that were less explained by the measured environmental pathways. In an exploratory extension, state policy environments associated with smaller FH+ versus FH*−* economic-hardship gaps were projected to partially attenuate FH-related transition-energy gaps, especially in females. These findings suggest that familial SUD risk is embedded in brain dynamics through sex-distinct pathways, with economic hardship representing a potentially modifiable pathway linking familial risk to large-scale brain function in early adolescence.

## 1 Introduction

Approximately 17% of the U.S. population aged 12 years and older met criteria for a substance use disorder (SUD) in 2024 [1]. SUD is characterized by compulsive substance use despite harmful consequences. Although exposure to addictive substances is necessary for SUD to develop, only a minority of individuals who use substances progress to the disorder [2, 3]. This underscores the importance of identifying risk factors that confer vulnerability, particularly those that may be amenable to intervention [2, 3].

Across drug classes, addictive substances act on neural systems involved in reward processing and inhibitory control, and SUD is consistently associated with disruption of these circuits [4]. Vulnerability to SUD has likewise been linked to alterations in these same systems. In adolescence, such vulnerability is thought to arise in part from an imbalance in neurodevelopment, whereby reward-related systems mature more rapidly than top-down control systems [5–9]. This susceptibility may be further amplified in youth with a family history of SUD (FH+) relative to those without such a history (FH*−*), through the interplay of genetic and environmental influences [10–12]. Consequently, it is often difficult to disentangle neural features that reflect vulnerability preceding substance use from those that arise as a consequence of substance use, as well as to separate genetic from environmental contributions to risk. Adding further complexity, sex assigned at birth, hereafter referred to as sex, has repeatedly been shown to shape behavioral and neural alterations related to SUD and SUD risk [13–18].

Consistent with an elevated vulnerability prior to substance exposure, substance-naive youth with SUD family history exhibit neurodevelopmental alterations in systems supporting reward processing and behavioral control. Although much of this literature has focused on regional differences in brain structure and function, evidence from both SUD and populations at risk for SUD suggests that these alterations extend to the large-scale organization and dynamics of brain networks [19–25]. A network-level perspective is critical for understanding SUD risk, as the disorder appears to involve dysfunction across multiple interacting circuits involved in cognitive control, salience processing, and reward-related behavior [26–30]. Network control theory (NCT) provides a framework to characterize these large-scale brain dynamics by estimating the energy required to transition between brain activity states given the brain’s underlying structural connectivity, defined by white matter tract connections between regions. This transition energy measure provides a systems-level index of brain function that may be sensitive to early risk-related differences [29, 31, 32].

Family history of SUD captures both inherited genetic liability and adverse environmental context. Genetic studies indicate that SUD has a substantial heritable component. Twin and family studies typically estimate heritability at roughly 50%, whereas GWAS-based estimates of SNP heritability are considerably lower, often around 10%, suggesting that common genetic variants explain only a modest proportion of overall liability [10, 33, 34]. Meanwhile, environmental exposures closely linked to family history of SUD—including socioeconomic adversity, neglect, family stress, household instability, prenatal substance exposure, trauma, and parental mental illness—are also known to shape neurodevelopment and influence risk, both independently and through interaction with genetic liability [10–13, 35]. Thus, interpreting family-history-related brain differences requires considering both inherited and environmental pathways.

Prior work further suggests that the relative contributions of genetic and environmental influences to SUD risk may differ by sex, with some evidence of greater environmental sensitivity in females [13, 15, 36]. Socioeconomic status (SES) and family adversity have likewise been linked to sex-dependent effects on brain development, in some cases driving morphometric or functional measures in opposite directions in males and females [37, 38]. Such sexually divergent responses to the environment during early adolescence likely reflect, in part, concurrent neuroendocrine and neurodevelopmental changes that vary by sex. Early pubertal timing is associated with environmental stress and low SES in both sexes, though more strongly in females [39–42], and may therefore interact with adverse environments in a sex-specific manner to shape neurodevelopmental trajectories [43–45]. Furthermore, there is limited evidence that heritability estimates for SUD are higher in males than in females, although findings remain mixed [36, 46, 47]. Disentangling these pathways is essential for understanding how SUD risk is transmitted and why its neural correlates diverge across sexes, with implications for more targeted prevention efforts.

Our previous work applied NCT to resting-state fMRI (rsfMRI) and diffusion MRI (dMRI) data from the Adolescent Brain Cognitive Development (ABCD) Study [48] and showed that FH+ youth aged 9–10 years exhibited sex-specific alterations in brain network dynamics relative to FH*−* youth, prior to substance exposure [49]. Specifically, FH+ females required greater transition energy in higher-order networks, particularly the default mode network (DMN), whereas FH+ males required lower transition energy in lower-order networks, especially the dorsal and ventral attention networks (DAT and VAT), relative to their within-sex FH*−* peers. These distinct neural profiles may reflect known behavioral divergences in trajectories of SUD risk [15]. Females are more likely to follow stress-related, internalizing pathways to SUD, often involving negative affect and rumination, processes that may be reflected in elevated energetic demands within the DMN. By contrast, males more often follow impulsivity-driven, externalizing pathways marked by attentional deficits and behavioral disinhibition, which may be reflected in reduced energetic barriers within attentional networks [13, 50]. However, that study did not address which components of family history—inherited liability, environmental adversity, or their interplay—may drive these sex-specific neural differences.

Here, we extend these previous findings by testing whether environmental factors mediate the relationship between family history of SUD and brain network dynamics, and whether this mediation differs by sex. We hypothesized that environmental mediation of the effect of family history would be stronger in females, reflecting heightened sensitivity to early-life context, whereas risk in males would be characterized by more direct, less environmentally mediated effects. To capture environmental complexity in a data-driven manner, we used principal component analysis (PCA) to derive three broad dimensions of environmental adversity: economic hardship, family–school neglect, and parental mental illness. We then applied structural equation modeling (SEM) to NCT-derived metrics from the baseline ABCD Study cohort (*N* = 1,835) to test whether these environmental dimensions explain sex-specific variation in transition energy. SEM is a statistical framework that estimates a system of related relationships simultaneously, allowing both direct associations and indirect pathways through mediating variables to be examined within a single model [51]. Finally, because economic hardship emerged as the primary environmental mediator, we conducted an exploratory policy-linked projection analysis in the broader ABCD cohort to ask whether state-level policy environments associated with smaller FH+ versus FH*−* economic-hardship gaps would be expected to partially attenuate FH-related TE gaps through the economic hardship pathway. By mapping these pathways at global, network, and regional scales, and contextualizing environmental pathways in relation to broader policy environments, we sought to clarify how familial SUD risk becomes embedded in the developing brain and to identify potentially modifiable targets for prevention.

## 2 Results

### 2.1 Sample Characteristics

We began our analysis with the full ABCD baseline cohort (N = 11,868; age 8.9–11.1 years; 47.8% female, Release 5.1) to derive broad environmental dimensions of risk using principal component analysis. Subsequent mediation analyses were performed on a subsample (N = 1,835; age 8.9–11.0 years; 54.4% female). Participants were excluded from the full cohort if they had unusable MRI data, had used substances, were adopted, did not meet criteria for FH+ or FH*−* for SUD (see Section 5.1.3), or were missing required covariates (i.e., age, sex, race/ethnicity, puberty, or family ID). See Section 5.1.2 for details on exclusions. Key demographics, covariates, and outcomes in this subsample are summarized in Table 1. The four sex-by-family history groups differed in framewise displacement, global TE, all three environmental PCs, race/ethnicity, and pubertal initiation, as assessed using one-way ANOVAs for continuous variables and *χ*^2^ tests for categorical variables (*p <* 0.05); age and MRI scanner manufacturer did not differ significantly across groups (Table 1).

**Table 1:** Mediation analytic sample by family history and sex. Values are mean (SD) for continuous variables and *n* (%) for categorical levels. *p*: one-way ANOVA across the four sex*×*FH groups (continuous) or Pearson *χ*^2^ test (categorical).

| | | FH− male<br>$n = 623$ | FH− female<br>$n = 721$ | FH+ male<br>$n = 214$ | FH+ female<br>$n = 277$ | $p$ |
| --- | --- | --- | --- | --- | --- | --- |
| Age (years) |  | 10.04 (0.63) | 10.00 (0.62) | 10.00 (0.63) | 9.98 (0.61) | 0.437 |
| Mean FD (mm) |  | 0.082 (0.030) | 0.078 (0.027) | 0.083 (0.030) | 0.081 (0.028) | 0.015 |
| Global TE (z-score) |  | -0.04 (1.10) | 0.05 (0.96) | -0.21 (0.82) | 0.13 (0.94) | < 0.001 |
| Economic Hardship (PC1) |  | -0.82 (1.72) | -0.83 (1.88) | 0.63 (2.19) | 0.66 (2.17) | < 0.001 |
| Family-School Neglect (PC2) |  | -0.16 (1.53) | -0.48 (1.45) | 0.70 (1.62) | 0.49 (1.60) | < 0.001 |
| Parental Mental Illness (PC3) |  | -0.44 (1.31) | -0.03 (1.29) | 0.46 (1.61) | 0.84 (1.38) | < 0.001 |
| Puberty (started) |  | 153 (24.6%) | 479 (66.4%) | 60 (28.0%) | 194 (70.0%) | < 0.001 |
| MRI manufacturer | Siemens | 460 (73.8%) | 527 (73.1%) | 143 (66.8%) | 187 (67.5%) | 0.072 |
|  | GE | 163 (26.2%) | 194 (26.9%) | 71 (33.2%) | 90 (32.5%) |  |
| Race/ethnicity | Black | 30 (4.8%) | 66 (9.2%) | 21 (9.8%) | 33 (11.9%) | 0.003 |
|  | Hispanic | 102 (16.4%) | 139 (19.3%) | 36 (16.8%) | 58 (20.9%) |  |
|  | Other | 76 (12.2%) | 95 (13.2%) | 32 (15.0%) | 31 (11.2%) |  |
|  | White | 415 (66.6%) | 421 (58.4%) | 125 (58.4%) | 155 (56.0%) |  |

TE = transition energy. FD = framewise displacement. PC = principal component. Other includes Asian participants and all participants not classified as White, Black, or Hispanic.

### 2.2 Latent environmental dimensions are associated with a family history of SUD

To capture the primary environmental dimensions, PCA was performed on 43 variables selected based on established risk factors for SUD, spanning socioeconomic, household, neighborhood, school, peer, parenting, family climate, trauma, parental mental health, pregnancy, and prenatal substance use domains, in the full ABCD baseline cohort. Details of PCA variable selection and pre-processing are provided in Supplementary Section S2.1. The first three components explained 23.2% of the variance (PC1: 11.26%, PC2: 6.40%, PC3: 5.56%), with subsequent PCs each explaining less than 5% of the variance (Fig. 1A). The first three PCs were therefore retained for further analyses and oriented such that higher values reflected greater environmental SUD risk. This re-orientation was for interpretability and did not alter analyses.

**Figure 1.**
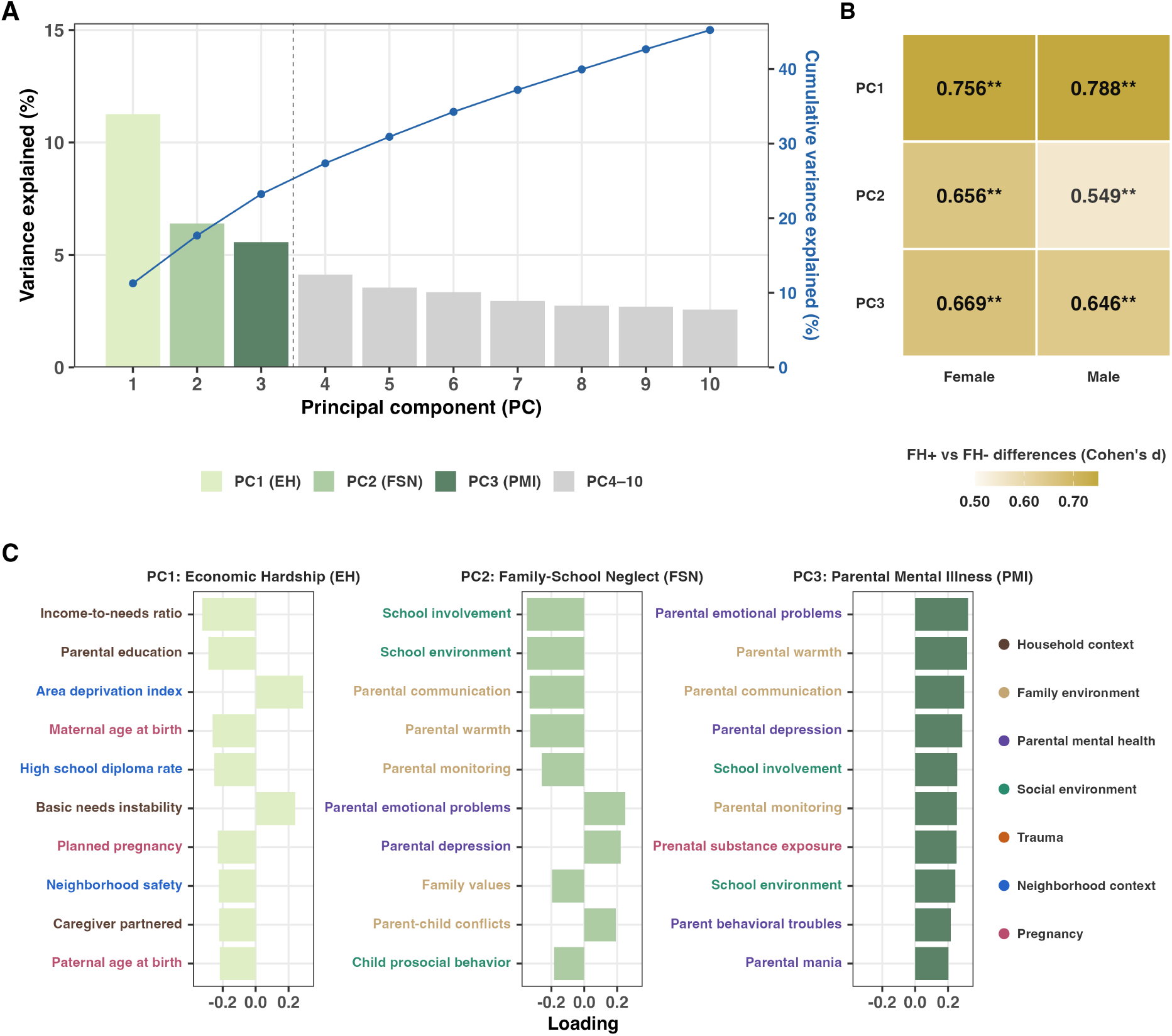
Latent dimensions of environmental risk. **(A)** Scree plot of the first ten principal components (PCs) from PCA of baseline environmental factors in the ABCD Study (Release 5.1; *N* = 11,868); PC1–PC3 were retained for mediation analyses. **(B)** Standardized mean differences (Cohen’s *d*) in PC scores between FH+ and FH*−* youth, shown separately by sex; *P*-values were FDR-adjusted across PC*×*sex comparisons. **(C)** Oriented loadings for PC1–PC3, labeled as economic hardship (EH), family–school neglect (FSN), and parental mental illness (PMI), showing the ten variables with the largest absolute loadings. In **(C)**, higher PC loading scores indicate greater family history-related risk. *^∗^p <* 0.05; *^∗∗^p*_FDR_ *<* 0.05.

Inspection of the loadings revealed interpretable domains for each of the first three components. PC1 reflected economic hardship (EH), characterized by greater neighborhood deprivation and lower household socioeconomic status. PC2 captured family–school neglect (FSN), reflecting reduced parental support, poorer school environment, and greater family disruption. PC3 indexed parental mental illness (PMI), with strong loadings on parental mental health symptoms and maternal prenatal substance use, although it also showed contributions from family- and school-support measures (Fig. 1C). Later components were less interpretable, showed weaker associations with family history of SUD, and explained less variance. See Supplementary Figures S9 and S10 for more details on the top 10 PCs.

Together, these findings identify three primary environmental dimensions: economic hardship, family–school neglect, and parental mental illness, which were included as parallel mediators in subsequent mediation analyses.

In the subsample retained after exclusions (*N* = 1,835), FH+ individuals had significantly higher scores than FH*−* individuals on each of the first three components in both sexes (Fig. 1B). Effect sizes for the FH+ versus FH*−* contrast were largest for PC1 (female *d* = 0.76; male *d* = 0.79) and remained moderate to large for PC2 and PC3 (range, *d* = 0.55–0.67), with all differences surviving multiple-comparison correction in both sexes (all *p*_FDR_ *<* 10*^−^*^9^). Together, these findings indicate robust associations between family history of SUD and multiple dimensions of environmental risk.

Although we refer to these PCs as *environmental* dimensions for simplicity, they likely also index heritable or biological liability, particularly PC3 (PMI), which showed strong loadings on parental mental health and maternal substance use alongside family and school variables. Importantly, all three dimensions nevertheless capture meaningful aspects of the broader risk context associated with family history of SUD.

### 2.3 TE calculation

Because our goal was to investigate environmental mediators of our previous findings of sex differences in brain dynamics among youth with a family history of SUD, we computed TE measures using the same approach as in our prior study [49]. Briefly, resting-state functional MRI (rsfMRI) and diffusion MRI were processed using previously published pipelines to extract regional time series and structural connectivity networks, based on the 68-region Desikan–Killiany atlas plus 16 subcortical and 2 cerebellar regions [52]. To calculate TE, we first identified recurring brain states by applying *k*-means clustering to regional rsfMRI time series (*k* =4). Consistent with our previous study, these four states comprised two anti-correlated meta-state pairs, which we labeled according to the network showing the greatest cosine similarity in patterns of high- and low-amplitude activity relative to the mean: DMN+, DMN*−*, VIS+, and VIS*−*. See Supplementary Figure S6 for details on clustering and a visualization of the four brain states. Using a group-average structural connectome calculated in a subset of 149 participants and subject-specific cluster centroids derived from each subject’s rsfMRI data, we computed the minimum energy required to transition between all pairs of brain states using our previously described NCT framework. In short, NCT models brain activity over time as a diffusion process on the structural connectome with a term for external input. The system has a closed-form solution that is used to calculate the amount of external input (energy) needed to complete a transition between any two brain activity states. First, pairwise TE was calculated for all transitions between brain states at the regional level, including persistence. Global and network-level TE were then obtained by summing regional TE across all regions or within each of 9 predefined networks, respectively [53]. TE at the global, network, and regional levels was computed by averaging across all pairwise transitions within each level. All TE measures were z-scored prior to modeling. See Supplementary Materials Section S1.1 for additional details. Global TE findings were consistent with our previous work, showing higher global TE for FH+ females and lower global TE for FH+ males compared to their within-sex FH*−* peers. See Supplementary Section S1.3.

### 2.4 Mediation framework and statistical testing

We then fit multi-group mediation models in lavaan (R) using robust maximum likelihood estimation, with sex as the grouping variable such that all paths were freely estimated in males and females [54]. Models included the three environmental PCs as parallel mediators: EH, FSN, and PMI. In these models, the *a*-paths represented the associations between family history of SUD and each mediator, the *b*-paths represented the associations between each mediator and TE while controlling for family history, and the *c’*-path represented the direct association between family history and TE after accounting for the mediators. Specific indirect effects were calculated as *a×b* for each mediator, and the total indirect effect was defined as the sum of the three specific indirect effects. The total effect (*c*) was defined as the sum of the direct and indirect effects. Because the direct and indirect effects could differ in direction, the total effect reflected their net sum. Models adjusted for age, puberty status, race/ethnicity, mean framewise displacement (FD), and scanner manufacturer, with cluster-robust standard errors to account for non-independence among siblings.

Directly estimated path coefficients (that is, *a*-paths, *b*-paths, and *c’*-paths) were evaluated within each sex using Wald tests, as these correspond to single regression parameters estimated by the model. Sex differences in these directly estimated paths were tested using Satorra–Bentler scaled likelihood ratio tests (LRTs), comparing the unconstrained model with nested models in which the parameter of interest was constrained to equality across sex [55]. By contrast, derived quantities, including specific indirect effects, total indirect effects, total effects, and female–male contrasts in these quantities, were evaluated using Monte Carlo confidence intervals and two-sided *p*-values based on 20,000 Monte Carlo draws generated by sampling parameter vectors from their joint asymptotic multivariate normal approximation using the model’s estimated coefficients and robust variance– covariance matrix. This approach was used because derived effects involve products or sums of coefficients and can therefore deviate from normal sampling distributions [56]. Multiple-comparison correction was applied using the Benjamini–Hochberg procedure within prespecified families of related tests [57]. See Methods Section 5.4.2 and Supplementary Section S3.3.5 for additional details on model specification and multiple-comparison correction.

### 2.5 Environmental mediation of global transition energy

Our parallel mediation model revealed that although all three environmental dimensions were strongly associated with family history of SUD in both sexes (all *p <* 10*^−^*^12^), pronounced sex differences emerged in the indirect pathways linking these environmental factors to global TE (Fig. 2A,B). Specifically, economic hardship (EH) emerged as the primary driver of this sex divergence. In females, family history of SUD exerted a significant, positive indirect effect on global TE via EH (*β* = 0.081, *p*_FDR_ = 0.006). In males, this pathway was negative and nominally significant before correction (*β* = *−*0.049, *p* = 0.049), but did not survive FDR correction (*p*_FDR_ = 0.081). Formal female–male comparison of the EH-specific indirect effect using Monte Carlo-derived contrasts showed that this effect was significantly larger in females than in males (Δ*β* = 0.130, *p*_FDR_ = 0.0018).

**Figure 2.**
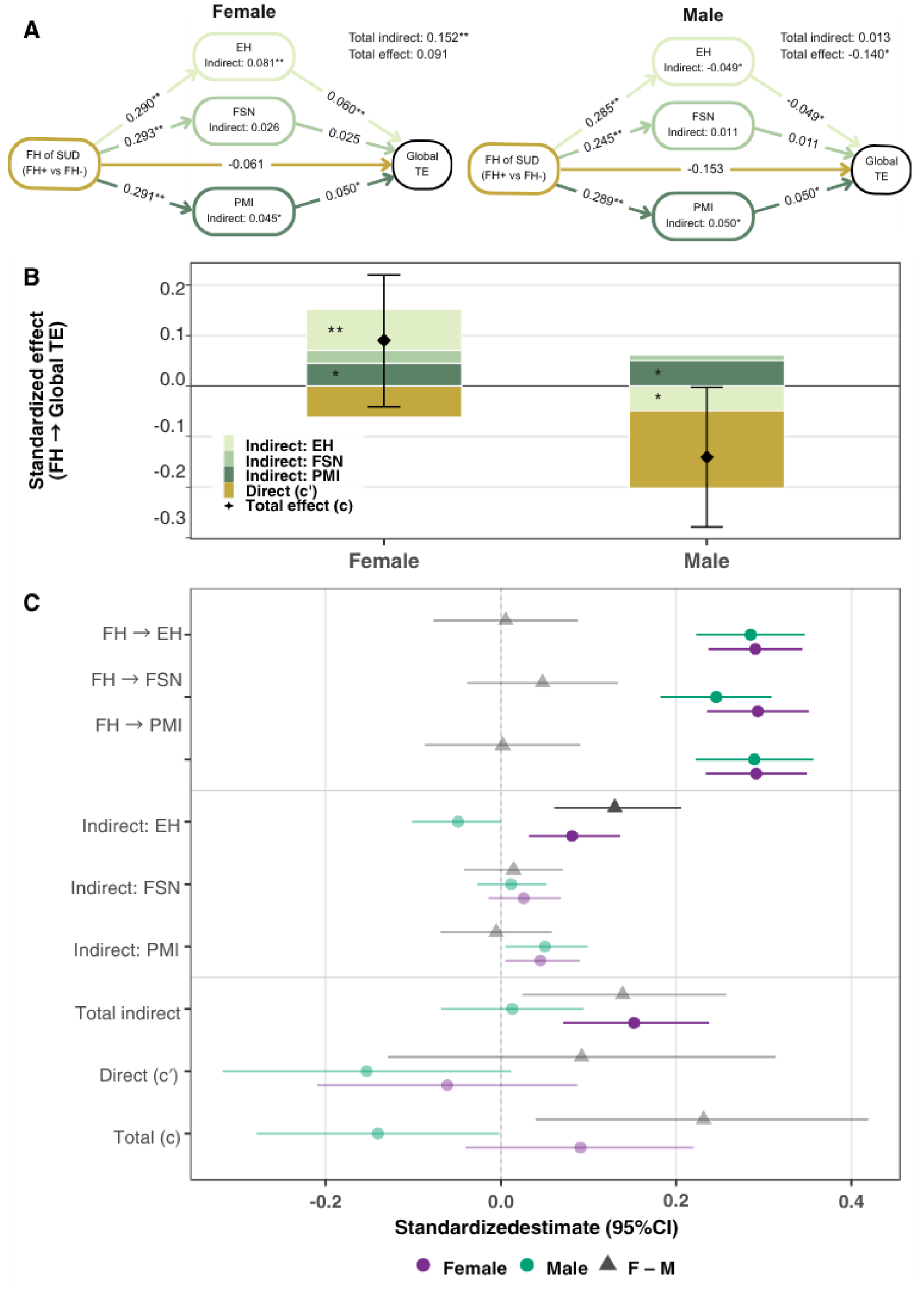
Environmental mediation of family history-related differences in global transition energy differs by sex. **(A)** Sex-stratified parallel mediation model with family history of SUD as the predictor, economic hardship (EH), family–school neglect (FSN), and parental mental illness (PMI) as parallel mediators, and global transition energy (TE) as the outcome. Displayed *a*-paths (FH *→* mediator) are fully standardized coefficients for interpretability; *b*-paths, direct effects, total indirect effects, and total effects are shown on the fitted model scale. Indirect effects were estimated from the fitted mediation model and Monte Carlo draws, and should not be inferred by multiplying the displayed rounded path coefficients. **(B)** Decomposition of the family-history effect on global TE into direct and mediator-specific indirect effects by sex. Black diamonds indicate total effects with 95% Monte Carlo confidence intervals. **(C)** Forest plot of path and effect estimates with 95% confidence intervals for females, males, and female–male contrasts. Purple circles represent females, green circles represent males, and gray triangles represent female–male contrasts. Filled markers indicate effects that survive Benjamini–Hochberg correction (*p*_FDR_ *<* 0.05); unfilled markers indicate effects that do not. TE = transition energy; EH = economic hardship; FSN = family–school neglect; PMI = parental mental illness.

Driven primarily by EH, the total indirect effect on global TE was significant in females (*β* = 0.152, *p*_FDR_ = 0.002) but not in males. The other environmental dimensions played a lesser role. Parental mental illness showed nominally significant positive indirect effects in both sexes, but neither survived multiple-comparison correction, whereas family–school neglect contributed minimally and was not significant.

The direct effects of family history of SUD on global TE were negative but not statistically significant in either sex. The total effect was positive in females (*β* = 0.091, *p* = 0.177) and negative in males (*β* = *−*0.140, *p* = 0.046), but neither survived multiple-comparison correction (*p*_FDR_ = 0.081 for males). Complete within-sex and sex-difference model statistics are provided in Supplementary Tables S4 and SS5.

### 2.6 Environmental mediation of network transition energy

We examined the direct and indirect effects of family history of SUD on TE across nine large-scale brain networks (Yeo 7-networks plus subcortical and cerebellar networks). Again, a clear sex divergence emerged: network-level alterations in females were driven predominantly by environmental mediation, whereas the male pattern showed weaker environmental mediation and more negative direct-effect estimates (Fig. 3A). In females, EH was again the strongest mediator of increases in TE. Specifically, we observed significant, positive EH-mediated indirect effects across the DMN (*β* = 0.090, *p*_FDR_ = 0.010), somatomotor (SOM; *β* = 0.067, *p*_FDR_ = 0.025), visual (VIS; *β* = 0.069, *p*_FDR_ = 0.025), and VAT (*β* = 0.064, *p*_FDR_ = 0.049) network TE. In the VIS and DMN networks, these environmentally mediated pathways were strong enough to drive significant total indirect effects of family history of SUD (*β* = 0.17, *p*_FDR_ *<* 0.001 and *β* = 0.13, *p*_FDR_ = 0.011, respectively).

**Figure 3.**
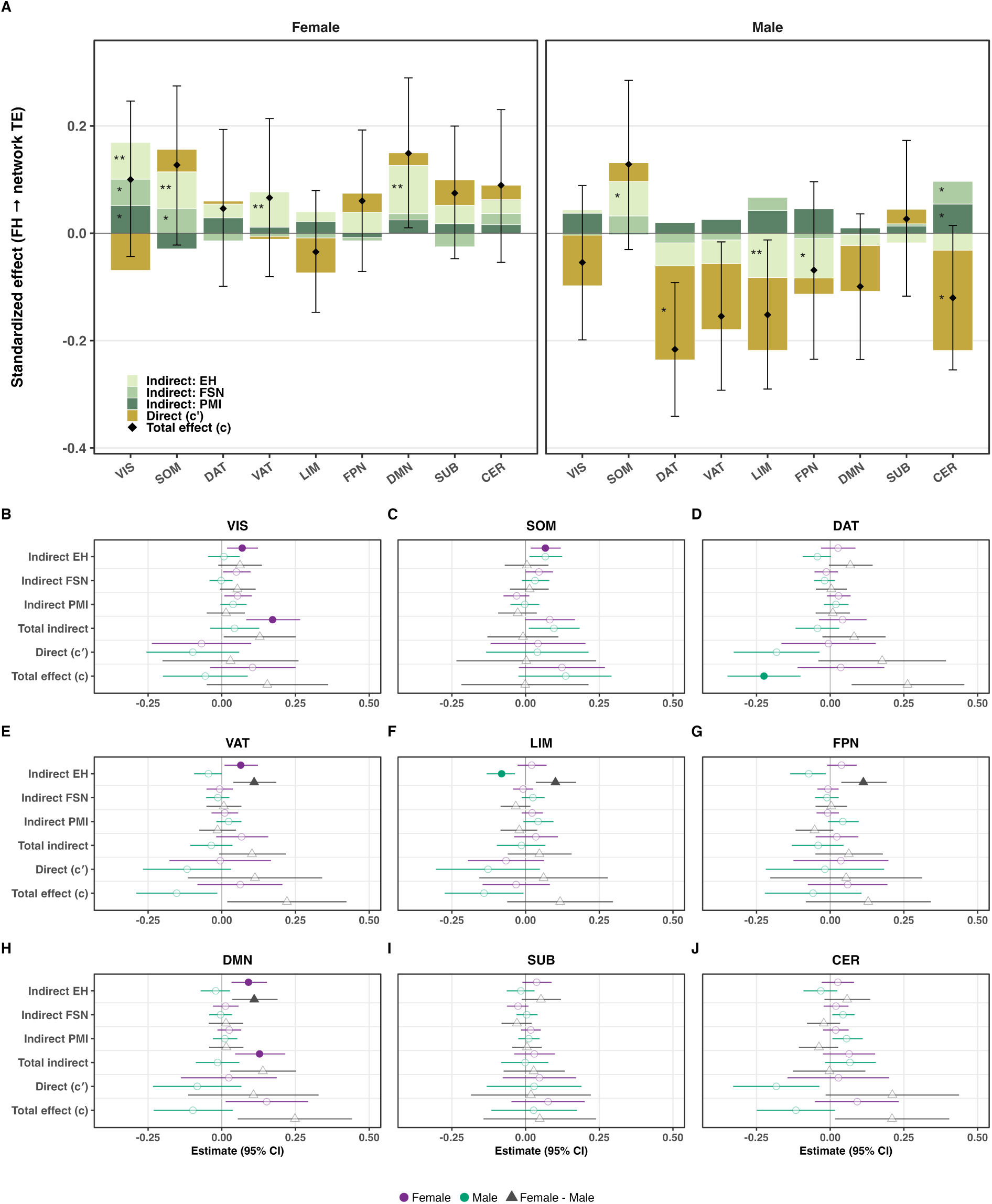
Network-level sex-stratified parallel mediation of family history of SUD on transition energy through environmental principal components. **(A)** Stacked bar plots showing decomposition of the effect of family history on network transition energy (TE) across nine networks into the direct effect (*c^′^*) and specific indirect effects via economic hardship (EH), family–school neglect (FSN), and parental mental illness (PMI), separately for females and males. Black diamonds and error bars indicate total effects with 95% Monte Carlo confidence intervals. *^∗^p <* 0.05; *^∗∗^p*_FDR_ *<* 0.05. **(B–J)** Forest plots for each network showing specific indirect effects, the direct effect (*c^′^*), the total indirect effect, and the total effect (*c*) for females, males, and the female–male contrast, with 95% confidence intervals. Symbols shown at full opacity and filled markers indicate effects that remain significant after Benjamini–Hochberg correction *p*_FDR_ *<* 0.05.

In males, EH-mediated effects were more limited and heterogeneous. The clearest negative EH-mediated indirect effect emerged within the limbic (LIM) network (*β* = *−*0.081, *p*_FDR_ = 0.002), while additional corrected EH-related effects in SOM and frontoparietal (FPN) were smaller and did not alter the broader pattern of weaker and less consistent environmental mediation in males. No direct effects survived multiple-comparison correction in either sex; however, the net effect of these pathways in males culminated in a significant overall reduction in TE within the DAT (total effect *β* = *−*0.22, *p*_FDR_ = 0.003), consistent with our previous work.

Formal sex-difference testing confirmed that this divergence was substantially localized to the EH pathway (Fig. 3B–J). Monte Carlo contrasts showed that the EH-specific indirect effect was significantly greater in females than in males across the FPN (Δ*β* = 0.11, *p*_FDR_ = 0.010), VAT (Δ*β* = 0.11, *p*_FDR_ = 0.010), DMN (Δ*β* = 0.11, *p*_FDR_ = 0.010), and LIM (Δ*β* = 0.10, *p*_FDR_ = 0.010) networks. In each case, the indirect effect was positive in females and negative in males. Sex differences in direct effects did not survive correction in any network; however, sex differences in the *total* effect were FDR-significant in the DAT and DMN networks (both *p*_FDR_ = 0.049), consistent with the localized male DAT total effect and female DMN mediation effect described above. See Supplementary Tables S6 and SS7 for full model statistics and sex-difference results.

Together, these results indicate sex-specific patterns of association between family history of SUD, environmental mediation, and macroscale brain dynamics. In females, positive indirect associations through economic hardship were relatively widespread, whereas in males, environmental indirect associations were more limited and heterogeneous, with a negative direct effect of family history in the dorsal attention network.

### 2.7 Environmental mediation of regional transition energy

To assess more fine-grained regional effects, we extended the parallel mediation framework to the 86 regional TE measures (Fig. 4). These models recapitulated the network-level dissociation: female regional patterns showed coherent, large-scale environmental mediation primarily through EH, whereas male regional patterns showed weaker environmental mediation and more negative direct-effect estimates.

**Figure 4.**
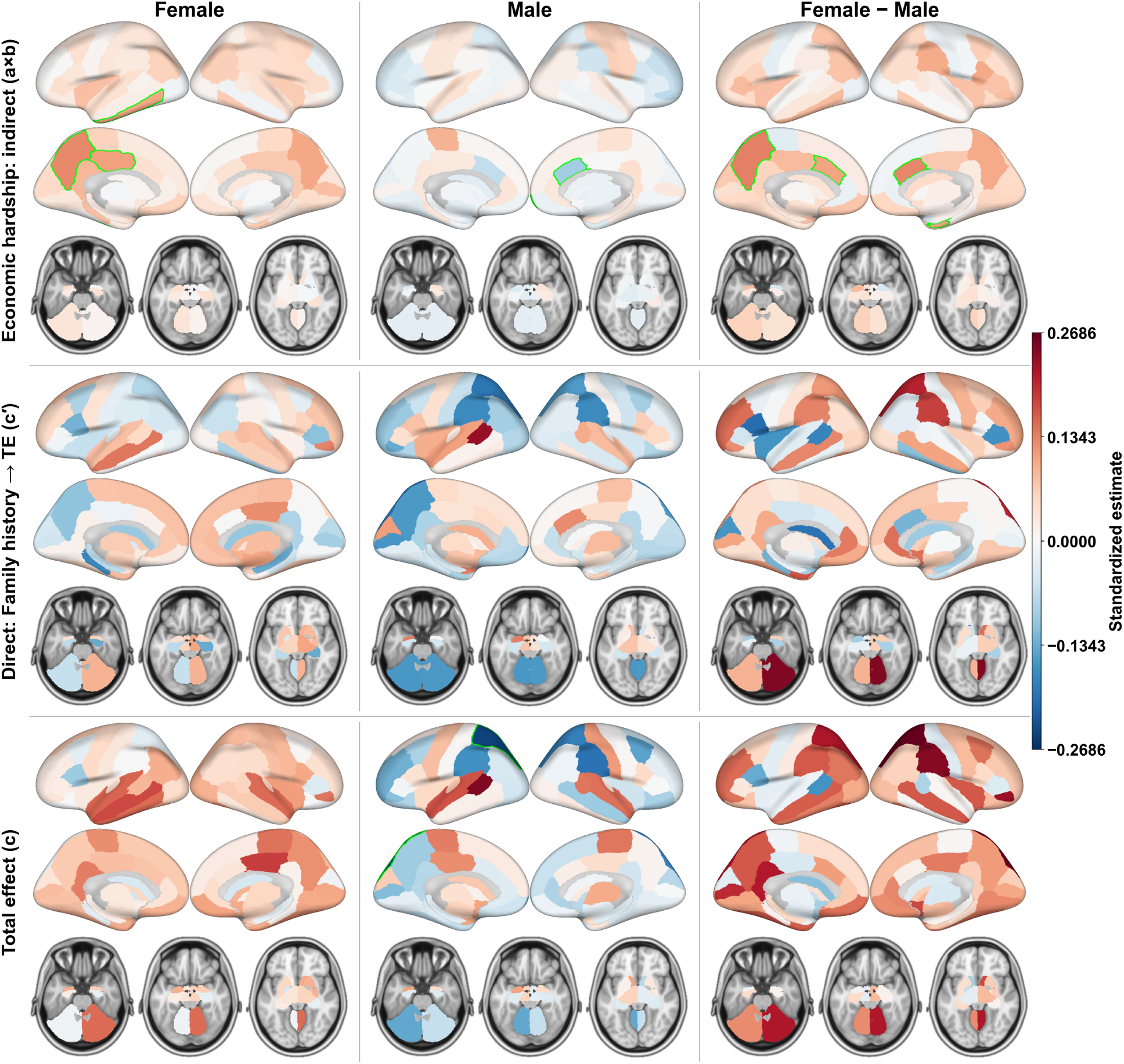
Regional mediation of family history of SUD on regional transition energy. Regional brain maps display effect estimates from sex-stratified parallel mediation models across 86 regions (68 cortical, 16 subcortical, and 2 cerebellar regions). Columns correspond to females (left), males (center), and the female–male sex difference (right). Rows show the economic hardship (EH) indirect effect (top), direct effect (*c^′^*; middle), and total effect (*c*; bottom). Warm colors indicate positive values and cool colors indicate negative values; the color scale is shared across panels. Regions that remain significant after Benjamini–Hochberg correction *p*_FDR_ *<* 0.05 are outlined in green.

In females, total indirect effects were positive in approximately 83% of regions, total effects were positive in approximately 83% of regions, and both were positive in the same region in approximately 66% of regions; EH-mediated specific indirect effects were positive in 91% of regions. After Benjamini–Hochberg adjustment (*p*_FDR_ *<* 0.05) across regions, significant female EH*→*TE *b*-paths localized to five core posterior midline and temporal/occipital regions: left posterior cingulate, bilateral precuneus, left inferior temporal cortex, and right pericalcarine cortex. These regions included posterior midline regions overlapping the DMN, consistent with the network most strongly implicated in increased TE among FH+ females in our previous work. EH-specific indirect effects survived correction in three overlapping regions: left inferior temporal cortex, left posterior cingulate, and left precuneus. Positive total indirect effects were significant in an overlapping but not identical set of five regions: left inferior parietal lobule, left inferior temporal cortex, left parahippocampal gyrus, left precuneus, and right pericalcarine cortex (see Supplementary Table SS8).

In males, direct-effect estimates (*c^′^*) were negative in 57% of regions, although none survived correction. Males’ total indirect effects were spatially heterogeneous and near zero overall; however, localized significant, negative EH-mediated effects emerged in higher-order cognitive hubs, specifically the right caudal anterior cingulate and right frontal pole. Correspondingly, the total effect of family history on regional TE survived correction only in the left superior parietal cortex.

Formal female–male contrasts confirmed this global shift, with positive differences (females *>* males) across approximately 77% of regions for both total and total indirect effects. Monte Carlo sex contrasts of the total indirect effect identified 11 regions at uncorrected significance (*p <* 0.05). After FDR correction, sex differences were significant in four regions for the indirect effect of EH (bilateral caudal anterior cingulate, left precuneus, and right entorhinal cortex), and in eight regions for the EH *b*-path: left amygdala, bilateral caudal anterior cingulate, left lateral orbitofrontal cortex, left precuneus, right entorhinal cortex, right frontal pole, and right pars orbitalis. In each case, EH*→*TE effects were larger in females than in males.

Together, these regional patterns align with the network-level findings, supporting a spatially organized distinction between environment-linked TE effects in females and more negative family-history-related TE estimates in males. Additional regional effect maps and full regional statistics are provided in Supplementary Figures SS15 and SS14, and Tables SS8 and SS9.

### 2.8 Policy-linked projection results

Given that economic hardship mediated FH-related differences in TE, we next asked whether broader state-level policy environments were associated with the FH+ versus FH*−* EH disparity. We then used the previously estimated EH*→*TE mediation paths to project whether policy-associated differences in EH would be expected to translate into smaller or larger FH-related TE gaps (Fig. 5). This analysis was intended to contextualize the mediation findings in relation to public-health policy environments, not to estimate causal effects of policy exposure.

**Figure 5.**
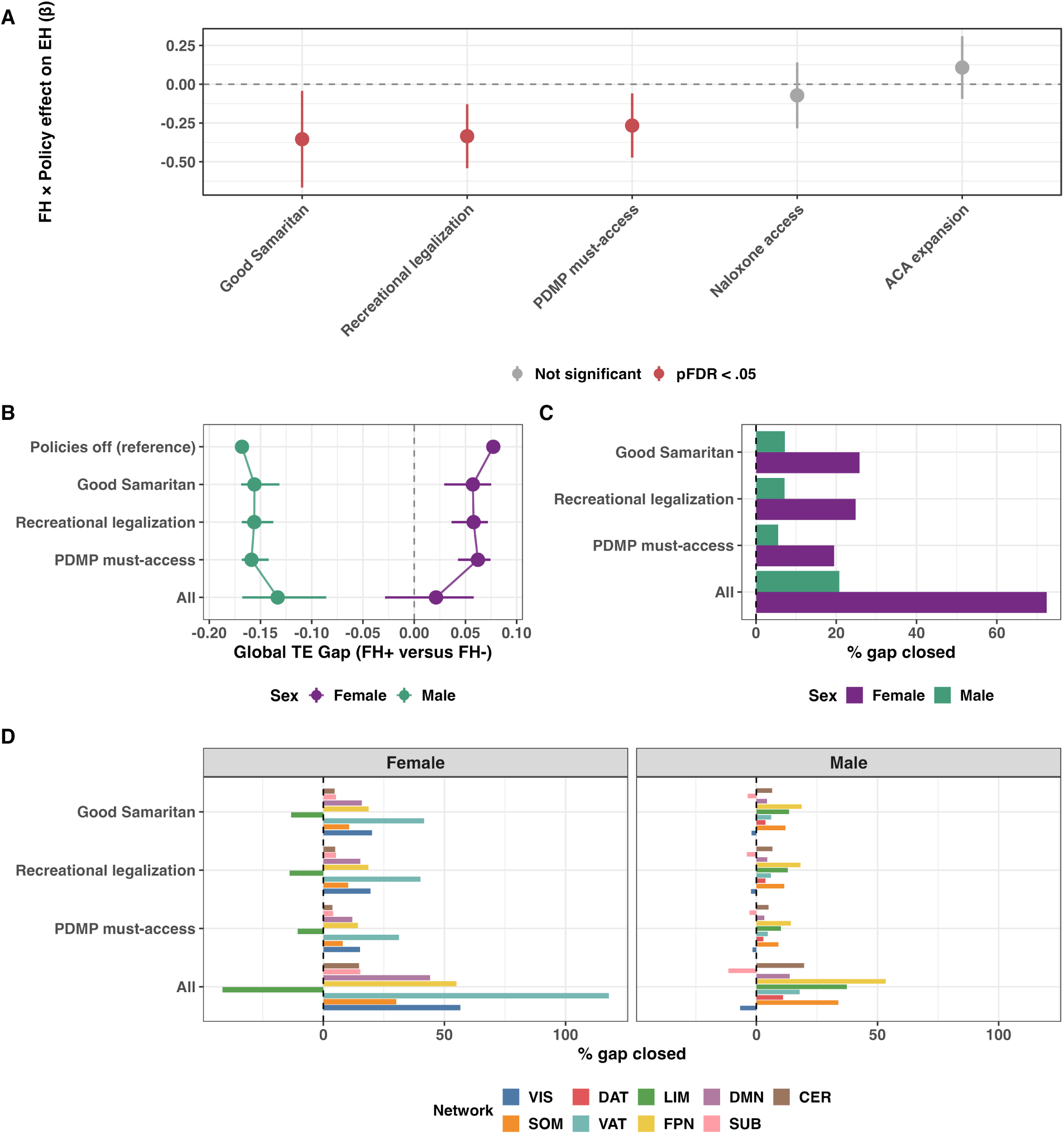
Exploratory policy-linked projection of economic hardship-associated transition energy gap closure. **(A)** Estimated FH*×*policy interaction effects from the joint reduced-domain policy model predicting economic hardship (EH) in the full ABCD cohort with environmental PC scores (*N* = 9,157). Negative coefficients indicate smaller FH+ versus FH*−* EH gaps in states where the policy was present. **(B)** Projected FH+ versus FH*−* global transition energy (TE) gaps under each policy scenario, obtained by propagating policy-associated EH-gap coefficients through the sex-specific EH*→*TE path from the mediation model. **(C)** Percent closure of the observed FH+ versus FH*−* global TE gap, shown separately for females and males. **(D)** Network-level projections from the same EH*→*TE mapping were exploratory and more heterogeneous across outcomes. TE = transition energy; EH = economic hardship; FH = family history of substance use disorder; PDMP = prescription drug monitoring program.

In the full ABCD cohort with environmental PC scores (*N* = 9,157), the joint reduced-domain model identified three of five FH*×*Policy interaction terms as FDR-significant predictors of EH (Fig. 5A). Good Samaritan law (*β*_FH_*_×_*_Policy_ = *−*0.354, *SE* = 0.159, *p*_FDR_ = .043), recreational cannabis legalization (*β* = *−*0.335, *SE* = 0.105, *p*_FDR_ = .007), and PDMP must-access requirements (*β* = *−*0.266, *SE* = 0.105, *p*_FDR_ = .029) were each associated with a smaller FH+ versus FH*−* EH gap in states where the policy was present compared to those without the policy present. ACA Medicaid expansion and naloxone access did not show significant FH*×*Policy interactions.

Propagating the EH-gap estimates through sex-specific EH*→*TE paths projected global TE shifts toward the FH*−* reference level for the three significant policy domains in both sexes (Fig. 5B,C). Individually, the projected global gap closures were largest for Good Samaritan law and recreational cannabis legalization in females, with smaller projected closures in males.

In the additive combined scenario, the FH*×*Policy coefficients for Good Samaritan law, recreational cannabis legalization, and PDMP must-access requirements summed to a combined EH-gap estimate of ΔEH_combined_ = *−*0.956 (*SE* = 0.218, 95% CI [*−*1.383*, −*0.528]). Propagating this combined estimate through the sex-specific EH*→*TE paths projected the global FH TE gap to shift from +0.077 SD to +0.021 SD in females, corresponding to 72.4% gap closure, and from *−*0.168 SD to *−*0.133 SD in males, corresponding to 20.7% gap closure (Fig. 5B,C).

Projected shifts at the network level were heterogeneous in direction and magnitude (Fig. 5D). One network-level projection showed closure exceeding the observed gap (female ventral attention network, 117.8%). Several regional projections also showed projected overshoot, whereas other network and regional outcomes showed negligible or reversed projected shifts. These heterogeneous patterns support interpreting the policy-linked projections as exploratory, associational, and pathway-specific rather than as causal estimates of policy effects.

### 2.9 Sensitivity Analyses

To assess whether pubertal status moderated the observed effects of family history of SUD in males and females, we conducted an exploratory parallel mediation analysis in which binary pubertal status (started versus not started) moderated all paths in the model. This analysis suggested that the positive total effect of family history on global TE in females remained directionally stable but somewhat attenuated across pubertal status (pre-pubertal *β* = 0.17; pubertal *β* = 0.06; neither significant). In males, the nominally significant negative total effect observed in pre-pubertal youth (*β* = *−*0.17, *p* = 0.035) was attenuated and no longer significant among pubertal youth (*β* = *−*0.05, *p* = 0.70), as a shift toward positive indirect environmental effects offset more of the negative direct-effect estimate than in pre-pubertal males (see Supplementary Section S3.4). Given the reduced sample sizes after stratification, these findings should be interpreted cautiously.

Additionally, we repeated the mediation analysis in a complete-case cohort restricted to subjects with all variables included in the PCA and therefore requiring no imputation (*N* = 1,250; see Supplementary Section S3.5). Results were largely consistent with the main analysis, and differences likely reflect reduced power due to the smaller sample size in each subgroup. We also repeated the PCA in two restricted cohorts: (1) only subjects included in the mediation analysis (Supplementary Section S2.6); and (2) the full ABCD baseline cohort restricted to subjects with complete PCA data (Supplementary Section S2.7). The resulting PCA components were highly correlated with those identified in the main analysis in both restricted cohorts (mediation cohort, *r ≥* 0.95; complete-case cohort, *r ≥* 0.98).

## 3 Discussion

Here, we show that a family history of SUD is associated with sex-divergent alterations in brain dynamics during early adolescence, and that these alterations arise through different pathways in females and males. FH+ youth of both sexes experienced greater economic hardship, family–school neglect, and parental mental illness than FH*−* youth. Thus, family history in this cohort did not function as a purely familial or genetic marker; rather, it indexed a more adverse early life context. However, the neurobiological consequences of this adversity differed markedly by sex. In females, family history-related differences in transition energies were driven primarily by positive indirect environmental effects, especially through economic hardship, with smaller concordant contributions from parental mental illness. This environmentally mediated pattern was evident globally, across several large-scale networks—most prominently the DMN, VIS, VAT and SOM networks—and regionally, where indirect effects were widespread and predominantly positive. In contrast, the total effect of family history in males was dominated by a negative direct effect and showed more limited and localized environmental mediation, including a negative EH-mediated indirect effect in the LIM network. Overall, these findings indicate that familial SUD risk is associated with different neurodevelopmental pathways in the two sexes: widespread environmentally mediated increases in TE in females, and more weakly mediated, largely direct reductions in TE in males.

Specifically, having an SUD family history was associated with a positive indirect effect on global TE through economic hardship for females, with a smaller positive contribution from parental mental illness factors. In contrast, the indirect environmental pathways were weaker in males, and the overall pattern was characterized primarily by a negative direct SUD family history effect not explained by the environmental dimensions measured here. Therefore, the observed sex differences were not simply that females showed higher TE and males showed lower TE, as found in our previous work as well [49]. Rather, while SUD family history was associated with greater adversity in both sexes, that adversity translated into higher TE in females only, whereas males exhibited an overall pattern of a strong negative direct effect of family history. Parental mental illness showed a smaller but directionally similar positive contribution, reinforcing the overall effect of increased TE in females and somewhat offsetting the decreased TE in males.

The network- and regional-level findings support this interpretation. In females, positive indirect effects were spatially widespread and emerged across several cortical systems, particularly the DMN, VIS, VAT and SOM networks. At the regional level, the female indirect effect of economic hardship was broad and predominantly positive, extending across cortex, especially posterior temporal and midline regions. In males, negative family-history-related TE estimates were more prominent. Environmental mediation was comparatively limited, network-specific, and mixed in direction, with negative EH-mediated effects most evident in LIM and FPN networks and a smaller positive EH-mediated effect in SOM. Together, these results suggest that the female profile reflects distributed environmental shaping of brain dynamics, whereas the male profile reflects liability not fully captured by (mostly) environmental dimensions modeled here.

Within the framework of network control theory, TE reflects the energetic cost required to move the brain between states under the constraints of structural connectivity [31]. Higher TE may therefore reflect more constrained or “sticky” dynamics, whereas lower TE may reflect more disinhibited dynamics. This interpretation remains inferential, but it provides a useful way to situate the present findings. From this perspective, the female pattern of higher TE, driven largely by economic hardship, is broadly consistent with a more rigid or internalizing-like mode of vulnerability, while the male pattern of lower TE is more consistent with a more disinhibited or externalizing-like mode of vulnerability [58–63]. This interpretation aligns with broader evidence that pathways to SUD more often involve internalizing symptoms in females and externalizing symptoms in males [64–67].

### 3.1 Females: Higher transition energy is driven primarily by economic hardship

In females, neural alterations associated with a family history of SUD are fundamentally reshaped by environmental context. A nominally negative direct effect of family history is offset by a positive and widespread indirect effect driven primarily by economic hardship. Financial scarcity may contribute to the embedding of familial risk in the brain through increased TE across the DMN, VIS, VAT, and SOM networks. Notably, alterations in core DMN hubs such as the posterior cingulate cortex and precuneus, as observed here, have been repeatedly implicated in self-referential processing, internally oriented thought, and internalizing psychopathology [68–71]. The concurrent involvement of lower-order networks (VIS, VAT, and SOM) suggests that this effect is not limited to higher-order, internally focused processing, but also extends to systems involved in sensory processing, bodily state, and attentional orienting [53, 72].

This widespread cross-network pattern is consistent with models of biological embedding, in which chronic financial instability may predispose the brain to remain in a stress-responsive state, with higher TE perhaps reflecting greater difficulty shifting out of that state [73]. This interpretation is also consistent with evidence that females may be especially sensitive to stress-related pathways to SUD [15]. Together, these results suggest that economic hardship may be particularly important for neurobiological risk among females with familial liability, potentially amplifying the effects of environmental adversity on brain dynamics in FH+ females relative to FH*−* females.

One plausible explanation for greater female environmental sensitivity is developmental timing. Females generally enter puberty earlier than males, and early adversity may accelerate pubertal maturation more strongly in girls [39–41]. In our sample, females were more advanced in pubertal development than males, and although FH+ youth of both sexes were more advanced than FH*−* youth, this difference was more pronounced in females. This raises the possibility that sex differences in the mediation of TE may be partly related to differences in hormonal and developmental timing [43, 74]. While puberty initiation was included as a covariate in all primary models, exploratory moderation analyses here revealed sex-specific trends warranting further investigation. Specifically, the total effect of family history on global TE in females remained positive but was not significant within either pubertal stratum, whereas the negative total effect in pre-pubertal males appeared to attenuate in pubertal males as it was offset by positive indirect environmental effects (Supplementary Section S3.4). These patterns must be interpreted cautiously, however, as the analyses were underpowered by a limited number of pubertal FH+ males (*n* = 60). Ultimately, these results suggest pubertal status may shape the expression of family history-related effects on TE, though adequately powered longitudinal designs are required to confirm this.

Another potential mechanism is greater female sensitivity to earlier prenatal factors. Prior work has shown that prenatal maternal cortisol is associated with female-specific differences in amygdala development and later affective symptomatology [75]. Although prenatal maternal stress was not directly examined here, future work should test whether female-specific effects of prenatal factors contribute to the shaping of brain dynamics.

### 3.2 Males: Lower transition energy is not driven by indirect environmental effects

The male pattern was strikingly different. Although FH+ males were similarly exposed to elevated environmental risk, these factors accounted for much less of the effect of family history of SUD on TE than they did in females. Instead, lower TE in males was driven primarily by negative family-history-related estimates, most clearly reflected in the DAT network and in the left superior parietal lobe. Negative indirect effects of economic hardship were also observed in the LIM and FPN networks, further contributing to this lower-TE profile.

The direct effect of family history indexes liability not captured by the environmental mediators and may reflect genetic influences, prenatal exposures, unmeasured features of the environment, or gene–environment interactions. Some evidence suggests that heritable influences on SUD may be stronger in males than in females, although the literature is mixed [36, 46, 47]. For example, a large study of Swedish twins and sibling pairs found that genetic factors accounted for a substantially larger proportion of the variance in alcohol use disorder in males than in females, despite a high overall genetic correlation between sexes [76]. Relatedly, a recent ABCD study reported that schizophrenia polygenic risk scores (PRS), which share considerable genetic liability with SUD [77], were more strongly associated with cognitive function in males than in females, although they were not associated with measured brain metrics [78]. We therefore cautiously interpret the unmediated direct effect as suggestive of stronger inherited liability in males, while recognizing that it may also reflect environmental or developmental influences not represented by the included mediators. Future models should incorporate PRS and related measures of inherited risk to further disentangle these possibilities.

Altered dynamics in the DAT and VAT networks have been associated with attentional deficits, impulsive behavior, and externalizing disorders [79–81]. As such, the lower energetic demands associated with SUD family history in these networks may reflect a neural risk profile for externalizing pathways to SUD. The more limited environmentally-mediated effects in males were concentrated in LIM and FPN networks, a pattern broadly consistent with prior work showing that socioeconomic adversity is linked to variation in frontolimbic systems in youth [82]. At the regional level, economic-hardship mediation was localized to the right caudal anterior cingulate and right frontal pole, regions central to salience monitoring, conflict processing, and executive function that have also shown SES-related alterations in activity [83]. Adversity-related alterations of anterior cingulate structure have also been reported in males, supporting the interpretation that when environmental mediation is present in FH+ males, it may preferentially manifest in cingulo-frontal systems involved in control and regulation rather than in the broader environmentally mediated pattern observed in females [84].

### 3.3 Convergent and shared pathways in males and females

Despite these robust sex differences, the effects of family history were not entirely distinct across males and females. In both sexes, family history was associated with greater environmental adversity, and the direct effect of family history on global TE was negative, although neither the direct nor the total effects were statistically significant after multiple-comparison correction. In females, positive environmental mediation contributed to a positive total-effect estimate, whereas the total-effect estimate remained negative in males. These opposing patterns suggest that environmental pathways may contribute differently to the association between family history and brain dynamics across sexes, although the global total effects should be interpreted cautiously.

There were also common patterns of sensitivity to the environmental context associated with family history. Positive EH-mediated effects in the SOM network were observed in both sexes and survived FDR correction in both females and males. This is consistent with prior findings that the somatomotor network is sensitive to socioeconomic and neighborhood adversity [85–87]. This convergence suggests that some network-level associations with environmental adversity may be shared even when the broader pattern differs by sex.

Although weaker and less consistent than economic hardship, the PMI dimension showed concordant positive indirect effects in both females and males, though these associations were not robust after multiple-comparison correction. This directional consistency may reflect shared aspects of familial risk, but the lack of statistical significance precludes strong conclusions about a common pathway. In females, the direction of the PMI indirect effect is consistent with the positive economically mediated pathway, whereas in males it occurs alongside a negative total-effect estimate. The broader pattern is consistent with evidence that the intergenerational transmission of psychopathology is often non-specific and transdiagnostic, while the expression of these outcomes can show sex-specific tendencies [88, 89]. The positive direction of the PMI effect, together with strong PMI loadings reflecting parental affective and depressive histories, may also be relevant to evidence that girls more often express psychopathology risk through internalizing symptoms during adolescence [13, 50].

Overall, although some effects were directionally similar in females and males, the sexes differed in the magnitude of associations, the relative contribution of measured environmental pathways, and the networks in which these effects were most strongly expressed. Longitudinal analyses are needed to better understand how sex, family history of SUD and other mental illnesses, and environmental context jointly relate to later behavioral patterns and SUD risk.

### 3.4 Implications for prevention and policy

These findings have implications for both mechanism and intervention. The main mechanistic implication is that familial SUD risk appears to contain at least two separable components: a direct liability signal and an adversity-linked environmental signal. In females, the environmental signal, especially economic hardship, appears to be more prominent. In males, the direct signal remains more prominent. This helps clarify why the same family-history exposure can lead to divergent neural signatures across sexes. Future work should aim to better delineate the relative contributions of genetic liability, prenatal exposures, and additional environmental factors to neural risk in this population.

Our exploratory policy-linked projection analysis extends this interpretation by asking whether broader state-level policy environments associated with smaller FH+ versus FH*−* EH disparities would be expected to translate into smaller FH-related TE gaps through the EH pathway. In this analysis, Good Samaritan laws, recreational cannabis legalization, and PDMP must-access requirements were each associated with smaller FH-related EH gaps. When these EH-gap estimates were propagated through the sex-specific EH*→*TE paths, they projected partial closure of the global FH TE gap in both sexes, with substantially larger projected closure in females. Thus, even though males showed a stronger direct family-history pathway, the policy-linked analysis suggests that modifiable environmental pathways may remain relevant in both sexes, even if they account for a smaller share of the observed TE pattern in males.

These projections should be interpreted cautiously. They do not show that any specific policy caused changes in EH or TE, nor are policies themselves neuroprotective. Rather, the significant policy indicators may partly index broader state-level policy contexts, harm-reduction infrastructures, or social environments that co-occur with reduced hardship among FH+ youth. This interpretation is consistent with emerging evidence that macrostructural policy environments, including state-level safety-net and anti-poverty policies, can moderate associations between socioeconomic disadvantage, brain development, and child mental health [90]. It is also consistent with causal evidence from poverty-reduction trials showing that direct economic support can alter early neurodevelopmental measures [91], and with broader models linking poverty, stress physiology, and child brain development [92].

The present results suggest that early adolescent girls with familial SUD risk may be especially sensitive to socioeconomic and policy-relevant environmental conditions during this developmental window. Such sensitivity may reflect differences in pubertal timing, stress responsivity, and the developmental organization of affective and self-referential neural systems [15, 93, 94]. Importantly, greater environmental sensitivity should not be interpreted as greater vulnerability alone. It may also indicate greater potential responsiveness to upstream interventions that reduce economic hardship, family stress, and structural barriers to care.

Together, these findings suggest that preventing the inter-generational transmission of SUD risk will likely require more than individual-level screening or clinical intervention. Family-history risk is embedded within broader social and economic contexts, and policy environments may shape how that risk is expressed in the developing brain. Although the present analysis is exploratory and associational, it highlights the possibility that structural interventions targeting economic hardship and harm-reduction infrastructure could have neurodevelopmental relevance, particularly for females, while still benefiting males through smaller but meaningful environment-linked pathways.

### 3.5 Limitations

Several limitations must be considered. First, the cross-sectional nature of the analysis restricts causal interpretation. Although the mediation models are useful for decomposing associations into direct and indirect paths, the results should be interpreted as a mechanistic framework rather than proof of mediation in a causal sense. Second, principal component analysis was useful for reducing a large set of environmental variables into interpretable dimensions, but this came at the cost of specificity. Economic hardship, family–school neglect, and parental mental illness are broad constructs, and each likely contains multiple distinct mechanisms that may not act uniformly. For instance, parental mental illnesses, including depression, mania, and affective disorders, could have distinct or even opposing associations with TE. More targeted modeling of specific exposures will therefore be important in follow-up work.

Third, the family history variable used here is heterogeneous. It combines inherited liability with correlated family and environmental exposures, does not distinguish parental from grandparental transmission, and does not separate substance-specific pathways. This relationship is further complicated by the possibility that some variables typically treated as environmental may themselves be partly genetically influenced through gene–environment correlation, whereby inherited traits can shape the environments individuals experience [95]. We did not directly incorporate PRS or explicit measures of gene–environment association. Future work should integrate genetic data more directly while remaining cautious about interpretation.

Fourth, although we adjusted for puberty, our measure was coarse, indexing whether puberty had started rather than pubertal timing or tempo. Pubertal development is likely important, particularly given evidence that socioeconomic conditions, puberty, and psychopathology interact in sex-specific ways [37, 43].

Fifth, the policy-linked projection analysis has additional limitations. Policy exposure was assigned at the ABCD site-state level rather than by confirmed residential state, and the policy indicators may capture broader state-level social, political, and public-health environments rather than isolated effects of specific laws. The combined projection also assumes additivity and independence among policy estimates. Therefore, these results should be interpreted as exploratory evidence that policy-relevant environments may shape the EH pathway, not as causal estimates of policy effects.

Finally, the interpretation of TE itself remains conceptual. The mapping of higher TE onto more constrained dynamics and lower TE onto disinhibition of dynamics is useful and theoretically grounded, but TE is not a direct measure of internalizing or externalizing psychopathology. Longitudinal work will be necessary to determine whether the sex-specific TE profiles identified here prospectively predict internalizing symptoms, externalizing symptoms, and eventual substance use initiation in distinct ways.

## 4 Conclusion

Childhood mental illness, substance use, and economic insecurity are rising in the United States, underscoring the urgency of identifying early mechanisms of SUD risk before disorder onset. Here, we show that even before substance exposure, children with a family history of substance use disorder exhibit widespread and sex-divergent alterations in the brain’s energetic landscape. Family history was associated with distinct patterns of transition energy across females and males, most prominently in the default mode network in females and the dorsal attention network in males. Importantly, these patterns implicated different environmental pathways. In females, family history-related differences in transition energy were associated with economic hardship and, to a lesser extent, parental mental illness. In males, environmental pathways were more limited and heterogeneous, suggesting that other factors may also contribute to family history-related differences in brain dynamics.

The exploratory policy-linked projection analysis further suggests that state-level policy environments associated with smaller FH-related EH disparities may partially attenuate FH-related TE gaps through the EH pathway. These projected effects were present in both sexes but were larger in females, consistent with the stronger environmentally mediated TE pathway observed in girls during early adolescence. Although these projections are associational, they underscore the potential importance of upstream, structural interventions for modifying the developmental context in which familial SUD risk is biologically embedded.

Together, these findings indicate that familial risk for SUD is associated with distinct sex-specific patterns of brain dynamics and environmental mediation long before substance initiation begins. They also point to children’s financial, social, family, and policy environments as potentially modifiable targets for reducing the intergenerational transmission of risk.

## 5 Methods

### 5.1 Study design and participants

#### 5.1.1 Sample Information

The Adolescent Brain Cognitive Development (ABCD) study is longitudinally tracking the brain development and health of a nationally representative sample of children aged 9-10 years old (at the time of enrollment) from 21 centers across the United States (https://abcdstudy.org; [48]). All parents or legal guardians provided written informed consent prior to participation in the study, and children provided verbal assent. Participants and their families received financial compensation for their time. Research protocols were approved by the institutional review board of the University of California, San Diego (no. 160091), and the institutional review boards of the 21 data collection sites [96]. The current study utilized neuroimaging data from the 2.0.1 release and non-imaging instruments from the baseline assessment updated to ABCD Data Release 5.1.

#### 5.1.2 Exclusion criteria

For the principal components analysis (PCA), we used the full baseline ABCD cohort (N = 11,868). Environmental variables were imputed within site, which allowed inclusion of participants with partial missingness in these measures [97, 98]. As a result, the subsequent NCT and mediation analyses were able to retain participants with missing income or parental education data, as in our previous work [49], yielding a slightly larger analytic sample than in our prior study (N = 1,835 versus N = 1,611).

For the mediation analysis, we sought to maximize sample size by including all participants with available imaging data (that is, post-quality-control rs-fMRI time series, mean framewise displacement, and scanner manufacturer), family history of SUD classification meeting criteria for FH+ or FH*−*, and complete covariate data (age, sex, race/ethnicity, family ID, and pubertal status). From the original ABCD cohort (N = 11,868), we excluded youth who: **(i)** did not pass strict MRI quality control and/or exclusion criteria established previously by [52, 99] (N = 9,506); **(ii)** were scanned on Philips scanners, in accordance with ABCD consortium guidelines (N = 2); **(iii)** did not meet criteria for either FH+ or FH*−* (Section 5.1.3; (N = 428)); **(iv)** were adopted (N = 32); **(v)** had prior substance use reported by either the child or parent (Section 5.1.4; (N = 48)); **(vi)** had missing pubertal status data (Section 5.1.5; (N = 15)); or **(vii)** were missing family ID information (N = 2). The number of Asian participants was limited and were thus recoded to the Other race/ethnicity category. The final analytic cohort included (N = 1,835) participants.

#### 5.1.3 Family History of Substance Use Disorder

We used the baseline Family History Module Screener (FHAM-S) [100, 101], in which parents reported substance use and psychopathology among first- and second-degree biological relatives. In the FHAM-S, drug or alcohol problems may include marital, work, school, legal (e.g., DUI), health, rehabilitation, heavy use, or social issues. Following prior work [20, 50, 102], participants were classified as FH+ if they had *≥* 1 parent or *≥* 2 grandparents with a history of SUD; FH*−* if no parents or grandparents had SUD. Participants with just one grandparent with SUD, presumed to have minimal genetic load [20], were excluded from analyses. Additionally, participants who were adopted were excluded due to limited environmental load of familial risk and/or inaccurate family history data (N = 32).

#### 5.1.4 Childhood Exposure to Substances

To isolate the effects of family history of SUD from substance use itself, we excluded youth who had initiated substance use according to parents or children themselves [103]. Participants were excluded if they self-reported lifetime use of more than a sip of alcohol, more than a puff of a cigarette/e-cigarettes, or any use of nicotine products, cannabis products, synthetic cannabinoids, cocaine, cathinones, methamphetamine, ecstasy/MDMA, ketamine, gamma-hydroxybutyrate, heroin, psilocybin, salvia, other hallucinogens, anabolic steroids, inhalants, or prescription misuse of stimulants, sedatives, opioid pain relievers or over-the-counter cough/cold medicine (N = 34). Parents were also asked about their child’s substance use including alcohol (consumed three or more drinks a day, consumed two drinks in the last 12 months) or used drugs (cocaine, marijuana, solvents, stimulants, tobacco, opioids, hallucinogens, sedatives, or other). If parents endorsed any of these, children were excluded from analyses (N = 14).

#### 5.1.5 Pubertal Status

We used parent-reported Pubertal Development Scale (PDS) summary scores from the baseline visit [104], as youth tend to overestimate their development at younger ages [105]. When parent reports were missing (N = 30), child-reported scores were used. PDS questions are sex-specific and were summarized into a five-level categorical stage: (1) pre-pubertal, (2) early pubertal, (3) mid-pubertal, (4) late pubertal, and (5) post-pubertal. Due to small sample sizes in higher stages (e.g., only one male in stage *>*3), we collapsed stages into a binary variable of: started (stage *≥* 2) or not started (stage 1). Females generally had more advanced pubertal stages than males, and FH+ youth of both sexes showed more advanced development than their respective FH*−* counterparts. The proportion of youth who had initiated puberty also differed across the four sex-by-family-history groups (Pearson *χ*^2^, *p <* 0.001. See Table 1).

### 5.2 Neuroimaging Data

#### 5.2.1 Parcellation

In our main results, we present analyses in which the rsfMRI timeseries and group-average structural connectome (SC) were parcellated using an 86-region atlas derived from FreeSurfer (FS86) combining the 68 region Desikan-Killiany (DK68) gyral atlas (34 cortical regions per hemisphere) with 16 subcortical structures (8 per hemisphere, excluding brainstem) and 2 cerebellar structures (1 per hemisphere) to render a whole brain anatomically-defined parcellation for each subject [106, 107]. Each cortical region was assigned to one of seven networks of the functionally-defined Yeo 7-network parcellation [53]. Subcortical regions were assigned to a subcortical network and cerebellar regions to a cerebellar network.

#### 5.2.2 Resting-state Functional Magnetic Resonance Imaging (rsfMRI)

We analyzed baseline rsfMRI data from the ABCD study using the same pre-processing and quality control pipeline described in our previous work [49], based on minimally processed scans as processed and described in [52, 99]. Briefly, Philips scanner data were excluded due to known post-processing issues. Pre-processing included removal of initial frames, alignment to T1 images via boundary-based registration (BBR), and censoring of volumes with framewise displacement *>* 0.3 mm or DVARS *>* 50, along with adjacent volumes. Runs were excluded if more than 50% of volumes were censored or if BBR cost exceeded 0.6.

Nuisance covariates, including global signal, motion parameters, ventricular and white matter signals, and their derivatives, were regressed out using non-censored volumes. Data were bandpass filtered (0.009 *≤ f ≤* 0.08 Hz), mapped to FreeSurfer fsaverage6 space, and smoothed using a 6-mm full-width half-maximum kernel. We then removed censored volumes and normalized BOLD time series by mean gray matter signal before filtering. After censoring, an average of 1158.3 *±* 289.18 rsfMRI frames remained per scan.

#### 5.2.3 Structural Connectomes

Diffusion MRI (dMRI) data from 149 individuals (58.39% female, aged 9-11) ABCD baseline assessment [48] were processed using the same pipeline as in our previous work [49]. Briefly, deterministic tractography with SIFT2 weighting and regional volume normalization was used to construct structural connectomes in the FS86 parcellation. The group-average SC matrix was symmetric with zeros along the diagonal. A group-average structural connectivity (SC) matrix was computed across the 149 participants. The use of a group-average SC is supported by prior network control theory work [108] and evidence suggesting that familial risk for substance use disorder is primarily reflected in functional, rather than structural, differences [21, 109]. Furthermore, in our previous work, we found our results were highly correlated when using group-average and individual SC matrices [49].

### 5.3 Network Control Theory Analyses

#### 5.3.1 Extraction of brain states

As in our previous work [49], all participants’ fMRI time series were concatenated and *k*-means clustering was applied to identify recurring patterns of brain activity (i.e., brain states). Pearson correlation was used as the distance metric, and clustering was repeated 10 times with random initializations, selecting the solution with the best separation. To assess stability, we repeated the clustering procedure 10 independent times and computed the adjusted mutual information (AMI) across resulting partitions. The partition with the highest total AMI relative to all others was selected for downstream analyses. Across runs, AMI values were consistently high (*>* 0.99), indicating highly stable clustering (see Supplementary Figure S6). The optimal number of clusters was set to *k* = 4, as established previously using the elbow criterion, where additional clusters explained *<* 1% incremental variance. Cluster centroids were assigned to one of nine *a priori* canonical resting-state networks (RSNs), with the seven cortical networks defined according to Yeo et al. [53]. Assignment to cortical networks was based on cosine similarity with binary RSN maps for interpretability, while subcortical and cerebellar regions were assigned to their respective networks. Because the mean BOLD signal was removed during band-pass filtering, positive centroid values reflect above-mean (high-amplitude) activity, whereas negative values reflect below-mean (low-amplitude) activity. Subject-specific centroids were computed by averaging all time points assigned to each cluster across an individual’s rsfMRI scans. No participant had an empty or sparsely represented state assignment, indicating that each subject-specific centroid was estimated from multiple time points rather than from rare or idiosyncratic state occurrences.

#### 5.3.2 Transition Energy Calculations

Transition energy (TE) was computed following the procedure described in our previous work [49]; we summarize the key steps here. Briefly, we modeled brain dynamics using a linear time-invariant system:

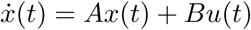

where *A* is a representative (group-average) structural connectivity matrix with dimensions *N*_regions_ *× N*_regions_ (86 × 86), estimated using deterministic tractography from a subset of ABCD participants (*N* = 149), as described above. The matrix *A* was normalized to ensure system stability, as described previously. We used the time horizon (*T* = 1.501), determined in our previous work in this cohort [49], which maximized the inverse relationship between transition energy and transition probability [110, 111]. Using this framework, we computed the minimum control energy required to transition between brain states defined via *k*-means clustering (Section 3.3.1). Pairwise regional TE was calculated for all state transitions, including persistence, yielding a *k* × *k* matrix per region (*k* = 4).

Global and network-level TE were obtained by summing regional TE across all regions or within predefined networks [53], respectively. Mean TE at the global, network, and regional levels was computed by averaging across all pairwise transitions. All TE measures were z-scored prior to modeling.

### 5.4 Environmental Mediation Analysis

#### 5.4.1 Principal component analysis: defining environmental mediators

To reduce the dimensionality of potentially correlated environmental exposures, we performed principal component analysis (PCA) on 43 household-, family-, prenatal-, school-, and neighborhood-level variables derived from the baseline visit of the ABCD Release 5.1 sample (*N* = 11,868 participants). The PCA input matrix comprised 11,868 participants and 43 environmental indicators (11,868 × 43) before centering, scaling, and component extraction. Variables were selected *a priori* to capture exogenous environmental factors relevant to substance use disorder (SUD) risk, excluding child diagnostic and behavioral measures.

The final PCA input set included indicators of socioeconomic position, material hardship, access to care, household composition, parental monitoring, household employment and caregiver structure, school environment, family climate, trauma exposure, neighborhood safety and crime, built environment (e.g., area deprivation index, population density, proximity to major roads), screen time, sleep duration, religious importance, and prenatal exposures. Several variables were harmonized into composite measures to improve interpretability and reduce redundancy, including parent–child communication, parental warmth, family climate, neighborhood safety, material hardship, and access-to-care barriers. Trauma exposures were collapsed into domain-level indicators (non-interpersonal trauma, violence exposure, and bereavement) to reduce sparsity.

Prior to PCA, the distribution of all variables were visually inspected and right-skewed variables (neighborhood violent crime, population density, proximity to major roads, and weekly screen time) were log-transformed using a log(1+x) transformation [112]. No variable had more than 15% missing data, and the prevalence of binary indicators ranged from 6% to approximately 74%. Missingness was handled using a site-aware imputation procedure, with median imputation for continuous variables and mode imputation for binary variables performed within site where possible; for participants without site information, global median/mode imputation was applied. This approach was motivated by the multi-site structure of the cohort and prior work using within-site imputation strategies in multi-site analyses [97, 98]. PCA was performed in R using prcomp with centering and scaling enabled [113]. The resulting components were highly robust to sample restriction: PCA in complete cases (participants with no missing data) was highly consistent with PCA in the full cohort (*r ≥* 0.98; Supplementary Figure S12), and PCA within the mediation-analysis cohort showed similarly consistent components (*r ≥* 0.95; Supplementary Figure S11).

Component retention was guided by inspection of the scree plot, cumulative variance explained (*>* 5%), and interpretability of component loadings. The first three components captured the largest share of variance and were examined in subsequent descriptive analyses. Because the sign of PCA components is arbitrary, component scores and loadings were oriented post hoc for interpretability using family history group differences. Components were sign-flipped only if FH+ mean scores were lower than FH*−* mean scores in both females and males. This procedure aimed to align higher component scores with the FH-associated direction while preserving variance structure and inter-subject relationships. Because PCA sign is arbitrary, this reorientation was performed only to aid interpretation and did not alter the variance explained, model fit, or statistical evidence for any associations.

To descriptively characterize associations with family history of SUD, component scores were compared between FH+ and FH*−* individuals within each sex using Cohen’s *d* and two-sample *t*-tests. *P*-values were adjusted using the Benjamini–Hochberg false discovery rate (FDR) procedure across the six primary PC1–PC3 component-by-sex comparisons; descriptive PC1–PC10 supplement analyses were corrected across all 20 component-by-sex comparisons. These analyses provided a descriptive assessment of associations corresponding to the a-path in mediation models and were not used to define the PCA solution.

Based on variance explained, interpretability, and the magnitude of family history-related differences, the first three components were retained for primary mediation analyses. Components four and five were not included in mediation models and are presented for descriptive characterization of variance structure and loading patterns. Final component labels were assigned based on dominant loading patterns: PC1 (economic hardship; EH), PC2 (family and school neglect; FSN), and PC3 (parental mental illness; PMI). PCA scores for these retained components were merged into the mediation dataset and entered directly as mediators in structural equation models.

#### 5.4.2 Structural Equation Modeling

Multigroup mediation models with biological sex as the grouping variable were fitted using structural equation modeling (SEM) in R (lavaan) with robust maximum likelihood estimation. A multigroup framework was used to allow all parameters to vary freely by sex without imposing linear interaction constraints. Mediators were modeled in parallel and allowed to covary, such that residual covariances among the environmental risk factors were freely estimated.

For each transition energy (TE) outcome (1 global, 9 network-level, and 86 regional measures), the following model was specified:

- **Exposure (X):** family history of SUD (FH+ vs. FH*−*)
- **Mediators (M):** three environmental principal components (economic hardship, family–school neglect, and parental mental illness), modeled simultaneously
- **Outcome (Y):** TE measure

This yielded 96 outcome-specific models with sex-specific parameter estimates for each TE outcome. All TE outcomes were z-scored prior to modeling.

Parallel mediation was specified such that each mediator was regressed on family history of SUD (*a*-paths), and each TE outcome was regressed simultaneously on all mediators and family history of SUD (*b*-paths and the direct effect). The direct effect (*c^′^*) represents the association between family history of SUD and TE after accounting for all mediators. Specific indirect effects for each mediator were computed as the product of the corresponding *a*- and *b*-paths (*a_i_b_i_*), and these were summed to obtain the total indirect effect (Σ*_i_a_i_b_i_*). The total effect (*c*) was defined as the sum of the direct and total indirect effects (*c* = *c^′^* + Σ*_i_ a_i_b_i_*).

#### 5.4.3 Model Covariates and Adjustment Variables

All mediation paths (i.e., a, b, and c*^′^* paths) were adjusted for: age (z-scored), pubertal status (binary: started vs. not started; see Section 5.1.5), and race/ethnicity using dummy-coded variables (White, Black, Hispanic, and Other). Due to the limited number of Asian subjects, Asian subjects were included as “Other”.

For paths in the model that had TE as the outcome (i.e., b and c*^′^* paths), we additionally adjusted for imaging-related covariates, including in-scanner head motion and MRI scanner manufacturer. To account for head motion, a known confound in pediatric neuroimaging, we included each subject’s mean framewise displacement (FD; z-scored), calculated across all non-outlier time points used to derive subject-specific centroids. Scanner manufacturer was included as a binary variable (GE vs. Siemens).

To account for non-independence among related individuals, family ID was incorporated as a clustering variable, and standard errors were estimated using cluster-robust methods. Participants in the ABCD Study include siblings and other relatives who share a family identifier; thus, observations from the same family are not statistically independent. We specified family ID as the clustering variable in SEMs estimated using maximum likelihood with robust standard errors. Under this approach, point estimates are obtained from a conventional single-level SEM, while standard errors and associated test statistics are adjusted for clustering using a sandwich-type estimator, allowing for arbitrary within-family correlation among residuals.

#### 5.4.4 Statistical Assessment

Within-sex path coefficients (*a*-paths, *b*-paths, and (c’)) were evaluated using Wald tests with maximum likelihood and robust standard errors (family ID as a clustering variable for cluster-robust SEs) as each corresponds to a single regression parameter directly estimated by the model. Specific indirect effects, total indirect effects, total effects, and female–male contrasts in these derived quantities were assessed using Monte Carlo simulation (20,000 draws) from the asymptotic multivariate normal distribution of model parameter estimates [56]. Confidence intervals were derived from the empirical percentile distribution of simulated effects, and two-sided (p)-values were computed from the proportion of simulated values crossing zero.

Sex differences in directly estimated paths (*a*, *b*, and c’) were tested using Satorra–Bentler scaled likelihood ratio tests (LRTs), comparing unconstrained multigroup models with models in which the parameter of interest was constrained to equality across sex [55]. Sex contrasts for derived quantities (specific indirect effects, total indirect effects, and total effects) were computed as the female–male difference in Monte Carlo draws and evaluated using the same percentile-based confidence intervals and two-sided (p)-values described above.

Multiple-comparison correction was applied hierarchically by analytic level and inferential class. Global, network, and regional TE were treated as separate inferential tiers. Within each level, Benjamini–Hochberg (BH) false discovery rate (FDR) correction was applied separately to three families: (i) path-level tests (*a*-paths, *b*-paths, and direct effects), (ii) mediation-effect tests (specific indirect effects, total indirect effects, and total effects), and (iii) formal sex-difference tests [57]. At the global level, corresponding within-sex tests from females and males were pooled within each inferential family. At the network level, correction was applied across the 9 networks within each combination of effect type and sex (for within-sex tests) or contrast type (for sex-difference tests). The same logic was applied at the regional level, correcting across the 86 regions within each corresponding stratum (see Supplementary Section S3.3.5 for additional details).

### 5.5 Exploratory policy-linked projection analysis

To contextualize the environmental mediation findings within broader public-health policy environments, we conducted an exploratory policy-linked projection analysis relating state-level policy indicators to economic hardship (EH) and, indirectly, to TE. This analysis was associational and did not refit the main mediation SEMs. Instead, it combined policy–EH estimates from mixed models with sex-specific EH*→*TE paths already estimated in the mediation analysis.

State-level policy indicators were drawn from ABCD Release 5.1 Linked External Data (LED), including Medicaid expansion under the Affordable Care Act (ACA), Good Samaritan laws, recreational cannabis legalization, naloxone access, and prescription drug monitoring program (PDMP) must-access requirements. Because residential state was not consistently available at the participant level, policy exposure was assigned using the state of the ABCD site at which the participant was assessed. Policy variables were coded as binary indicators, with policy absent coded as 0 and policy present coded as 1 at baseline.

Several LED indicators represented overlapping or nested policy implementations, so we used a reduced-domain model with five policy domains: ACA Medicaid expansion, any Good Samaritan law, recreational cannabis legalization, naloxone access based on standing/protocol access, and PDMP must-access requirements. In the full ABCD cohort with environmental PC scores and complete covariates (*N* = 9,157), we fit a single linear mixed model predicting EH. The model included FH, sex, the five policy indicators, FH*×*Policy interactions, age, pubertal initiation, race/ethnicity indicators, and a random intercept for state. The primary estimates were the FH*×*Policy interaction coefficients, representing the difference in the FH+ minus FH*−* EH gap for policy-present versus policy-absent state environments. Benjamini–Hochberg false discovery rate (FDR) correction was applied across the five interaction terms.

Policy-linked TE projections were calculated by propagating the estimated FH*×*Policy effect on EH through the sex-specific EH*→*TE path from the main multi-group mediation models:

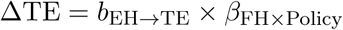

Uncertainty was quantified using Monte Carlo simulation with 20,000 draws, sampling *β*_FH_*_×_*_Policy_ and *b*_EH_*_→_*_TE_ independently from normal approximations based on their estimated coefficients and standard errors. This approximation did not account for covariance between the policy and mediation estimates.

Observed FH+ minus FH*−* TE gaps in the imaging cohort were used as reference gaps for percent gap-closure calculations:

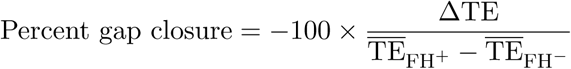

Percent gap closure was not interpreted when the observed FH TE gap was near zero 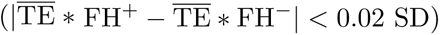. For the combined projection, FH*×*Policy coefficients were summed across FDR-significant policy domains, with the standard error approximated as 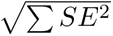 under an independence approximation. All projections assume linearity, additivity for the combined scenario, and independence between policy and mediation estimates, and should be interpreted as exploratory, model-based projections through the EH pathway rather than causal estimates of policy effects.

## Software and Implementation

Custom scripts were developed using MATLAB (R2023a), Python (3.11), and R (4.5.2). All mediation analyses were conducted in R: structural equation models were fit with lavaan (version 0.6.20; [54]), and Monte Carlo confidence intervals were obtained using MASS (mvrnorm; [114]). MATLAB was used for *k*-means clustering and network control theory-based transition energy (TE) calculations, based on publicly available code from Cornblath et al. [110] (https://github.com/ejcorn/brain_states) and Singleton et al. [111] (https://github.com/singlesp/energy_landscape). Python (version 3.11) was used for figure generation and brain map rendering using brainmontageplot (v1.4.2; https://github.com/kjamison/brainmontageplot).

## Data availability

Access to ABCD study data is restricted to protect participants’ privacy. Users must create an account through the National Institute of Mental Health Data Archive and they may then complete the necessary steps to gain access. Researchers with access to the ABCD data will be able to download the data: https://nda.nih.gov/study.html?id=1368. The data used in this study are from the Adolescent Brain Cognitive Development (ABCD) Study (https://abcdstudy.org), a longitudinal study supported by the NIH and other federal partners. Access to the ABCD data is subject to controlled access and requires registration through the National Institute of Mental Health Data Archive (NDA): https://nda.nih.gov. Researchers with approved access can download the raw and processed data used in this study. The ABCD neuroimaging data used in this study were downloaded from https://doi.org/10.15154/1504041 and the non-imaging data were from the 5.1 release (https://doi.org/10.15154/z563-zd24). Preprocessed imaging derivatives provided by Chen et al. (2022) [99] and Ooi et al. (2022) [52] were obtained from NDA Study 1368: https://nda.nih.gov/study.html?id=1368.

## Acknowledgments

The authors would like to thank the ABCD study community of researchers and staff involved in data collection, and especially the ABCD participants and their families for their commitment to the study. The authors also thank B.T. Thomas Yeo’s lab, specifically Leon Qi Rong Ooi, for providing the preprocessed MRI data. This research was supported in part by a grant from the Ann S. Bowers Foundation to L.S., C.G., and A.K. through the Ann S. Bowers Women’s Brain Health Initiative. M.H. is supported by the National Research Fund, Luxembourg. RF1 MH123232 (A.K.) was provided by the National Institute of Mental Health of the National Institutes of Health. T32 DA03980 (L.S.) was provided by the National Institute on Drug Abuse of the National Institutes of Health. Grant AA028840 (Q.Z.) was provided by the National Institute on Alcohol Abuse and Alcoholism of the National Institutes of Health. Q.Z. also received a Brain and Behavior Research Foundation Young Investigator Grant.

Data used in the preparation of this article were obtained from the Adolescent Brain Cognitive Development (ABCD) Study (https://abcdstudy.org), held in the NIMH Data Archive (NDA). This is a multisite, longitudinal study designed to recruit more than 10,000 children age 9-10 and follow them over 10 years into early adulthood. The ABCD Study® is supported by the National Institutes of Health and additional federal partners under award numbers U01DA041048, U01DA050989, U01DA051016, U01DA041022, U01DA051018, U01DA051037, U01DA050987, U01DA041174, U01DA041106, U01DA041117, U01DA041028, U01DA041134, U01DA050988, U01DA051039, U01DA041156, U01DA041025, U01DA041120, U01DA051038, U01DA041148, U01DA041093, U01DA041089, U24DA041123, U24DA041147. A full list of supporters is available at https://abcdstudy.org/federal-partners.html. A listing of participating sites and a complete listing of the study investigators can be found at https://abcdstudy.org/consortium_members/. ABCD consortium investigators designed and implemented the study and/or provided data but did not necessarily participate in the analysis or writing of this report. This manuscript reflects the views of the authors and may not reflect the opinions or views of the NIH or ABCD consortium investigators. The ABCD data repository grows and changes over time. Access to ABCD study data is restricted to protect participants’ privacy. Users must create an account through the National Institute of Mental Health Data Archive and they may then complete the necessary steps to gain access. Researchers with access to the ABCD data will be able to download the data: https://nda.nih.gov/study.html?id=1368.

## Supplementary Information

## S1 Network control theory analysis

### S1.1 Calculation of transition energy

Following previous work [105, 106], we applied *k*-means clustering to regional rsfMRI time series data (86-region atlas) to identify *k* recurring patterns of brain activity, termed “brain states”. For each subject, we assigned each individual frame to a brain state and calculated subject-specific brain state centroids. We then applied NCT to calculate the global, network, and region-level TE required to complete brain state transitions. For this, we utilized a group-average structural connectome (derived from diffusion MRI from a subset of individuals in this dataset) as previously done [103], and as supported by observations that functional abnormalities precede white matter changes in FH+ youth [20, 115] (see Supplementary Materials in our previous paper ([45]) for a replication of our results using individual structural connectomes). We calculated both the pairwise and mean TE for all levels of analysis: global, network and regional. Initial TE calculations result in “pairwise regional TE” values, i.e., 86 (number of regions) *k × k* matrices. For each matrix, we summed pairwise regional TE of all regions belonging to each of the seven Yeo networks [48] plus subcortical and cerebellar networks to yield “pairwise network TE” values, i.e., 9 (number of networks) *k × k* matrices. We derived a single “pairwise global TE” *k × k* matrix by summing across all 86 regions pairwise TEs for each transition *k × k* matrix. We also averaged all entries in pairwise TE matrices to derive mean TE values for each region, network and globally, resulting in: 86 mean regional TE values, 9 mean network TEs, and a single value for mean global TE. Thus, pairwise TE represents the energy required of a given region, network or whole brain to complete transitions between each pair of brain states, whereas mean TE represents the energy required to transition across the entire state space (i.e., all pairwise transitions).

It is important to distinguish the four brain states from the nine networks used to calculate TE values, although both derive from a common nine-network parcellation (Yeo 7-networks plus subcortical and cerebellar networks) [48]. Brain states are the four *k*-means clusters, assigned to the network whose high or low activity best explained each centroid. These descriptive labels did not influence analyses and are denoted with “+” or “–” to indicate above- or below-mean activity (e.g., DMN+, VIS–). Network TE values, by contrast, are calculated by summing regional TE across all regions in a given network and are referenced using the nine network names alone (e.g., DAT, VIS, DMN).

### S1.2 Clustering results

We based our choice of *k* =4 for the *k*-means clustering analysis of brain states on our previous analysis in the same cohort. Here, we briefly summarize the procedure used to select *k* =4 in our prior study [45]. We performed 10 repetitions of *k*-means clustering for *k* =2 to *k* =13. The maximum value (*k* =13) was chosen such that the square of the number of clusters was less than the minimum number of frames available for any subject.

We quantified the variance explained by clustering as the ratio of between-cluster variance to total variance in the data [105, 106, 116]. To select *k*, we plotted the gain in explained variance as *k* increased and identified the point at which additional clusters provided diminishing returns. The elbow of the variance-explained curve occurred between *k* =4 and *k* =5. In addition, after *k* =4, the increase in explained variance fell below 1%, the threshold used previously for selecting *k* [105, 106]. In the present analysis, we chose to use *k* =4 as we did in the main results of our previous work in which we also replicated our findings in *k* =5.

Given the slight change in our cohort relative to the previous analysis, we re-ran *k*-means clustering to identify *k* =4 brain states in the current sample. We followed the same procedure as previously: we performed 10 repetitions of *k*-means clustering at *k* =4 and selected the solution with the lowest error. To ensure that this solution represented a consistent global minimum, we repeated the entire procedure 10 times and compared the adjusted mutual information (AMI) across the resulting partitions. The partition with the maximum summed AMI across all other partitions was selected for analysis. This process confirmed that the *k*-means solution was highly stable, with AMI *>* 0.999.

As expected, the clustering results were consistent with our previous work. The identified brain states comprised two pairs of anti-correlated activity patterns (i.e., meta-states): one dominated by high- and low-amplitude activity in the default mode network (DMN+/DMN*−*) and the other by high- and low-amplitude activity in the visual network (VIS+/VIS*−*).

### S1.3 Replication of the sex-by-family-history interaction effect on global TE

To replicate the sex *×* family-history interaction effect on global TE reported in our previous study [45] within the slightly expanded cohort used here, we fit an ANCOVA model predicting global TE in FH+ and FH*−* youth. As in the prior analysis, the ANCOVA model included sex, family history of SUD, sex *×* family history of SUD, age, race/ethnicity, parental education, household income, prenatal substance exposure, parental mental health, in-scanner motion, pubertal stage, MRI scanner manufacturer, family history of SUD *×* income and sex *×* puberty interaction. The analysis sample comprised *N* = 1,757 youth (54.3% female), aged 107–132 months (*≈*8.9–11.0 years), with complete data on all model terms. Benjamini–Hochberg FDR correction (*p*_FDR_) was applied across all model terms.

We again observed a significant sex *×* family-history interaction effect that survived FDR correction, such that FH+ females had greater global TE than FH*−* females, whereas FH+ males had lower global TE than FH*−* males.

**Figure S6:**
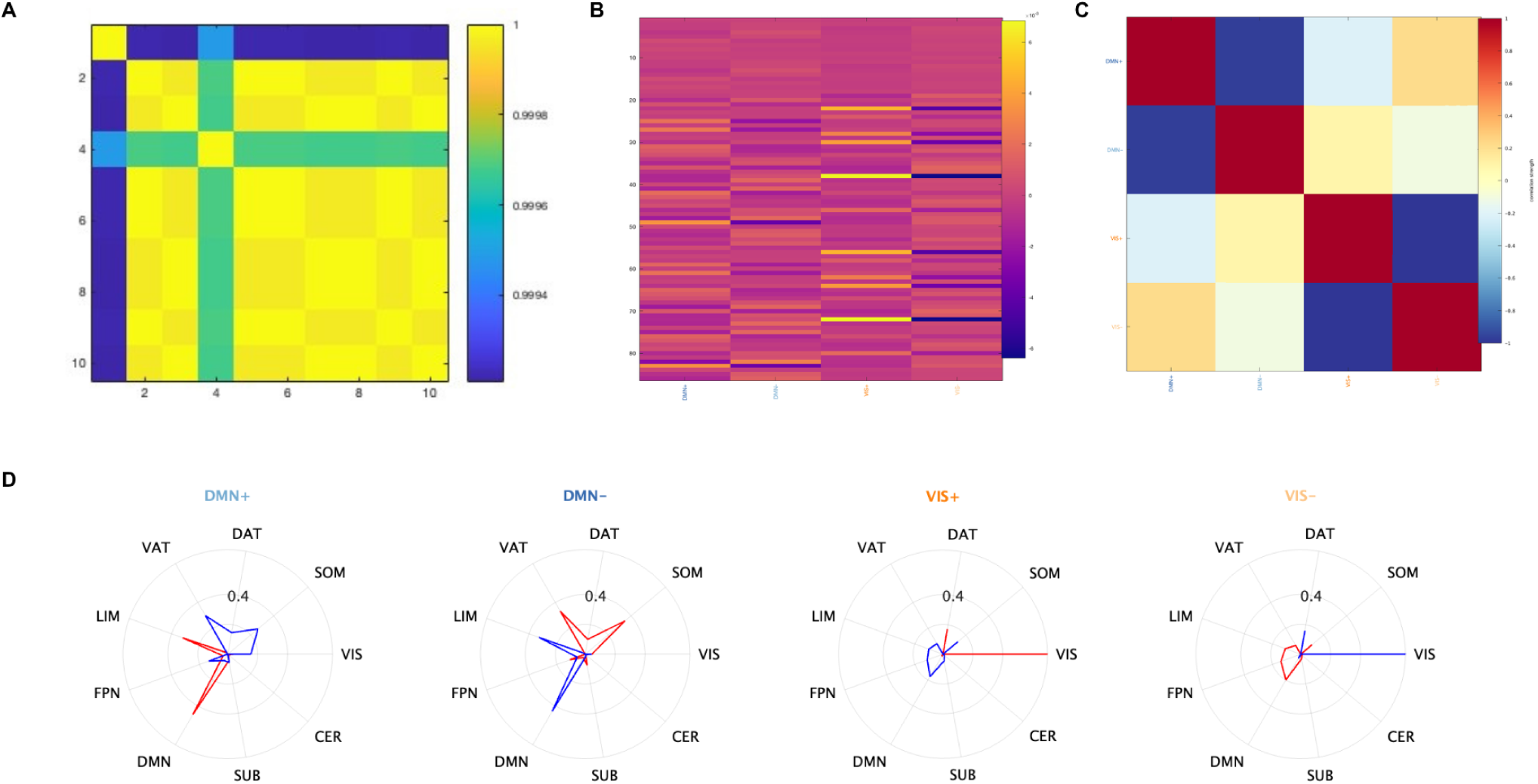
**(A)** Adjusted mutual information (AMI) shared between 10 independently generated partitions of the data at *k* =4. Values range from 0 to 1, with 1 indicating identical partitions. AMI between all partitions was *>* 0.999. **(B)** Centroids of each cluster, defined as the mean of all time points assigned to that cluster across all subjects. **(C)** Pearson correlation values between each pair of clusters. Dark blue squares indicate strong negative correlations between clusters belonging to the same meta-state. **(D)** Cosine similarity with canonical resting-state networks [48], calculated separately for the positive (high-amplitude) and negative (low-amplitude) components of each brain state’s group-average centroid. Each brain state was assigned the label of the network with the maximal cosine similarity, with the sign indicating whether the maximal similarity was determined from the positive (+) or negative (-) component. CER = cerebellum, DAT = dorsal attention network, DMN = default mode network, FPN = frontoparietal network, LIM = limbic network, SOM = somatomotor network, SUB = subcortex, VAT = ventral attention network, VIS = visual network.

## S2 Principal components analysis

### S2.1 PCA construction details

Principal component analysis (PCA) was run on environmental variables derived from the entire cohort of the ABCD Study (release 5.1). Most variables were derived from baseline data, with retrospective breastfeeding exposure obtained from the 3-year follow-up when available. Overall, 43 variables entered the PCA after pre-processing. Variable names and short definitions are given in Supplementary Table 1.

For several constructs, composites were computed prior to PCA, including: trauma-domain summaries (any exposure within domain, with community and domestic violence items pooled for the violence domain); partnership status from marital codes; household caregiver and unemployment indicators from employment codes; parenting row-means (communication mean, warmth mean); means for basic-needs instability, access-to-care barriers, and neighborhood safety; and family-climate and conflict aggregates.

**Figure S7:**
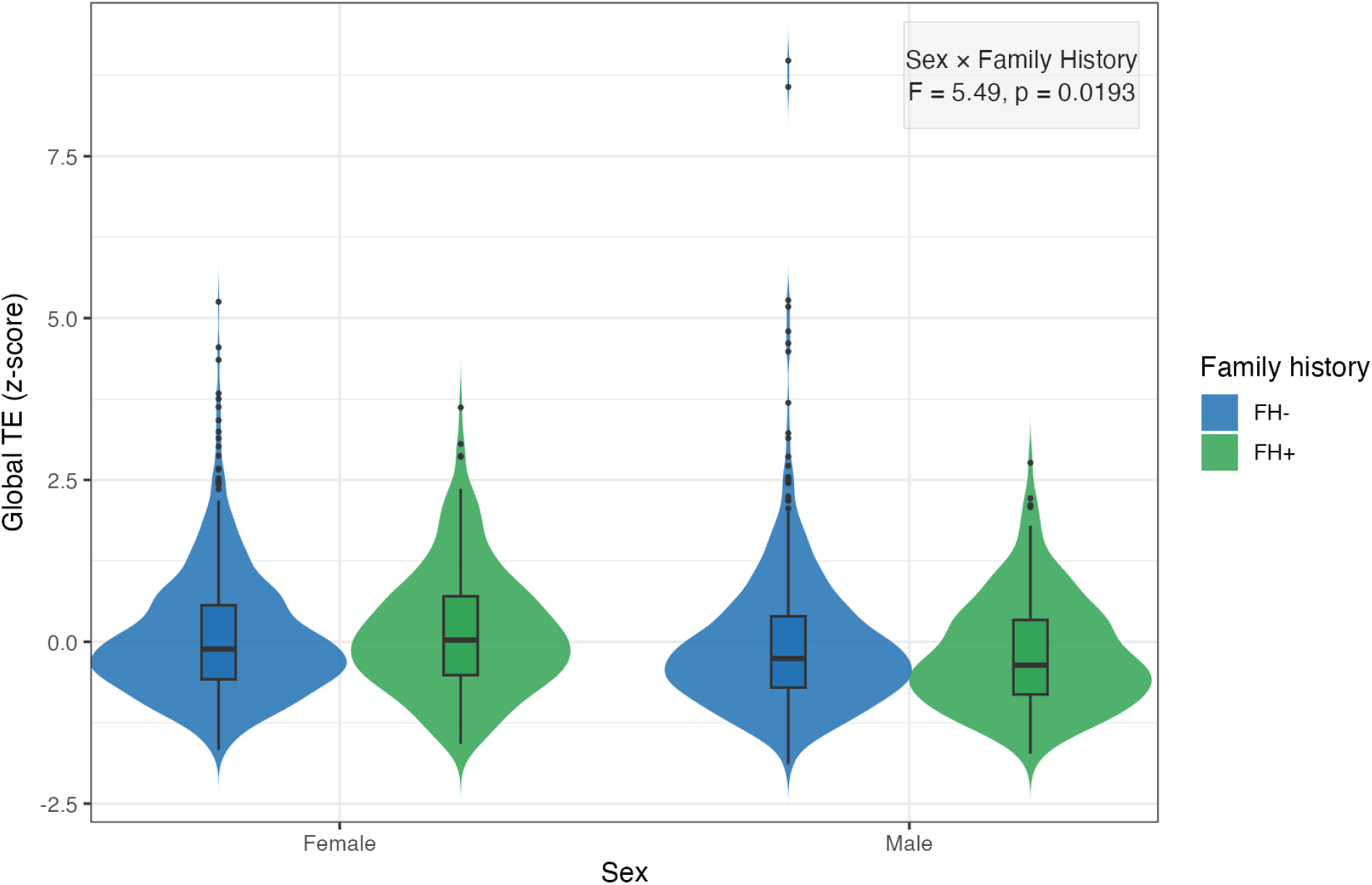
Sex-by-family history interaction effect on global transition energy.

**Table S2:**
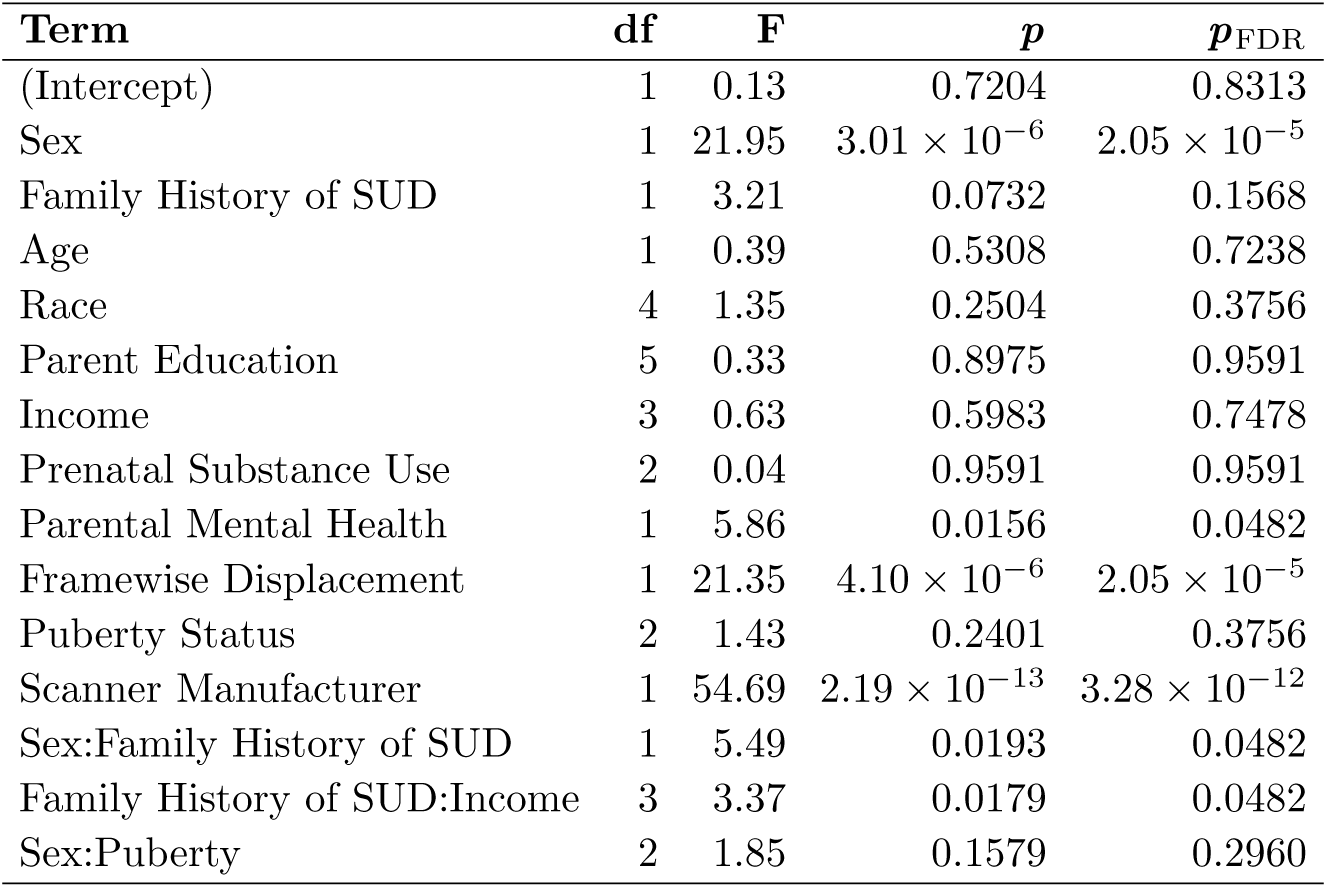
ANCOVA of global transition energy.

To reduce skew in non-negative, heavy-tailed metrics, a log-transformation (log1p) was applied *before* imputation to `neighborhood_violent_crime, pop_density, prox_major_road, and screen_week`.

Missing values were then imputed once: for continuous variables, the median of non-missing values within a participant’s study site; for binary/categorical variables, the mode within site. If site was missing or a site-specific value could not be computed, cohort-wide median (continuous) or mode (binary/categorical) was used. Any variable with zero variance after imputation was excluded. The highest missingness of all variables was 14.87% for substance use during breastfeeding (1,765 / 11,868), then income to needs ratio (10.45%), area deprivation index percentage (7.41%), and parent depression (6.34%).

PCA was performed in R using prcomp on the fully observed (post-imputation) matrix with each input variable standardized (mean 0, SD 1) before extraction of components. The first ten principal components (PCs) were retained for descriptive reporting (loadings, variance, group contrasts); primary mediation models used PC1–PC3 as environment mediators. In the full sample, PC1 explained 11.3% of total variance in the scaled inputs, and PCs 1–5 together explained 30.9%.

Component scores and loadings were subsequently oriented with respect to family history of substance use disorder: for each PC, the sign was reflected when, in both sexes separately, the mean score was lower in FH+ than in FH*−* so that higher scores correspond to the direction associated with FH+ in that situation. This sign convention affects interpretation of loadings and scores but not the proportion of variance explained.

### S2.2 Assessment of collinearity among environmental variables prior to PCA

Before PCA, we examined correlations among all environmental input variables to assess overlap and potential multi-collinearity. Using the same imputed data matrix entered into the PCA, we computed Pearson correlations between all pairs of variables and visually inspected the correlation matrix for clustering and highly overlapping measures. In the final PCA input set, off-diagonal correlations ranged from *r* = *−*0.473 to *r* = 0.73, indicating moderate shared variance without extreme collinearity. These results supported the suitability of the retained variables for PCA.

**Figure S8:**
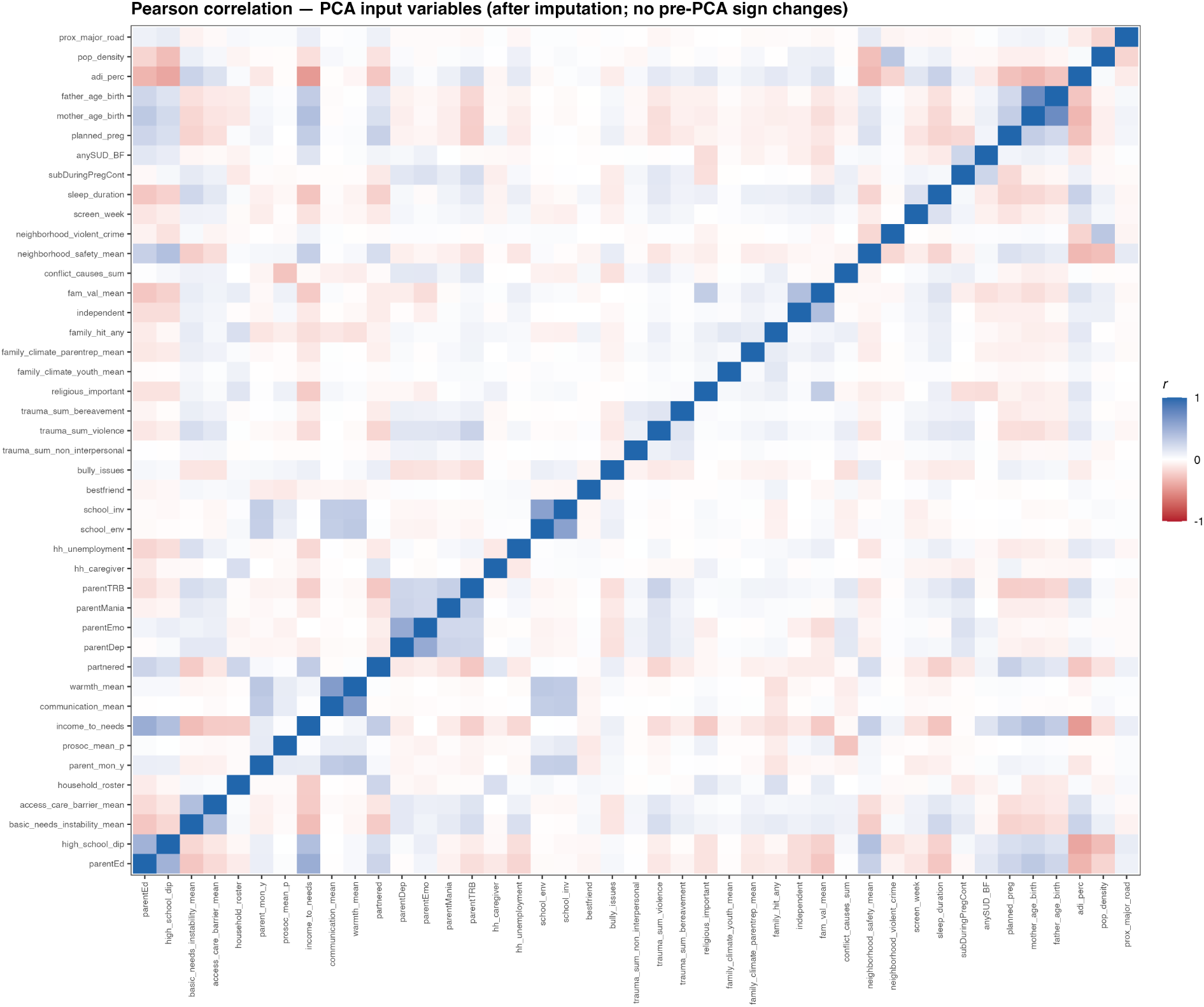
Correlation matrix of environment variables entered into PCA.

### S2.3 Environmental variables included in the principal component analysis

**Table S3:**
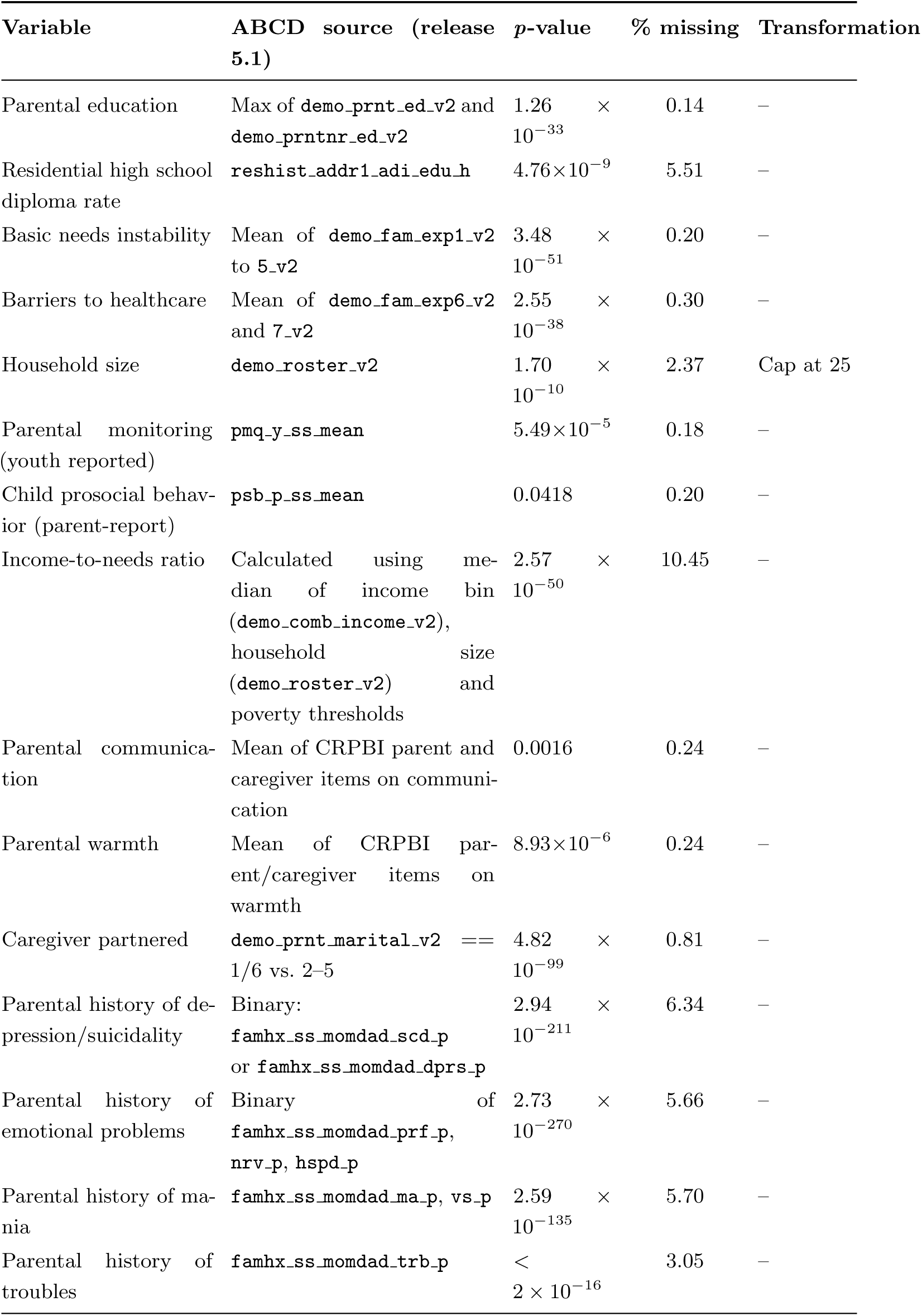
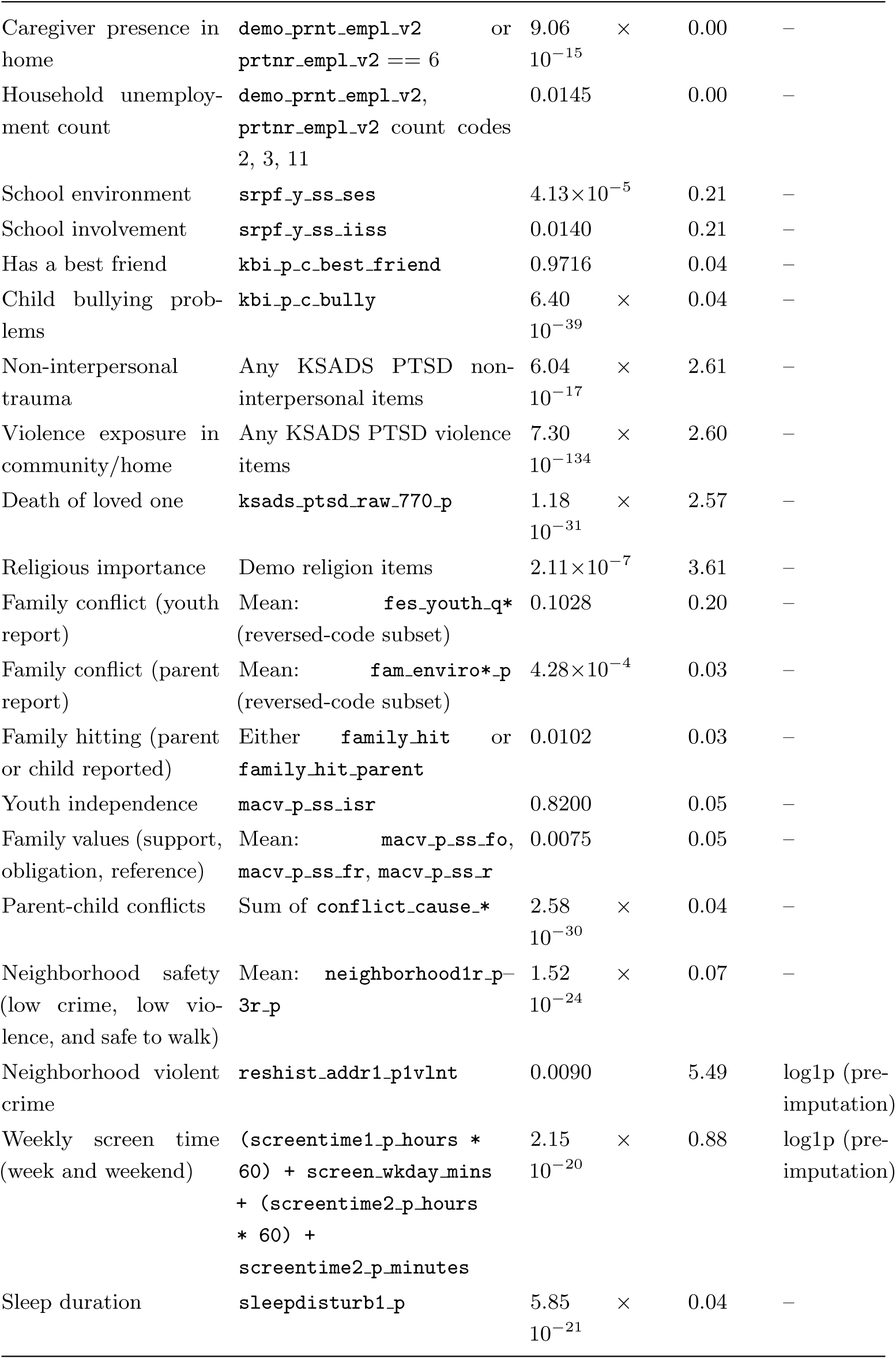
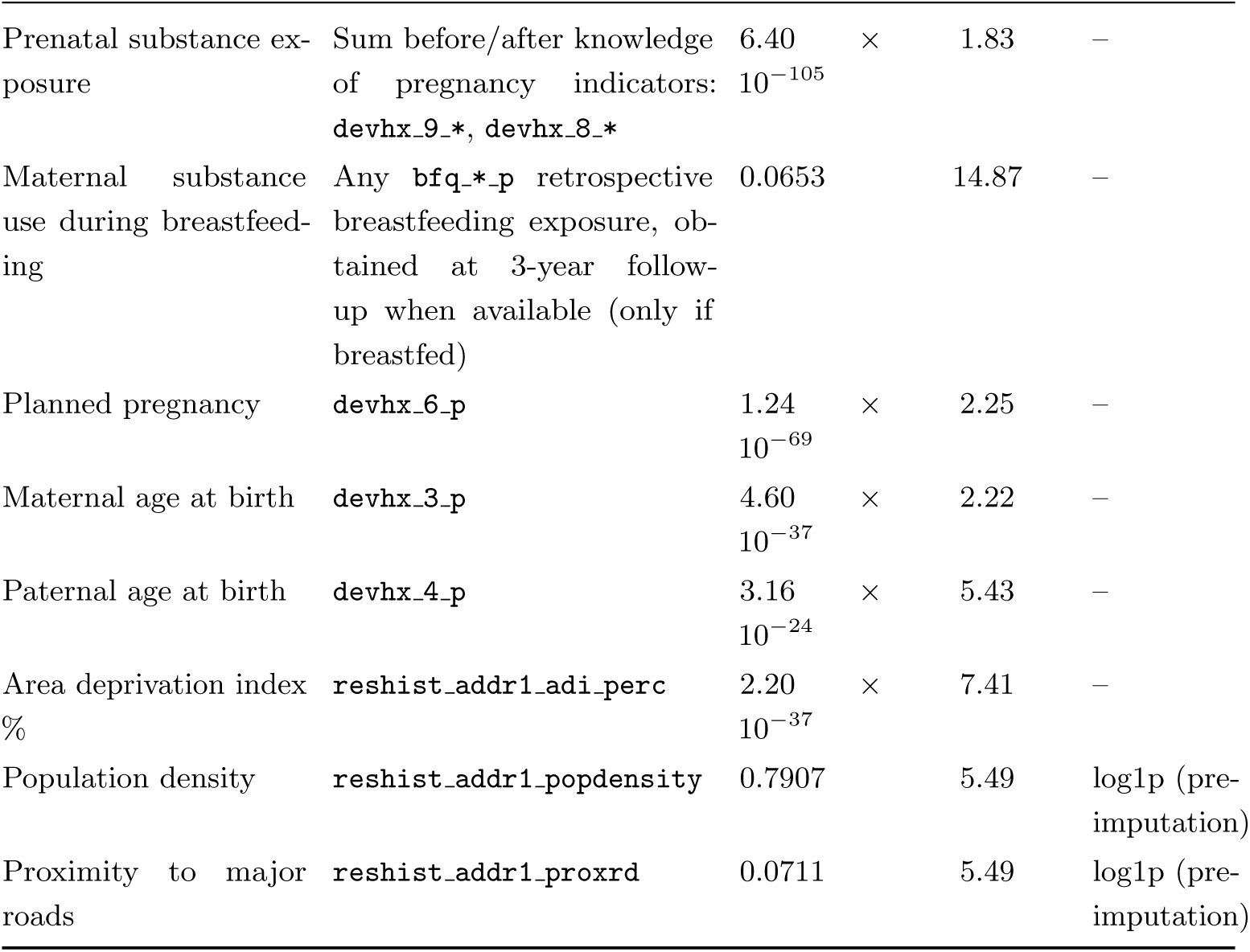
Environment PCA inputs. . ABCD fields (release 5.1), brief label, *p*-value from *t*-test or *χ*^2^ test, missingness before imputation, and transformation or composite construction.

### S2.4 PCA loadings across environmental components

The top 10 principal components (PCs) cumulatively explained 45.2% variance. The first three PCs captured the most interpretable and conceptually relevant dimensions of environmental variation (economic hardship, family-school neglect, and parental mental illness) and were therefore retained as mediators in the main analyses. In contrast, PCs 4–10 showed more heterogeneous loading patterns and were not included as mediators, as they explained less variance and were less readily interpretable as coherent latent constructs. Below, we report the ten variables with the largest absolute loadings for each of the first ten PCs. Within each component, variables are ordered by absolute loading magnitude, with red bars indicating positive loadings and blue bars indicating negative loadings.

**Figure S9:**
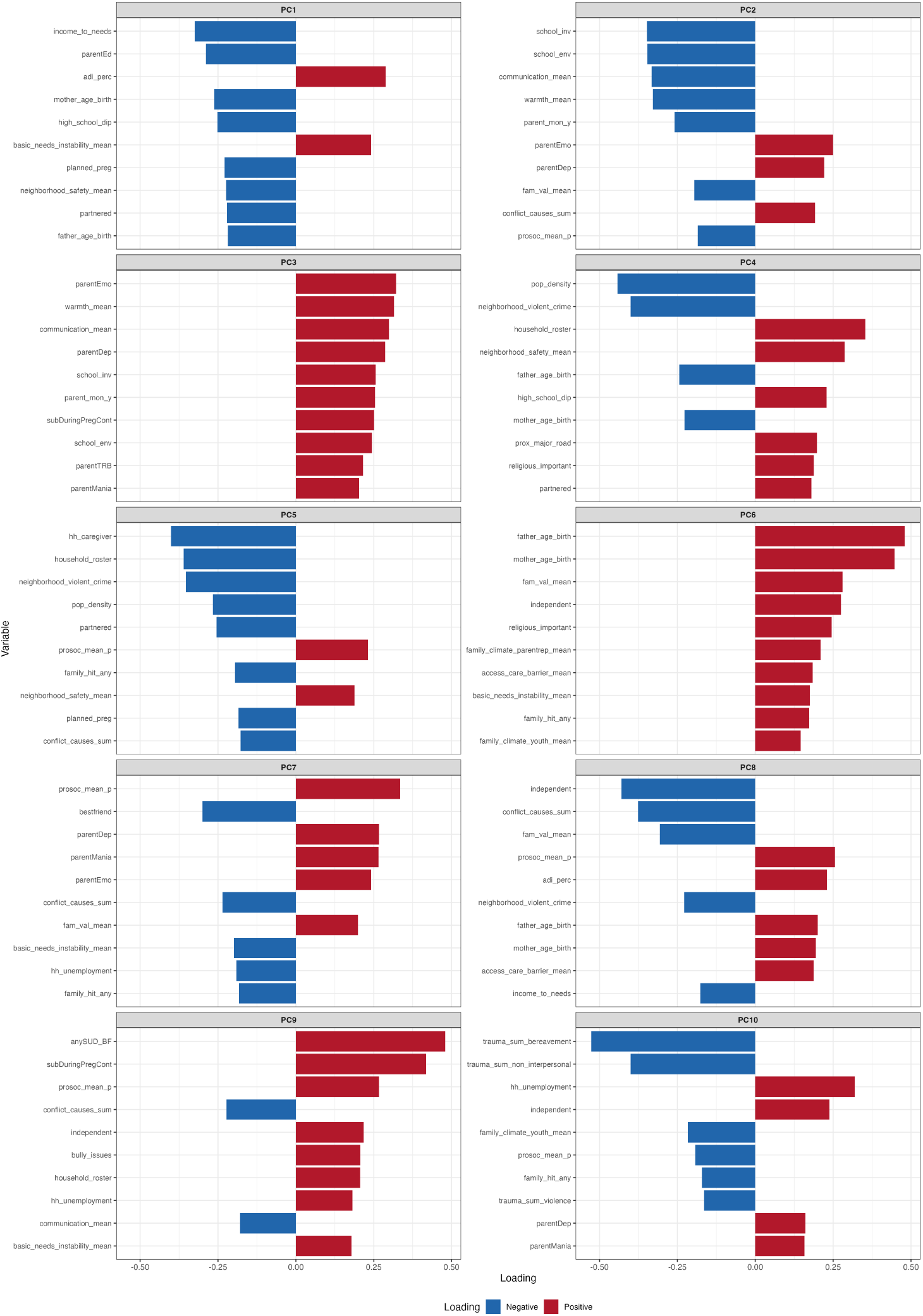
Top 10 loadings (absolute value) of first 10 principal components of environmental risk factors.

### S2.5 Family history of SUD differences across the top 10 environmental principal components

We compared FH+ and FH*−* youth on each of the first 10 environmental principal components separately in females and males within the mediation cohort. Group differences were summarized using Cohen’s *d*, and significance was assessed with Benjamini–Hochberg false discovery rate (BH-FDR) correction across 20 tests (10 PCs *×* 2 sexes).

The largest and most consistent differences related to family history of SUD were observed for PC1, PC2, and PC3 in both sexes. FH+ youth showed higher scores than FH*−* youth on all three components, with medium-to-large effect sizes in both females and males (all *p*_FDR_ *<* 0.001). These results support the selection of PC1–PC3 as the primary environmental mediators in the main analyses, as they were both interpretable and strongly associated with family history status.

In contrast, family history of SUD group differences for PCs 4–10 were smaller and less consistent across sex. A subset of later components showed nominal or FDR-surviving differences in only one sex, including PC4 and PC5 in males, PC7 in both sexes, and PC8 and PC9 in females, whereas the remaining components showed weak or negligible family history effects. Overall, these findings indicate that the first three PCs captured the most robust family history-related environmental variation in this cohort.

**Figure S10:**
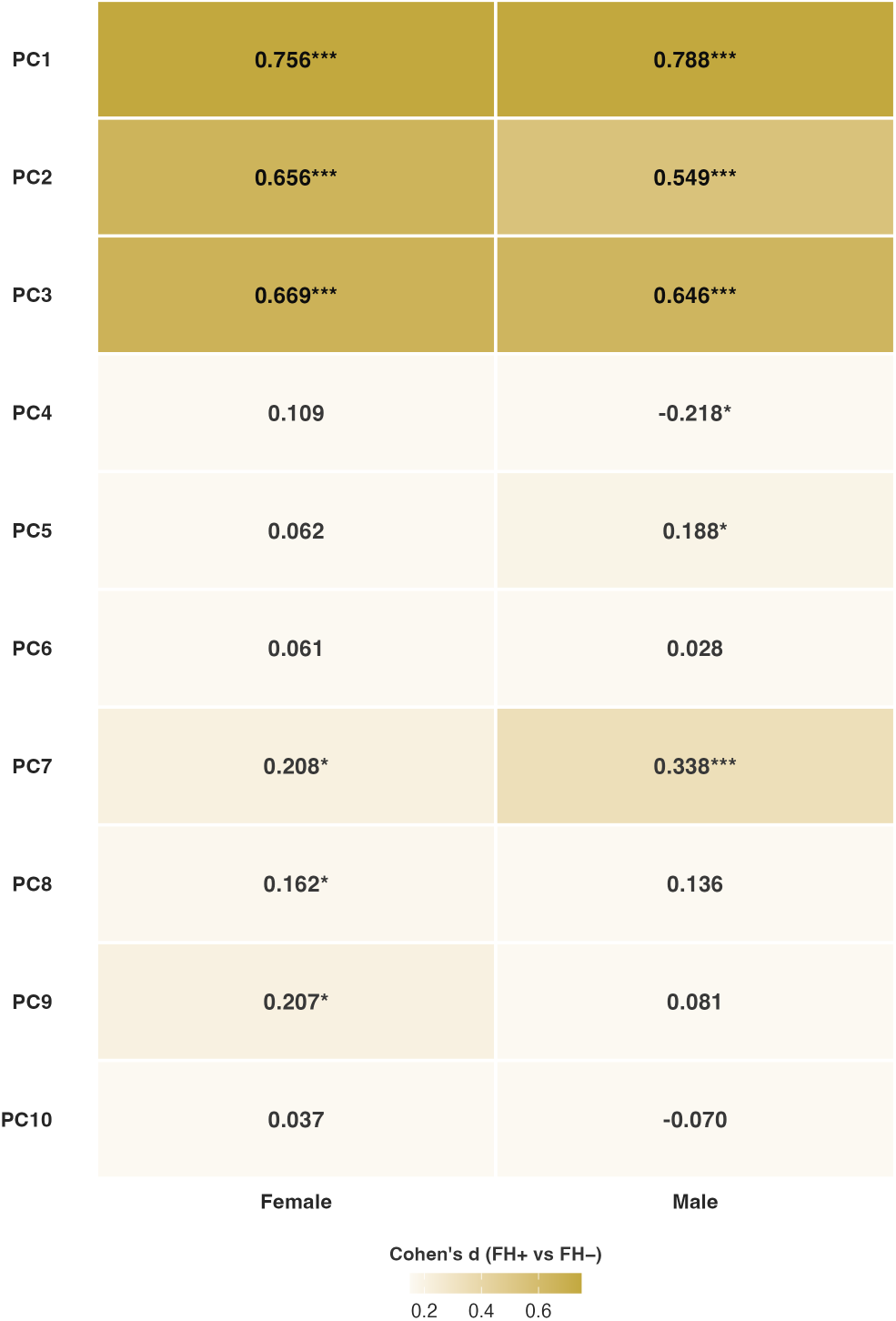
Family history differences across environmental principal components in the mediation cohort. Heatmap showing Cohen’s *d* effect sizes for FH+ versus FH*−* comparisons on environmental PCs 1–10, shown separately for females and males. Positive values indicate higher PC scores in FH+ youth relative to FH*−* youth. Asterisks denote BH-FDR-corrected significance across 20 tests (*^∗^p <* 0.05, *^∗∗^p <* 0.01, *^∗∗∗^p <* 0.001).

### S2.6 PCA fit restricted to the mediation analytic cohort

Main analyses used environment PC scores (PC1–PC3) applied to the full curated environmental sample (with site-level imputation of missing inputs). To assess whether the PC axes are stable when PCA is fit *only* within the cohort used for the NCT and mediation analyses (*N* = 1,835), we repeated PCA on the same 43 input variables using the identical pre-processing and imputation rules on just those subjects. For each component *n ∈ {*1, 2, 3*}*, the mediation-cohort PC*n* was multiplied by *−*1 if its Pearson correlation with main PC*n* was negative, so the heatmap reflects sign-aligned agreement. Overall, the correlation between the top 3 PCs was very high (*r ≥* 0.95).

**Figure S11:**
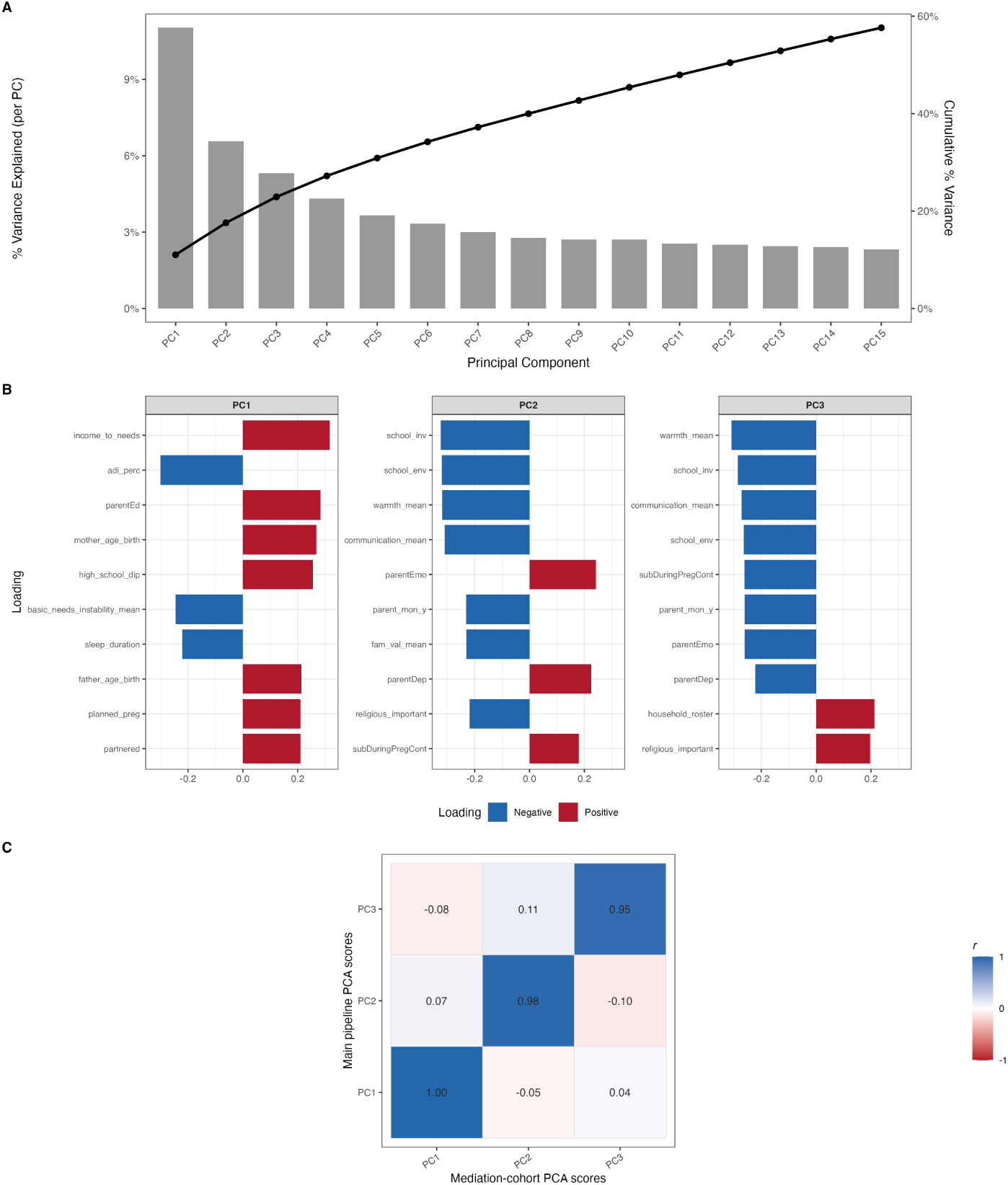
Environment PCA fit within the mediation analytic cohort only (*N* = 1,835; same 43 inputs and pre-processing as the main PCA). **(A)** Scree plot: percent variance explained per principal component (bars) and cumulative percent variance (line). **(B)** Top 10 absolute loadings for PC1, PC2, and PC3 (red = positive loading, blue = negative). **(C)** Correlation matrix comparing PC scores (PC1–PC3; rows) to PC scores from the mediation-only fit (columns; mediation PCs sign-aligned to the main PCs per component). Cell entries are Pearson *r*.

### S2.7 Complete-case Principal Component Analysis

The primary PCA used site-level imputation for missing values on individual input variables. As a sensitivity check, we repeated PCA using only participants with no missing data on any of the 43 environment variables entering the PCA (*N* = 6,862). For each component *n ∈ {*1, 2, 3*}*, PC*n* was multiplied by *−*1 if its Pearson correlation with main PC*n* was negative, so the heatmap reflects sign-aligned agreement with the main PCA. Overall, the Pearson correlation between the top 3 PCs was very high (*r ≥* 0.98) between the PCA results from the main analysis (full cohort with imputation) and complete-case subset.

**Figure S12:**
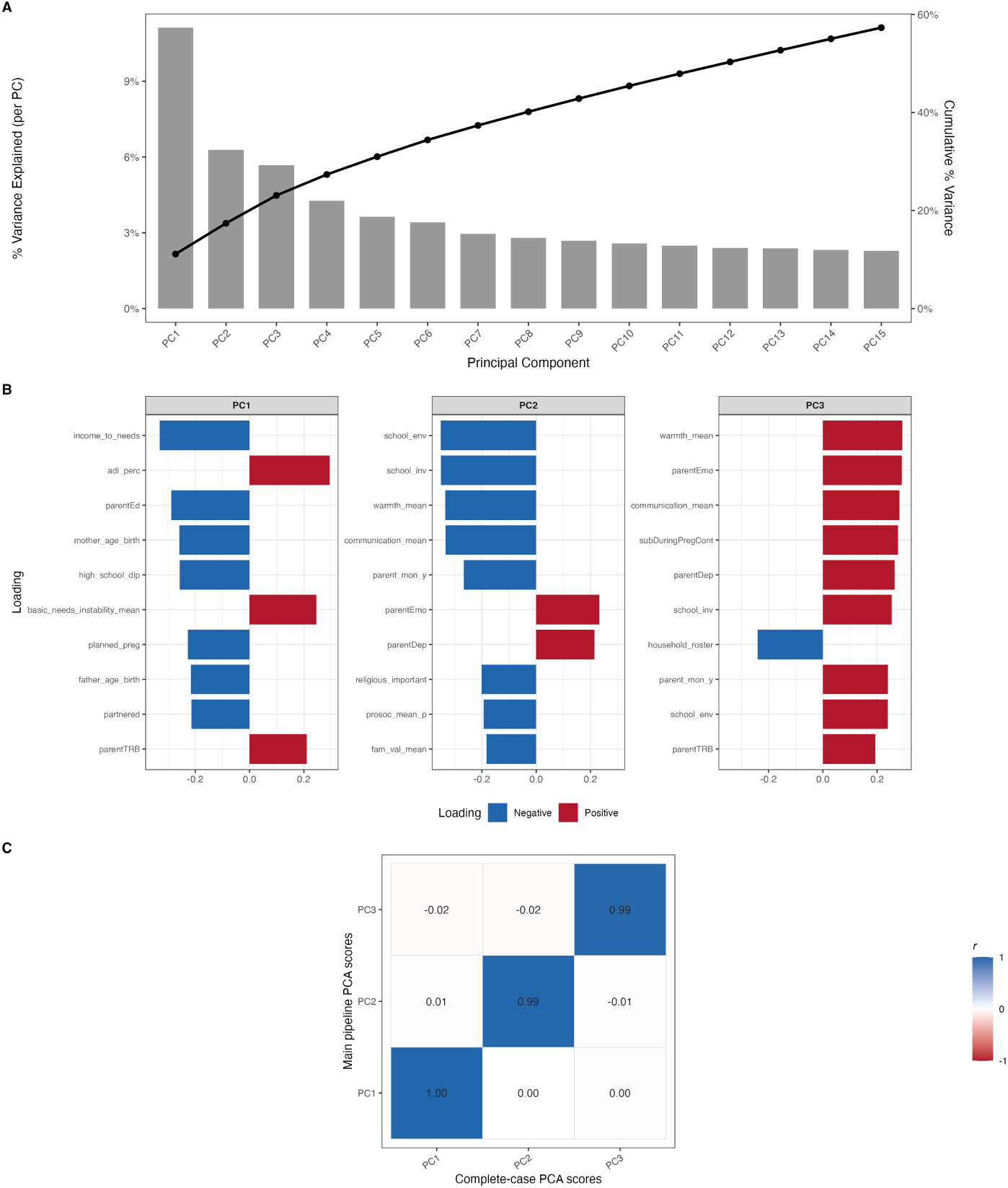
Environment PCA on subjects with complete data on all 43 PCA input variables (no imputation; *n* = 6,862). **(A)** Scree plot: percent variance explained per principal component (bars) and cumulative percent variance (line). **(B)** Top 10 absolute loadings for PC1, PC2, and PC3. **(C)** Pearson correlations between main PC scores (rows) and complete-case PC scores (columns).

## S3 SEM Mediation Analysis

### S3.1 Global TE Mediation Analysis

#### S3.1.1 Table: Global TE mediation results

**Table S4:** Within-sex mediation results for global transition energy. *a*-paths (FH *→* mediator) are reported as fully standardized coefficients, with delta-method standard errors and 95% confidence intervals. *b*-paths (mediator *→* global TE), direct effects (*c^′^*), and indirect and total effects of family history of SUD are shown separately by sex and remain on the original scales of the fitted model. Indirect effects correspond to environmental principal components: EH (economic hardship), FSN (family–school neglect), and PMI (parental mental illness). *a*-paths, *b*-paths, and direct effects were evaluated using Wald tests with maximum likelihood and robust standard errors, with family ID as a clustering variable for cluster-robust SEs. Indirect and total effects were estimated via Monte Carlo simulation (20,000 draws) with percentile-based confidence intervals. Benjamini–Hochberg FDR correction was applied in two pools: path-level within-sex tests (*a*-paths, *b*-paths, and direct effects; 14 tests total) and indirect/total-effect tests across both sexes (10 tests total).

| Sex | Effect | Estimate | SE | z | p | 95% CI | pFDR |
| --- | --- | --- | --- | --- | --- | --- | --- |
| *a-paths (FH → mediator)* |  |  |  |  |  |  |  |
| Male | EH | 0.285 | 0.032 | 8.59 | < 0.001 | [0.223, 0.347] | < 0.001 |
| Male | FSN | 0.245 | 0.032 | 7.37 | < 0.001 | [0.182, 0.309] | < 0.001 |
| Male | PMI | 0.289 | 0.034 | 7.93 | < 0.001 | [0.222, 0.356] | < 0.001 |
| Female | EH | 0.290 | 0.027 | 10.16 | < 0.001 | [0.237, 0.344] | < 0.001 |
| Female | FSN | 0.293 | 0.030 | 9.27 | < 0.001 | [0.235, 0.351] | < 0.001 |
| Female | PMI | 0.291 | 0.029 | 9.31 | < 0.001 | [0.234, 0.349] | < 0.001 |
| *b-paths (Mediator → global TE)* |  |  |  |  |  |  |  |
| Male | EH | -0.038 | 0.020 | -1.96 | 0.050 | [-0.077, 0.000] | 0.070 |
| Male | FSN | 0.013 | 0.022 | 0.58 | 0.563 | [-0.030, 0.056] | 0.563 |
| Male | PMI | 0.053 | 0.024 | 2.22 | **0.027** | [0.006, 0.099] | **0.047** |
| Female | EH | 0.060 | 0.018 | 3.25 | **0.001** | [0.024, 0.096] | **0.002** |
| Female | FSN | 0.025 | 0.020 | 1.26 | 0.206 | [-0.014, 0.065] | 0.240 |
| Female | PMI | 0.050 | 0.023 | 2.16 | **0.031** | [0.005, 0.096] | **0.048** |
| *Direct effects (FH → global TE)* |  |  |  |  |  |  |  |
| Male | Direct | -0.153 | 0.084 | -1.83 | 0.068 | [-0.318, 0.011] | 0.086 |
| Female | Direct | -0.061 | 0.076 | -0.81 | 0.417 | [-0.209, 0.087] | 0.449 |
| *Indirect and total effects (Monte Carlo CI)* |  |  |  |  |  |  |  |
| Male | EH | -0.049 | — | — | 0.049 | [-0.101, -0.000] | 0.081 |
| Male | FSN | 0.011 | — | — | 0.566 | [-0.028, 0.052] | 0.629 |
| Male | PMI | 0.050 | — | — | 0.030 | [0.005, 0.099] | 0.075 |
| Male | Total indirect | 0.013 | — | — | 0.761 | [-0.068, 0.094] | 0.761 |
| Male | Total effect | -0.140 | — | — | 0.046 | [-0.278, -0.002] | 0.081 |
| Female | EH | 0.081 | — | — | **0.001** | [0.032, 0.137] | **0.005** |
| Female | FSN | 0.026 | — | — | 0.202 | [-0.014, 0.068] | 0.252 |
| Female | PMI | 0.045 | — | — | 0.029 | [0.005, 0.090] | 0.075 |
| Female | Total indirect | 0.152 | — | — | < 0.001 | [0.071, 0.237] | **0.002** |
| Female | Total effect | 0.091 | — | — | 0.177 | [-0.041, 0.220] | 0.252 |

#### S3.1.2 Table: Sex-difference global TE mediation results

**Table S5:** Sex differences in directly estimated paths (*a*-paths, *b*-paths, and the direct effect) were tested using Satorra–Bentler scaled likelihood ratio tests comparing unconstrained multigroup models with models constraining the parameter to equality across sex. Sex contrasts for derived quantities (specific indirect, total indirect, and total effects) were computed as female *–* male differences from Monte Carlo draws (20,000 draws total), with percentile-based confidence intervals and two-sided *p*-values. Benjamini–Hochberg FDR correction was applied jointly across all 12 sex-difference tests.

| Effect | $\Delta\chi^2$ | df | P | P<sub>FDR</sub> |
| --- | --- | --- | --- | --- |
| *Directly estimated paths (Satorra–Bentler scaled LRT)* |  |  |  |  |
| a-path: EH | 0.109 | 1 | 0.7414 | 0.890 |
| a-path: FSN | 0.509 | 1 | 0.4756 | 0.890 |
| a-path: PMI | 0.181 | 1 | 0.6704 | 0.890 |
| b-path: EH | 14.473 | 1 | $1.42 \times 10^{-4}$ | **0.002** |
| b-path: FSN | 0.174 | 1 | 0.6764 | 0.890 |
| b-path: PMI | 0.005 | 1 | 0.9443 | 0.944 |
| Direct ($c'$) | 0.674 | 1 | 0.4117 | 0.890 |
|  | Estimate | MC 95% CI | P | P<sub>FDR</sub> |
| *Derived quantities: female – male contrast (Monte Carlo CI)* |  |  |  |  |
| EH indirect | 0.130 | [0.061, 0.206] | $3.00 \times 10^{-4}$ | **0.002** |
| FSN indirect | 0.014 | [-0.042, 0.071] | 0.6136 | 0.890 |
| PMI indirect | -0.005 | [-0.069, 0.058] | 0.8629 | 0.941 |
| Total indirect | 0.139 | [0.024, 0.257] | 0.0170 | 0.057 |
| Total effect | 0.231 | [0.040, 0.419] | 0.0190 | 0.057 |

#### S3.1.3 Global total effect: decomposition by absolute path magnitude

We summarized the relative contribution of each path to the parallel decomposition of the family-history effect on global TE when magnitudes are compared on an absolute scale. On each Monte Carlo draw, we computed the absolute contribution of the direct path (*c^′^*) and of each indirect path via the EH, FSN, and PMI components, expressed as a share of the sum of those absolute contributions; shares therefore sum to 100% within sex. The bar height reports the Monte Carlo mean of that percentage. When a path’s estimate is opposite in sign to the estimate of the total effect, the displayed bar and its 95% Monte Carlo confidence interval are given a negative sign so that direction relative to the total effect is visible while preserving the magnitude interpretation of the bar length.

**Figure S13:**
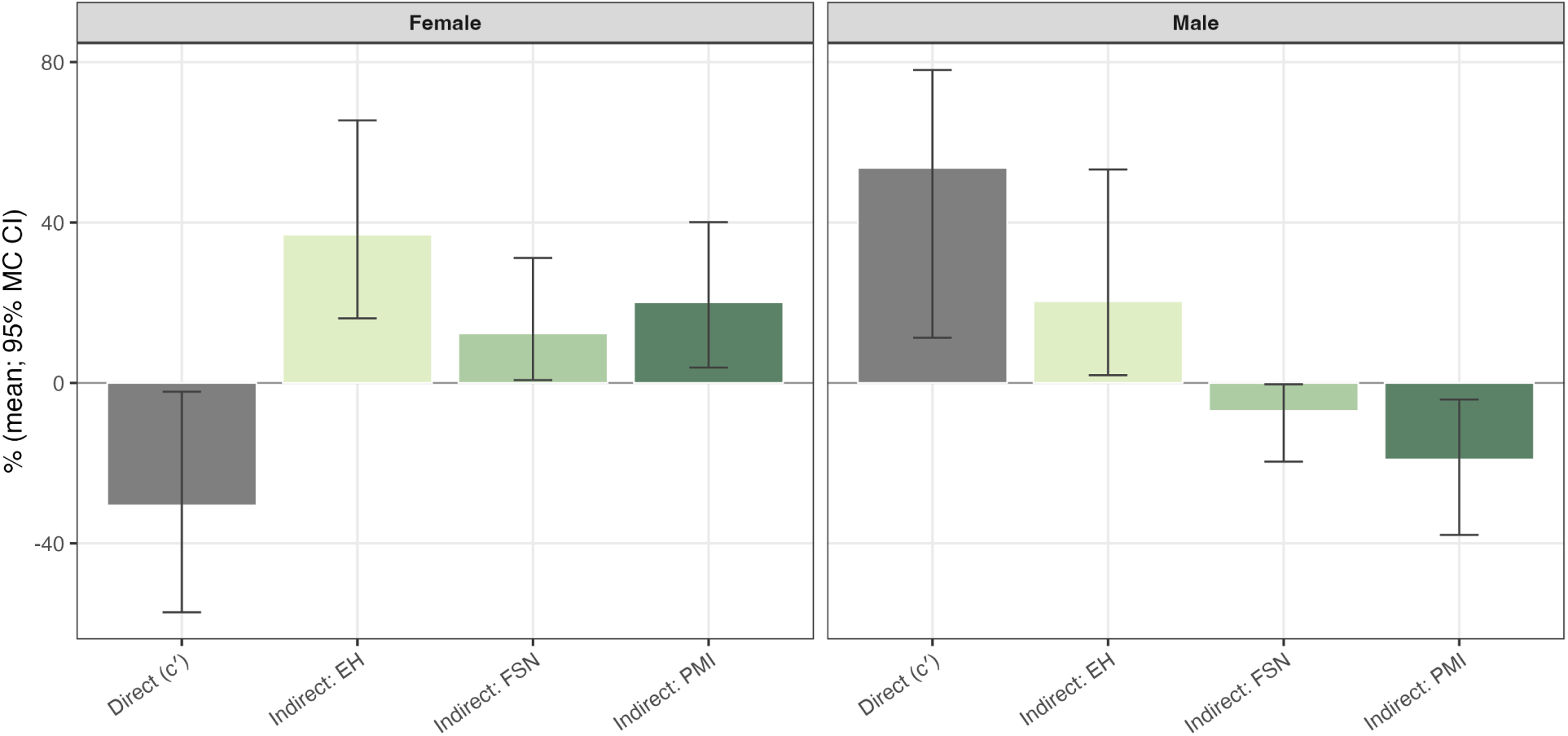
Global TE mediation decomposition by absolute path shares (female vs. male). Bars show the mean percentage share of each path’s absolute contribution relative to the sum of absolute contributions across the direct path (*c^′^*) and the three indirect paths (EH, FSN, PMI) for the global TE outcome. Error bars are 95% Monte Carlo confidence intervals. A negative bar indicates that the path’s estimate is opposite in sign to the total effect.

### S3.2 Network TE Mediation Analysis

#### S3.2.1 Network TE mediation results

**Table S6:**
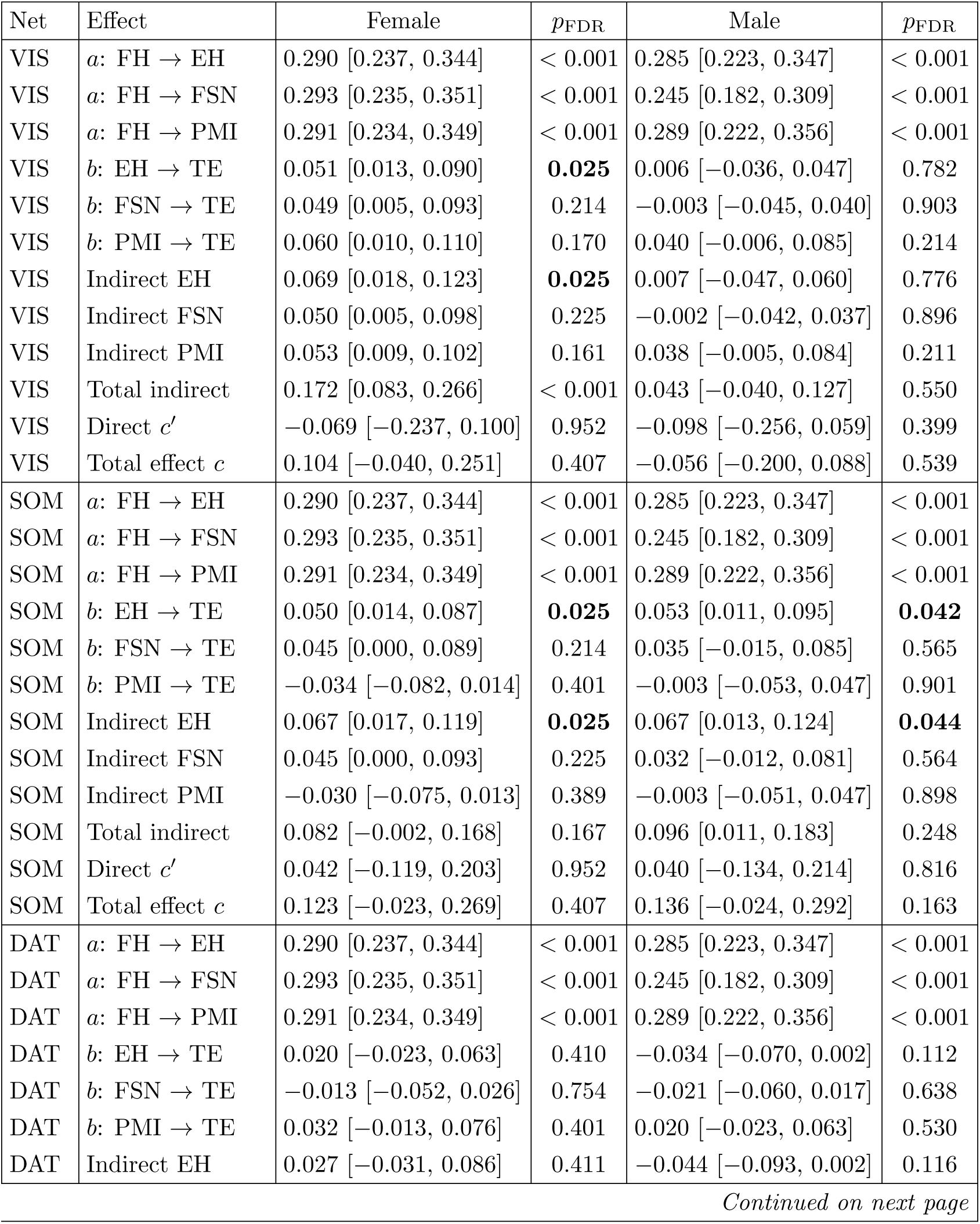
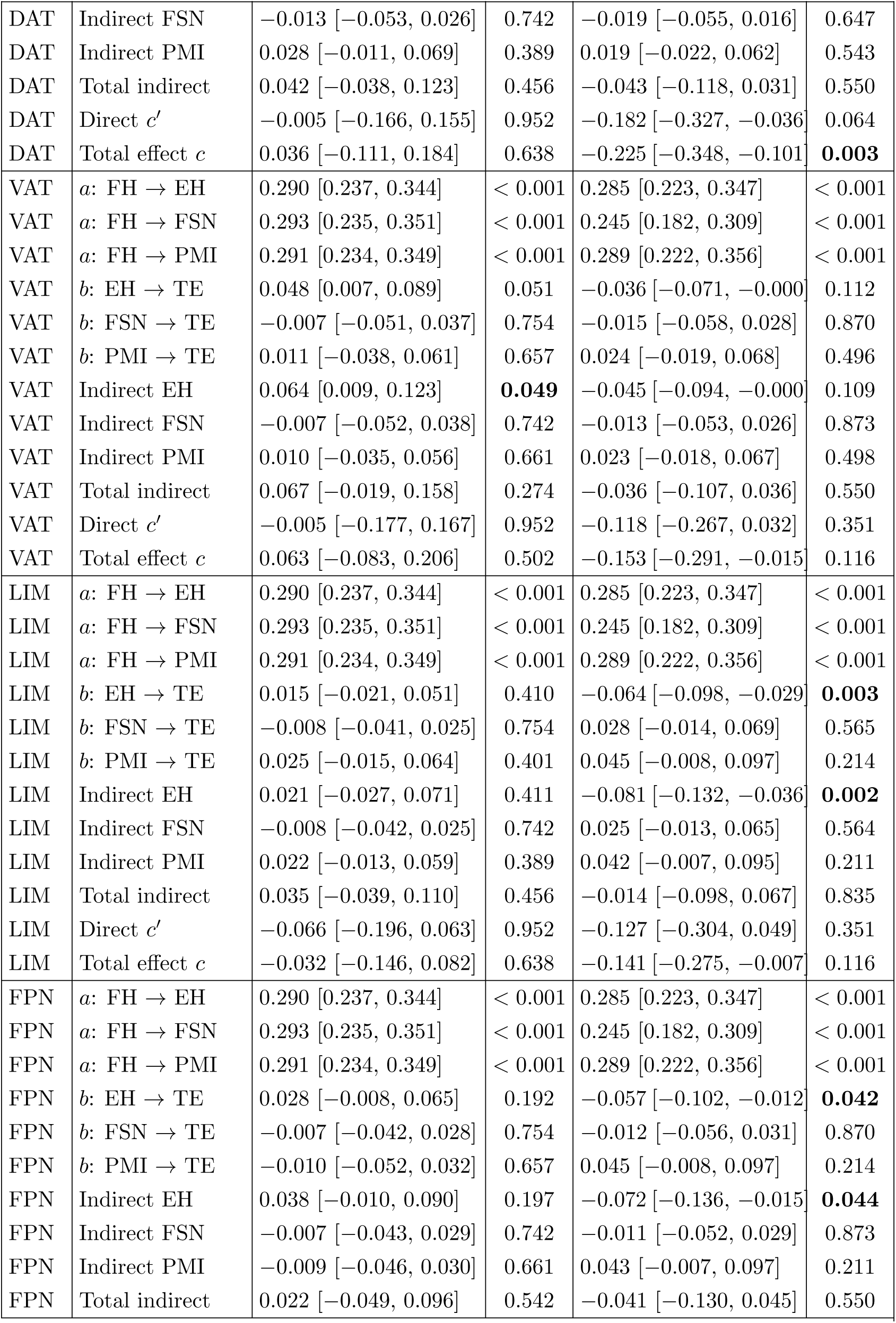
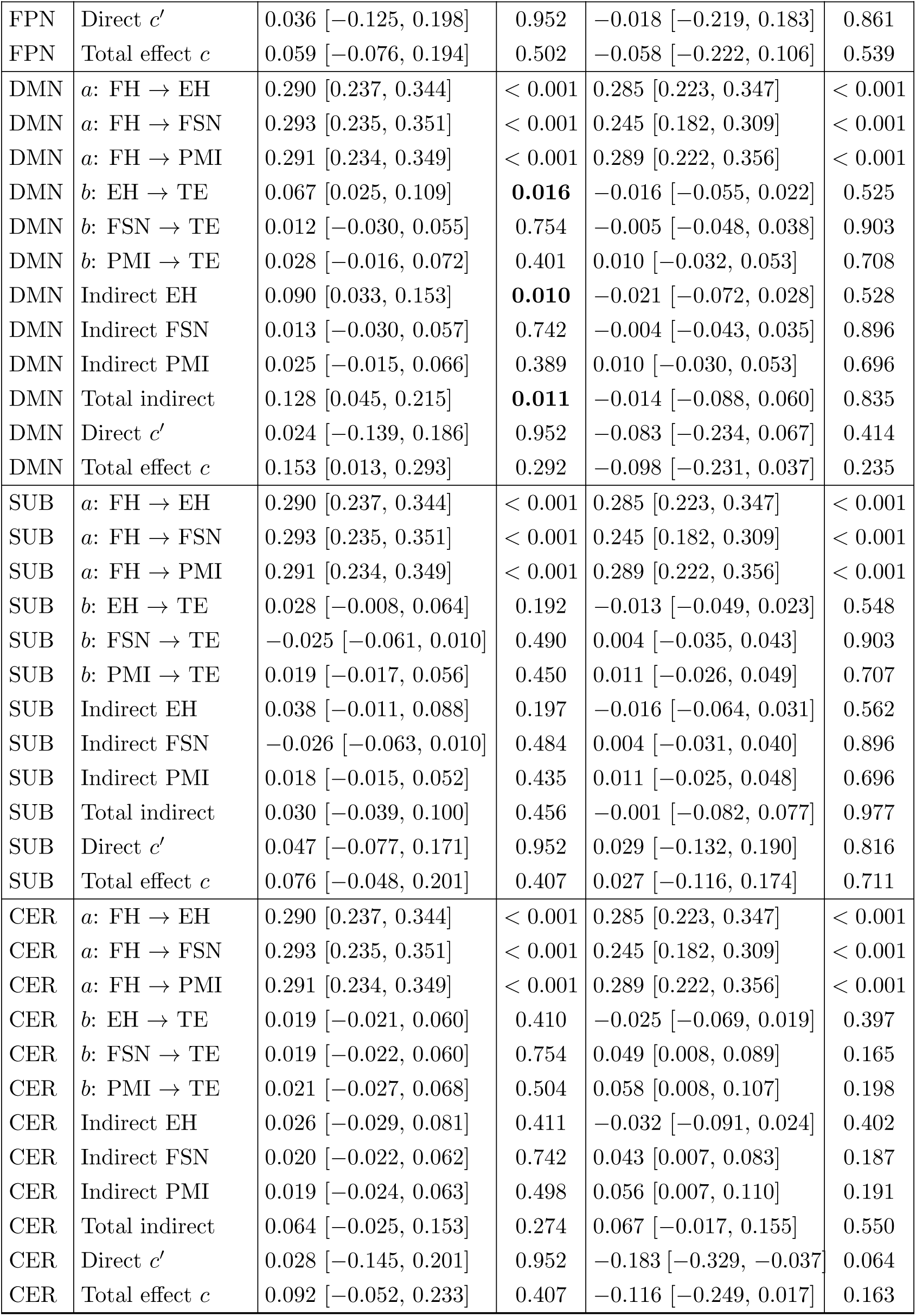
Network TE mediation estimates and 95% CIs. Fully standardized coefficients are shown for *a*-paths (FH *→* mediator); *b*-paths, direct effects, indirect effects, and total effects are on the fitted model scale. *p*_FDR_: Benjamini–Hochberg adjustment across the nine predefined networks (Yeo 7 networks plus subcortical and cerebellar networks) within each effect class and sex (*n* = 9 tests per family)

| Net | Effect | Female | $p_{\text{FDR}}$ | Male | $p_{\text{FDR}}$ |
| --- | --- | --- | --- | --- | --- |
| VIS | $a$: FH → EH | 0.290 [0.237, 0.344] | < 0.001 | 0.285 [0.223, 0.347] | < 0.001 |
| VIS | $a$: FH → FSN | 0.293 [0.235, 0.351] | < 0.001 | 0.245 [0.182, 0.309] | < 0.001 |
| VIS | $a$: FH → PMI | 0.291 [0.234, 0.349] | < 0.001 | 0.289 [0.222, 0.356] | < 0.001 |
| VIS | $b$: EH → TE | 0.051 [0.013, 0.090] | **0.025** | 0.006 [−0.036, 0.047] | 0.782 |
| VIS | $b$: FSN → TE | 0.049 [0.005, 0.093] | 0.214 | −0.003 [−0.045, 0.040] | 0.903 |
| VIS | $b$: PMI → TE | 0.060 [0.010, 0.110] | 0.170 | 0.040 [−0.006, 0.085] | 0.214 |
| VIS | Indirect EH | 0.069 [0.018, 0.123] | **0.025** | 0.007 [−0.047, 0.060] | 0.776 |
| VIS | Indirect FSN | 0.050 [0.005, 0.098] | 0.225 | −0.002 [−0.042, 0.037] | 0.896 |
| VIS | Indirect PMI | 0.053 [0.009, 0.102] | 0.161 | 0.038 [−0.005, 0.084] | 0.211 |
| VIS | Total indirect | 0.172 [0.083, 0.266] | < 0.001 | 0.043 [−0.040, 0.127] | 0.550 |
| VIS | Direct $c'$ | −0.069 [−0.237, 0.100] | 0.952 | −0.098 [−0.256, 0.059] | 0.399 |
| VIS | Total effect $c$ | 0.104 [−0.040, 0.251] | 0.407 | −0.056 [−0.200, 0.088] | 0.539 |
| SOM | $a$: FH → EH | 0.290 [0.237, 0.344] | < 0.001 | 0.285 [0.223, 0.347] | < 0.001 |
| SOM | $a$: FH → FSN | 0.293 [0.235, 0.351] | < 0.001 | 0.245 [0.182, 0.309] | < 0.001 |
| SOM | $a$: FH → PMI | 0.291 [0.234, 0.349] | < 0.001 | 0.289 [0.222, 0.356] | < 0.001 |
| SOM | $b$: EH → TE | 0.050 [0.014, 0.087] | **0.025** | 0.053 [0.011, 0.095] | **0.042** |
| SOM | $b$: FSN → TE | 0.045 [0.000, 0.089] | 0.214 | 0.035 [−0.015, 0.085] | 0.565 |
| SOM | $b$: PMI → TE | −0.034 [−0.082, 0.014] | 0.401 | −0.003 [−0.053, 0.047] | 0.901 |
| SOM | Indirect EH | 0.067 [0.017, 0.119] | **0.025** | 0.067 [0.013, 0.124] | **0.044** |
| SOM | Indirect FSN | 0.045 [0.000, 0.093] | 0.225 | 0.032 [−0.012, 0.081] | 0.564 |
| SOM | Indirect PMI | −0.030 [−0.075, 0.013] | 0.389 | −0.003 [−0.051, 0.047] | 0.898 |
| SOM | Total indirect | 0.082 [−0.002, 0.168] | 0.167 | 0.096 [0.011, 0.183] | 0.248 |
| SOM | Direct $c'$ | 0.042 [−0.119, 0.203] | 0.952 | 0.040 [−0.134, 0.214] | 0.816 |
| SOM | Total effect $c$ | 0.123 [−0.023, 0.269] | 0.407 | 0.136 [−0.024, 0.292] | 0.163 |
| DAT | $a$: FH → EH | 0.290 [0.237, 0.344] | < 0.001 | 0.285 [0.223, 0.347] | < 0.001 |
| DAT | $a$: FH → FSN | 0.293 [0.235, 0.351] | < 0.001 | 0.245 [0.182, 0.309] | < 0.001 |
| DAT | $a$: FH → PMI | 0.291 [0.234, 0.349] | < 0.001 | 0.289 [0.222, 0.356] | < 0.001 |
| DAT | $b$: EH → TE | 0.020 [−0.023, 0.063] | 0.410 | −0.034 [−0.070, 0.002] | 0.112 |
| DAT | $b$: FSN → TE | −0.013 [−0.052, 0.026] | 0.754 | −0.021 [−0.060, 0.017] | 0.638 |
| DAT | $b$: PMI → TE | 0.032 [−0.013, 0.076] | 0.401 | 0.020 [−0.023, 0.063] | 0.530 |
| DAT | Indirect EH | 0.027 [−0.031, 0.086] | 0.411 | −0.044 [−0.093, 0.002] | 0.116 |
| DAT | Indirect FSN | −0.013 [−0.053, 0.026] | 0.742 | −0.019 [−0.055, 0.016] | 0.647 |
| DAT | Indirect PMI | 0.028 [−0.011, 0.069] | 0.389 | 0.019 [−0.022, 0.062] | 0.543 |
| DAT | Total indirect | 0.042 [−0.038, 0.123] | 0.456 | −0.043 [−0.118, 0.031] | 0.550 |
| DAT | Direct $c'$ | −0.005 [−0.166, 0.155] | 0.952 | −0.182 [−0.327, −0.036] | 0.064 |
| DAT | Total effect $c$ | 0.036 [−0.111, 0.184] | 0.638 | −0.225 [−0.348, −0.101] | **0.003** |
| VAT | $a$: FH → EH | 0.290 [0.237, 0.344] | < 0.001 | 0.285 [0.223, 0.347] | < 0.001 |
| VAT | $a$: FH → FSN | 0.293 [0.235, 0.351] | < 0.001 | 0.245 [0.182, 0.309] | < 0.001 |
| VAT | $a$: FH → PMI | 0.291 [0.234, 0.349] | < 0.001 | 0.289 [0.222, 0.356] | < 0.001 |
| VAT | $b$: EH → TE | 0.048 [0.007, 0.089] | 0.051 | −0.036 [−0.071, −0.000] | 0.112 |
| VAT | $b$: FSN → TE | −0.007 [−0.051, 0.037] | 0.754 | −0.015 [−0.058, 0.028] | 0.870 |
| VAT | $b$: PMI → TE | 0.011 [−0.038, 0.061] | 0.657 | 0.024 [−0.019, 0.068] | 0.496 |
| VAT | Indirect EH | 0.064 [0.009, 0.123] | **0.049** | −0.045 [−0.094, −0.000] | 0.109 |
| VAT | Indirect FSN | −0.007 [−0.052, 0.038] | 0.742 | −0.013 [−0.053, 0.026] | 0.873 |
| VAT | Indirect PMI | 0.010 [−0.035, 0.056] | 0.661 | 0.023 [−0.018, 0.067] | 0.498 |
| VAT | Total indirect | 0.067 [−0.019, 0.158] | 0.274 | −0.036 [−0.107, 0.036] | 0.550 |
| VAT | Direct $c'$ | −0.005 [−0.177, 0.167] | 0.952 | −0.118 [−0.267, 0.032] | 0.351 |
| VAT | Total effect $c$ | 0.063 [−0.083, 0.206] | 0.502 | −0.153 [−0.291, −0.015] | 0.116 |
| LIM | $a$: FH → EH | 0.290 [0.237, 0.344] | < 0.001 | 0.285 [0.223, 0.347] | < 0.001 |
| LIM | $a$: FH → FSN | 0.293 [0.235, 0.351] | < 0.001 | 0.245 [0.182, 0.309] | < 0.001 |
| LIM | $a$: FH → PMI | 0.291 [0.234, 0.349] | < 0.001 | 0.289 [0.222, 0.356] | < 0.001 |
| LIM | $b$: EH → TE | 0.015 [−0.021, 0.051] | 0.410 | −0.064 [−0.098, −0.029] | **0.003** |
| LIM | $b$: FSN → TE | −0.008 [−0.041, 0.025] | 0.754 | 0.028 [−0.014, 0.069] | 0.565 |
| LIM | $b$: PMI → TE | 0.025 [−0.015, 0.064] | 0.401 | 0.045 [−0.008, 0.097] | 0.214 |
| LIM | Indirect EH | 0.021 [−0.027, 0.071] | 0.411 | −0.081 [−0.132, −0.036] | **0.002** |
| LIM | Indirect FSN | −0.008 [−0.042, 0.025] | 0.742 | 0.025 [−0.013, 0.065] | 0.564 |
| LIM | Indirect PMI | 0.022 [−0.013, 0.059] | 0.389 | 0.042 [−0.007, 0.095] | 0.211 |
| LIM | Total indirect | 0.035 [−0.039, 0.110] | 0.456 | −0.014 [−0.098, 0.067] | 0.835 |
| LIM | Direct $c'$ | −0.066 [−0.196, 0.063] | 0.952 | −0.127 [−0.304, 0.049] | 0.351 |
| LIM | Total effect $c$ | −0.032 [−0.146, 0.082] | 0.638 | −0.141 [−0.275, −0.007] | 0.116 |
| FPN | $a$: FH → EH | 0.290 [0.237, 0.344] | < 0.001 | 0.285 [0.223, 0.347] | < 0.001 |
| FPN | $a$: FH → FSN | 0.293 [0.235, 0.351] | < 0.001 | 0.245 [0.182, 0.309] | < 0.001 |
| FPN | $a$: FH → PMI | 0.291 [0.234, 0.349] | < 0.001 | 0.289 [0.222, 0.356] | < 0.001 |
| FPN | $b$: EH → TE | 0.028 [−0.008, 0.065] | 0.192 | −0.057 [−0.102, −0.012] | **0.042** |
| FPN | $b$: FSN → TE | −0.007 [−0.042, 0.028] | 0.754 | −0.012 [−0.056, 0.031] | 0.870 |
| FPN | $b$: PMI → TE | −0.010 [−0.052, 0.032] | 0.657 | 0.045 [−0.008, 0.097] | 0.214 |
| FPN | Indirect EH | 0.038 [−0.010, 0.090] | 0.197 | −0.072 [−0.136, −0.015] | **0.044** |
| FPN | Indirect FSN | −0.007 [−0.043, 0.029] | 0.742 | −0.011 [−0.052, 0.029] | 0.873 |
| FPN | Indirect PMI | −0.009 [−0.046, 0.030] | 0.661 | 0.043 [−0.007, 0.097] | 0.211 |
| FPN | Total indirect | 0.022 [−0.049, 0.096] | 0.542 | −0.041 [−0.130, 0.045] | 0.550 |

| Net | Effect | Female | $p_{FDR}$ | Male | $p_{FDR}$ |
| --- | --- | --- | --- | --- | --- |
| FPN | Direct $c'$ | 0.036 [−0.125, 0.198] | 0.952 | −0.018 [−0.219, 0.183] | 0.861 |
| FPN | Total effect $c$ | 0.059 [−0.076, 0.194] | 0.502 | −0.058 [−0.222, 0.106] | 0.539 |
| DMN | $a$: FH → EH | 0.290 [0.237, 0.344] | < 0.001 | 0.285 [0.223, 0.347] | < 0.001 |
| DMN | $a$: FH → FSN | 0.293 [0.235, 0.351] | < 0.001 | 0.245 [0.182, 0.309] | < 0.001 |
| DMN | $a$: FH → PMI | 0.291 [0.234, 0.349] | < 0.001 | 0.289 [0.222, 0.356] | < 0.001 |
| DMN | $b$: EH → TE | 0.067 [0.025, 0.109] | **0.016** | −0.016 [−0.055, 0.022] | 0.525 |
| DMN | $b$: FSN → TE | 0.012 [−0.030, 0.055] | 0.754 | −0.005 [−0.048, 0.038] | 0.903 |
| DMN | $b$: PMI → TE | 0.028 [−0.016, 0.072] | 0.401 | 0.010 [−0.032, 0.053] | 0.708 |
| DMN | Indirect EH | 0.090 [0.033, 0.153] | **0.010** | −0.021 [−0.072, 0.028] | 0.528 |
| DMN | Indirect FSN | 0.013 [−0.030, 0.057] | 0.742 | −0.004 [−0.043, 0.035] | 0.896 |
| DMN | Indirect PMI | 0.025 [−0.015, 0.066] | 0.389 | 0.010 [−0.030, 0.053] | 0.696 |
| DMN | Total indirect | 0.128 [0.045, 0.215] | **0.011** | −0.014 [−0.088, 0.060] | 0.835 |
| DMN | Direct $c'$ | 0.024 [−0.139, 0.186] | 0.952 | −0.083 [−0.234, 0.067] | 0.414 |
| DMN | Total effect $c$ | 0.153 [0.013, 0.293] | 0.292 | −0.098 [−0.231, 0.037] | 0.235 |
| SUB | $a$: FH → EH | 0.290 [0.237, 0.344] | < 0.001 | 0.285 [0.223, 0.347] | < 0.001 |
| SUB | $a$: FH → FSN | 0.293 [0.235, 0.351] | < 0.001 | 0.245 [0.182, 0.309] | < 0.001 |
| SUB | $a$: FH → PMI | 0.291 [0.234, 0.349] | < 0.001 | 0.289 [0.222, 0.356] | < 0.001 |
| SUB | $b$: EH → TE | 0.028 [−0.008, 0.064] | 0.192 | −0.013 [−0.049, 0.023] | 0.548 |
| SUB | $b$: FSN → TE | −0.025 [−0.061, 0.010] | 0.490 | 0.004 [−0.035, 0.043] | 0.903 |
| SUB | $b$: PMI → TE | 0.019 [−0.017, 0.056] | 0.450 | 0.011 [−0.026, 0.049] | 0.707 |
| SUB | Indirect EH | 0.038 [−0.011, 0.088] | 0.197 | −0.016 [−0.064, 0.031] | 0.562 |
| SUB | Indirect FSN | −0.026 [−0.063, 0.010] | 0.484 | 0.004 [−0.031, 0.040] | 0.896 |
| SUB | Indirect PMI | 0.018 [−0.015, 0.052] | 0.435 | 0.011 [−0.025, 0.048] | 0.696 |
| SUB | Total indirect | 0.030 [−0.039, 0.100] | 0.456 | −0.001 [−0.082, 0.077] | 0.977 |
| SUB | Direct $c'$ | 0.047 [−0.077, 0.171] | 0.952 | 0.029 [−0.132, 0.190] | 0.816 |
| SUB | Total effect $c$ | 0.076 [−0.048, 0.201] | 0.407 | 0.027 [−0.116, 0.174] | 0.711 |
| CER | $a$: FH → EH | 0.290 [0.237, 0.344] | < 0.001 | 0.285 [0.223, 0.347] | < 0.001 |
| CER | $a$: FH → FSN | 0.293 [0.235, 0.351] | < 0.001 | 0.245 [0.182, 0.309] | < 0.001 |
| CER | $a$: FH → PMI | 0.291 [0.234, 0.349] | < 0.001 | 0.289 [0.222, 0.356] | < 0.001 |
| CER | $b$: EH → TE | 0.019 [−0.021, 0.060] | 0.410 | −0.025 [−0.069, 0.019] | 0.397 |
| CER | $b$: FSN → TE | 0.019 [−0.022, 0.060] | 0.754 | 0.049 [0.008, 0.089] | 0.165 |
| CER | $b$: PMI → TE | 0.021 [−0.027, 0.068] | 0.504 | 0.058 [0.008, 0.107] | 0.198 |
| CER | Indirect EH | 0.026 [−0.029, 0.081] | 0.411 | −0.032 [−0.091, 0.024] | 0.402 |
| CER | Indirect FSN | 0.020 [−0.022, 0.062] | 0.742 | 0.043 [0.007, 0.083] | 0.187 |
| CER | Indirect PMI | 0.019 [−0.024, 0.063] | 0.498 | 0.056 [0.007, 0.110] | 0.191 |
| CER | Total indirect | 0.064 [−0.025, 0.153] | 0.274 | 0.067 [−0.017, 0.155] | 0.550 |
| CER | Direct $c'$ | 0.028 [−0.145, 0.201] | 0.952 | −0.183 [−0.329, −0.037] | 0.064 |
| CER | Total effect $c$ | 0.092 [−0.052, 0.233] | 0.407 | −0.116 [−0.249, 0.017] | 0.163 |

#### S3.2.2 Sex-differences in network TE mediation

**Table S7:**
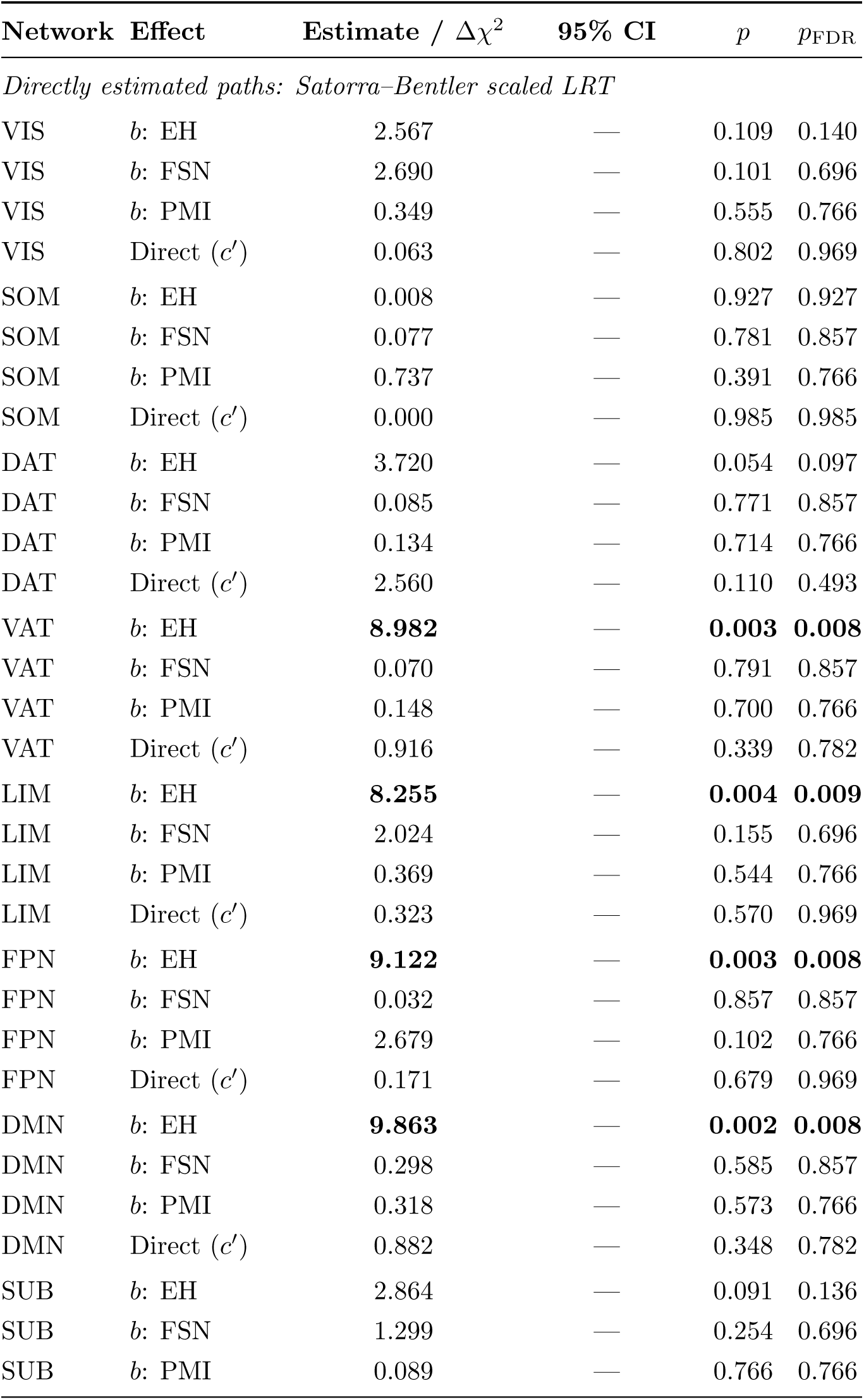
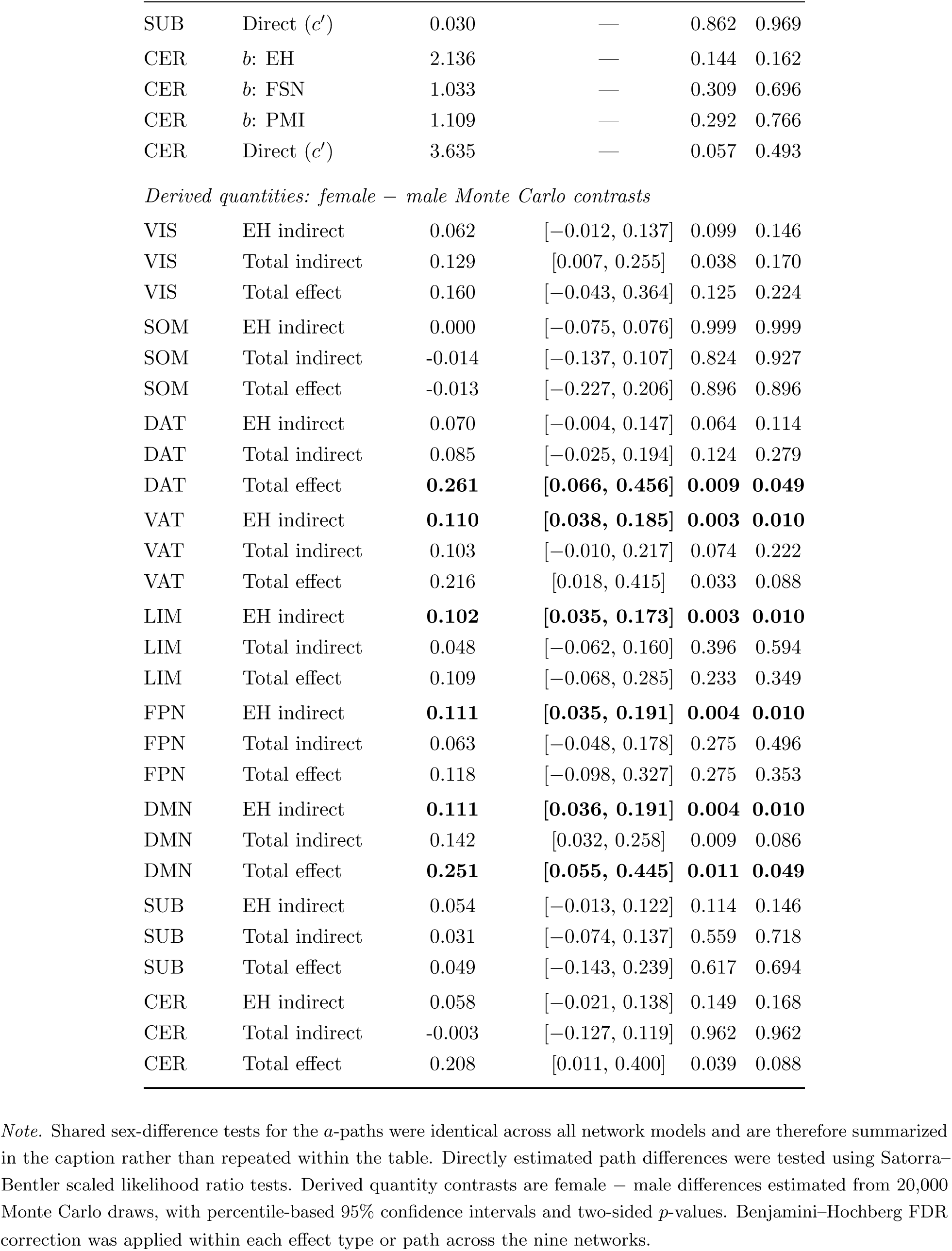
Sex-difference results for network transition energy mediation. Shared *a*-path sex-difference tests were omitted because they were identical across networks (EH: Δ*χ*^2^ = 0.109, *p* = 0.741; FSN: Δ*χ*^2^ = 0.509, *p* = 0.476; PMI: Δ*χ*^2^ = 0.181, *p* = 0.670). Shown here are network-specific sex-difference tests for directly estimated paths (Satorra–Bentler scaled LRTs) and derived quantities (female *−* male Monte Carlo contrasts).

| Network | Effect | Estimate / $\Delta\chi^2$ | 95% CI | $p$ | $p_{\text{FDR}}$ |
| --- | --- | --- | --- | --- | --- |
| *Directly estimated paths: Satorra–Bentler scaled LRT* |  |  |  |  |  |
| VIS | $b$: EH | 2.567 | — | 0.109 | 0.140 |
| VIS | $b$: FSN | 2.690 | — | 0.101 | 0.696 |
| VIS | $b$: PMI | 0.349 | — | 0.555 | 0.766 |
| VIS | Direct ($c'$) | 0.063 | — | 0.802 | 0.969 |
| SOM | $b$: EH | 0.008 | — | 0.927 | 0.927 |
| SOM | $b$: FSN | 0.077 | — | 0.781 | 0.857 |
| SOM | $b$: PMI | 0.737 | — | 0.391 | 0.766 |
| SOM | Direct ($c'$) | 0.000 | — | 0.985 | 0.985 |
| DAT | $b$: EH | 3.720 | — | 0.054 | 0.097 |
| DAT | $b$: FSN | 0.085 | — | 0.771 | 0.857 |
| DAT | $b$: PMI | 0.134 | — | 0.714 | 0.766 |
| DAT | Direct ($c'$) | 2.560 | — | 0.110 | 0.493 |
| VAT | $b$: EH | **8.982** | — | **0.003** | **0.008** |
| VAT | $b$: FSN | 0.070 | — | 0.791 | 0.857 |
| VAT | $b$: PMI | 0.148 | — | 0.700 | 0.766 |
| VAT | Direct ($c'$) | 0.916 | — | 0.339 | 0.782 |
| LIM | $b$: EH | **8.255** | — | **0.004** | **0.009** |
| LIM | $b$: FSN | 2.024 | — | 0.155 | 0.696 |
| LIM | $b$: PMI | 0.369 | — | 0.544 | 0.766 |
| LIM | Direct ($c'$) | 0.323 | — | 0.570 | 0.969 |
| FPN | $b$: EH | **9.122** | — | **0.003** | **0.008** |
| FPN | $b$: FSN | 0.032 | — | 0.857 | 0.857 |
| FPN | $b$: PMI | 2.679 | — | 0.102 | 0.766 |
| FPN | Direct ($c'$) | 0.171 | — | 0.679 | 0.969 |
| DMN | $b$: EH | **9.863** | — | **0.002** | **0.008** |
| DMN | $b$: FSN | 0.298 | — | 0.585 | 0.857 |
| DMN | $b$: PMI | 0.318 | — | 0.573 | 0.766 |
| DMN | Direct ($c'$) | 0.882 | — | 0.348 | 0.782 |
| SUB | $b$: EH | 2.864 | — | 0.091 | 0.136 |
| SUB | $b$: FSN | 1.299 | — | 0.254 | 0.696 |
| SUB | $b$: PMI | 0.089 | — | 0.766 | 0.766 |
| SUB | Direct ($c'$) | 0.030 | — | 0.862 | 0.969 |
| CER | $b$: EH | 2.136 | — | 0.144 | 0.162 |
| CER | $b$: FSN | 1.033 | — | 0.309 | 0.696 |
| CER | $b$: PMI | 1.109 | — | 0.292 | 0.766 |
| CER | Direct ($c'$) | 3.635 | — | 0.057 | 0.493 |
| *Derived quantities: female – male Monte Carlo contrasts* |  |  |  |  |  |
| VIS | EH indirect | 0.062 | [−0.012, 0.137] | 0.099 | 0.146 |
| VIS | Total indirect | 0.129 | [0.007, 0.255] | 0.038 | 0.170 |
| VIS | Total effect | 0.160 | [−0.043, 0.364] | 0.125 | 0.224 |
| SOM | EH indirect | 0.000 | [−0.075, 0.076] | 0.999 | 0.999 |
| SOM | Total indirect | −0.014 | [−0.137, 0.107] | 0.824 | 0.927 |
| SOM | Total effect | −0.013 | [−0.227, 0.206] | 0.896 | 0.896 |
| DAT | EH indirect | 0.070 | [−0.004, 0.147] | 0.064 | 0.114 |
| DAT | Total indirect | 0.085 | [−0.025, 0.194] | 0.124 | 0.279 |
| DAT | Total effect | **0.261** | **[0.066, 0.456]** | **0.009** | **0.049** |
| VAT | EH indirect | **0.110** | **[0.038, 0.185]** | **0.003** | **0.010** |
| VAT | Total indirect | 0.103 | [−0.010, 0.217] | 0.074 | 0.222 |
| VAT | Total effect | 0.216 | [0.018, 0.415] | 0.033 | 0.088 |
| LIM | EH indirect | **0.102** | **[0.035, 0.173]** | **0.003** | **0.010** |
| LIM | Total indirect | 0.048 | [−0.062, 0.160] | 0.396 | 0.594 |
| LIM | Total effect | 0.109 | [−0.068, 0.285] | 0.233 | 0.349 |
| FPN | EH indirect | **0.111** | **[0.035, 0.191]** | **0.004** | **0.010** |
| FPN | Total indirect | 0.063 | [−0.048, 0.178] | 0.275 | 0.496 |
| FPN | Total effect | 0.118 | [−0.098, 0.327] | 0.275 | 0.353 |
| DMN | EH indirect | **0.111** | **[0.036, 0.191]** | **0.004** | **0.010** |
| DMN | Total indirect | 0.142 | [0.032, 0.258] | 0.009 | 0.086 |
| DMN | Total effect | **0.251** | **[0.055, 0.445]** | **0.011** | **0.049** |
| SUB | EH indirect | 0.054 | [−0.013, 0.122] | 0.114 | 0.146 |
| SUB | Total indirect | 0.031 | [−0.074, 0.137] | 0.559 | 0.718 |
| SUB | Total effect | 0.049 | [−0.143, 0.239] | 0.617 | 0.694 |
| CER | EH indirect | 0.058 | [−0.021, 0.138] | 0.149 | 0.168 |
| CER | Total indirect | −0.003 | [−0.127, 0.119] | 0.962 | 0.962 |
| CER | Total effect | 0.208 | [0.011, 0.400] | 0.039 | 0.088 |

*Note.* Shared sex-difference tests for the $a$ -paths were identical across all network models and are therefore summarized in the caption rather than repeated within the table. Directly estimated path differences were tested using Satorra–Bentler scaled likelihood ratio tests. Derived quantity contrasts are female – male differences estimated from 20,000 Monte Carlo draws, with percentile-based 95% confidence intervals and two-sided $p$ -values. Benjamini–Hochberg FDR
correction was applied within each effect type or path across the nine networks.

### S3.3 Regional TE Mediation Analysis

#### S3.3.1 Indirect effects for three environmental mediators on regional TE brain plots

**Figure S14:**
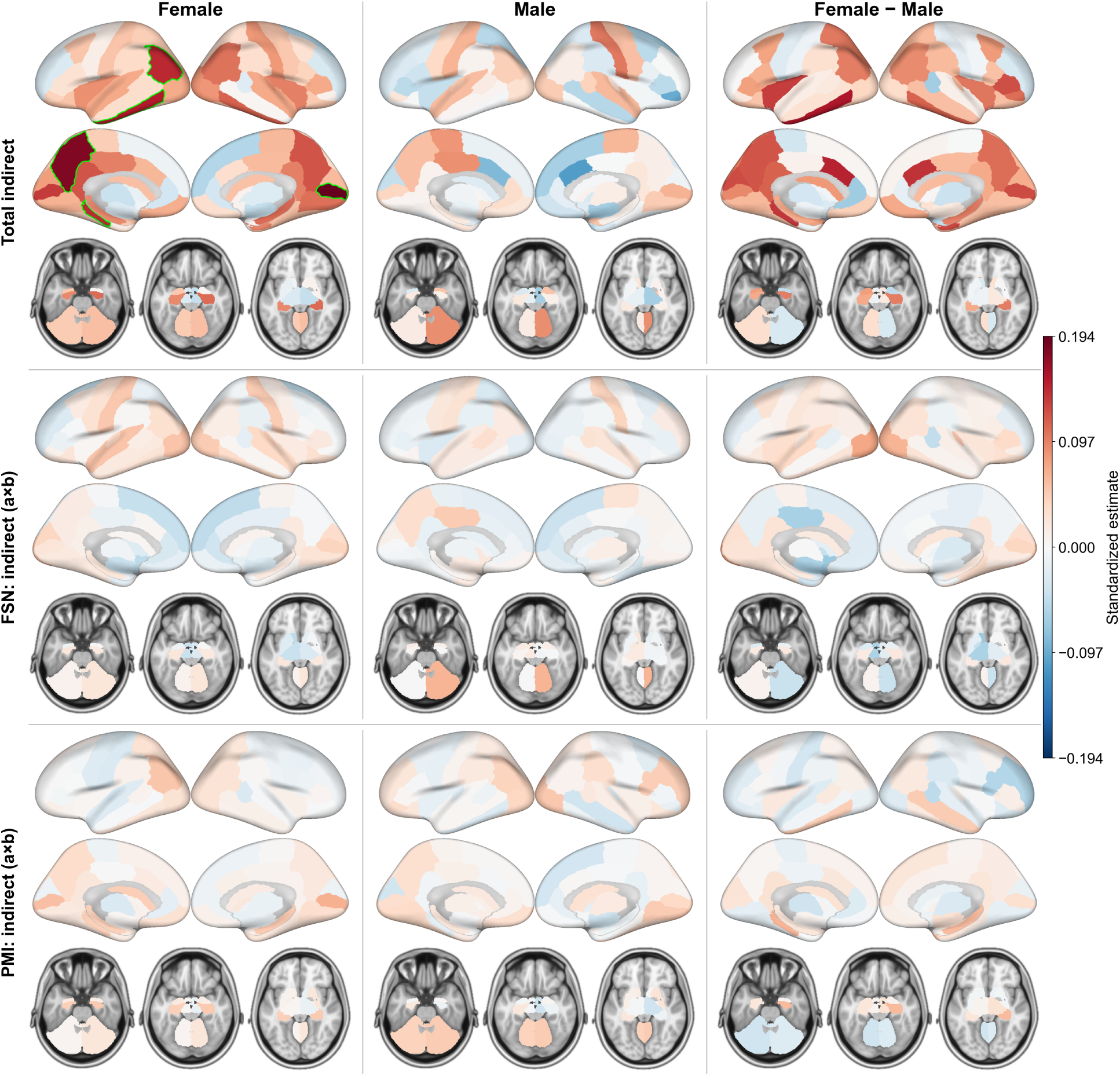
Region-wise total and specific indirect effects of family history on transition energy in each region of the FS86 parcellation. Rows correspond to the total indirect effect, the specific indirect effect of family-school neglect (FSN), and the specific indirect effect of parental mental illness (PMI); columns show effects in females, males, and the female minus male contrast. Color scale is symmetric (red = positive, blue = negative) with limits shared across all panels in this figure. Regions with effects significant after FDR correction are outlined in green.

#### S3.3.2 *b*-path associations of three environmental mediators with regional TE

**Figure S15:**
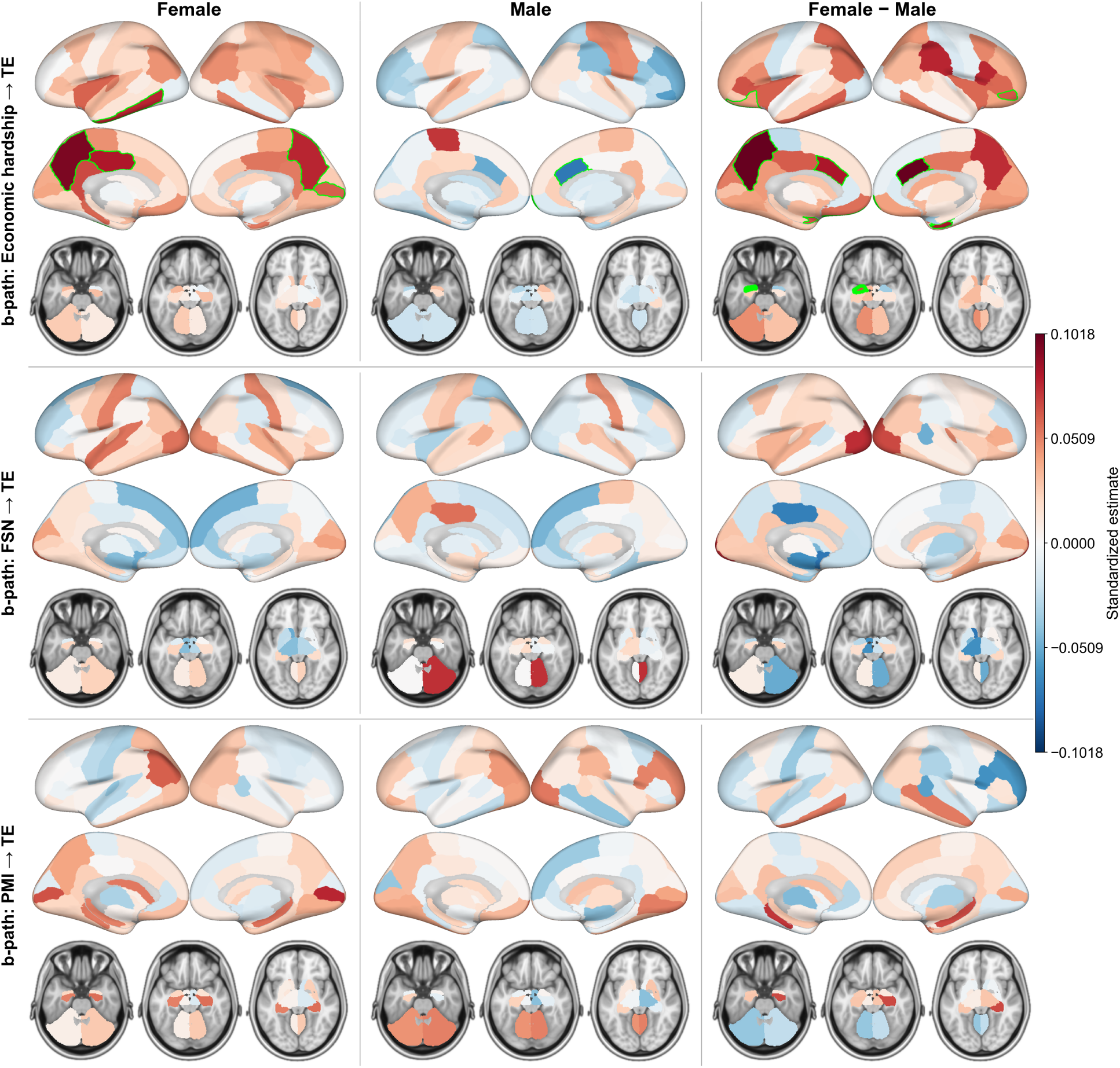
Region-wise *b*-path associations between each environmental mediator and transition energy of each region of the FS86 parcellation. Rows correspond to economic hardship (EH) *→* TE (PC1), family-school neglect (FSN) *→* TE (PC2), and parental mental illness (PMI) *→* TE (PC3) *b*-paths; columns show estimates in females, males, and the female minus male contrast. Maps use the same symmetric color scale for all panels in this figure (red = positive, blue = negative). Regions with effects significant after FDR correction are outlined in green.

#### S3.3.3 Significant regional effects in males and females

**Table S8:** Significant mediation effects on regional transition energies in males and females.

| Sex | Effect type | Region name | Estimate | *p* | *p*<sub>FDR</sub> |
| --- | --- | --- | --- | --- | --- |
| Female | EH b-path | ctx-lh-inferiortemporal | 0.077 | < 0.001 | 0.011 |
| Female | EH b-path | ctx-lh-posteriorcingulate | 0.083 | < 0.001 | 0.008 |
| Female | EH b-path | ctx-lh-precuneus | 0.096 | < 0.001 | 0.011 |
| Female | EH b-path | ctx-rh-pericalcarine | 0.061 | 0.003 | 0.047 |
| Female | EH b-path | ctx-rh-precuneus | 0.077 | 0.003 | 0.047 |
| Female | EH indirect | ctx-lh-inferiortemporal | 0.104 | < 0.001 | 0.014 |
| Female | EH indirect | ctx-lh-posteriorcingulate | 0.111 | < 0.001 | 0.014 |
| Female | EH indirect | ctx-lh-precuneus | 0.129 | < 0.001 | 0.014 |
| Female | Total indirect | ctx-lh-inferiorparietal | 0.146 | < 0.001 | 0.009 |
| Female | Total indirect | ctx-lh-inferiortemporal | 0.154 | 0.003 | 0.050 |
| Female | Total indirect | ctx-lh-parahippocampal | 0.145 | < 0.001 | 0.006 |
| Female | Total indirect | ctx-lh-precuneus | 0.180 | < 0.001 | < 0.001 |
| Female | Total indirect | ctx-rh-pericalcarine | 0.194 | < 0.001 | < 0.001 |
| Male | EH b-path | ctx-rh-caudalanteriorcingulate | -0.073 | < 0.001 | 0.008 |
| Male | EH b-path | ctx-rh-frontalpole | -0.072 | < 0.001 | 0.011 |
| Male | EH indirect | ctx-rh-caudalanteriorcingulate | -0.092 | < 0.001 | < 0.001 |
| Male | EH indirect | ctx-rh-frontalpole | -0.091 | < 0.001 | 0.017 |
| Male | Total effect | ctx-lh-superiorparietal | -0.250 | < 0.001 | < 0.001 |

#### S3.3.4 Significant sex differences in regional effects

**Table S9:** Significant Female*−*Male contrasts of mediation effects on regional transition energies.

| Contrast | Region name | Estimate | Test | $p$ | $p_{\text{FDR}}$ |
| --- | --- | --- | --- | --- | --- |
| EH indirect | ctx-lh-caudalanteriorcingulate | 0.108 | MC | 0.002 | 0.041 |
| EH indirect | ctx-lh-precuneus | 0.135 | MC | 0.001 | 0.041 |
| EH indirect | ctx-rh-caudalanteriorcingulate | 0.132 | MC | < 0.001 | 0.017 |
| EH indirect | ctx-rh-entorhinal | 0.111 | MC | 0.002 | 0.041 |
| b-path (EH) | Left-Amygdala | 8.395 | LRT | 0.004 | 0.040 |
| b-path (EH) | ctx-lh-caudalanteriorcingulate | 11.075 | LRT | < 0.001 | 0.019 |
| b-path (EH) | ctx-lh-lateralorbitofrontal | 10.144 | LRT | 0.001 | 0.025 |
| b-path (EH) | ctx-lh-precuneus | 11.570 | LRT | < 0.001 | 0.019 |
| b-path (EH) | ctx-rh-caudalanteriorcingulate | 16.161 | LRT | < 0.001 | 0.005 |
| b-path (EH) | ctx-rh-entorhinal | 12.284 | LRT | < 0.001 | 0.019 |
| b-path (EH) | ctx-rh-frontalpole | 9.255 | LRT | 0.002 | 0.034 |
| b-path (EH) | ctx-rh-parsorbitalis | 8.638 | LRT | 0.003 | 0.040 |

#### S3.3.5 Multiple comparison correction

Multiple-comparison correction was applied hierarchically according to both analytic level and inferential class. This strategy was designed to control false discovery across comparable sets of hypotheses while avoiding overly conservative pooling across analytically distinct questions. The three analytic levels (i.e., global, network, and regional) were treated separately and Benjamini-Hochberg false discovery rate (FDR) correction was not applied across all tests in the project simultaneously, but rather within prespecified families of comparable tests at each analytic level.

Within each analytic level, correction families were defined by effect type. Path-level parameters and mediation-effect estimates were treated separately, because they represent distinct inferential targets. Specifically, the *a*-paths, *b*-paths, and direct effects (*c^′^*) were grouped as path-level tests, as these are constituent model parameters estimated directly within the structural equation model. In contrast, the specific indirect effects (*a × b*), the total indirect effect, and the total effect were grouped as mediation-effect tests, as these are derived quantities. Formal sex-difference tests were treated as a third inferential class.

At the global level, FDR correction was applied separately to: (i) all within-sex path-level tests, including all *a*-paths, *b*-paths, and direct effects estimated in females and males; (ii) mediation-effect tests pooled across both sexes (10 tests: three specific indirect effects, the total indirect effect, and the total effect in females and males); and (iii) all formal sex-difference tests for global TE, including sex contrasts for directly estimated paths and sex contrasts for indirect, total indirect, and total effects.

At the network level, FDR correction was applied across the 9 networks within each inferential family. Thus, for each sex separately, all network-specific tests for a given path-level parameter (e.g., the EH *b*-path in females across the 9 networks) were corrected together, and the same logic was applied to all other *a*-paths, *b*-paths, and direct effects. Likewise, mediation-effect tests were corrected across the 9 networks within each sex and effect type (e.g., the EH-specific indirect effect in males across the 9 networks; the total indirect effect in females across the 9 networks; the total effect in males across the 9 networks). Sex-difference tests at the network level were corrected separately across the 9 networks within each contrast type (e.g., the sex difference in the EH indirect effect across the 9 networks).

At the regional level, the same logic was extended across the 86 regions. For each sex separately, all regional tests for a given path-level parameter were corrected together across regions, and all regional tests for a given mediation-effect quantity were likewise corrected together across regions. For example, the female EH *b*-path was corrected across all 86 regions as one family, the male EH-specific indirect effect was corrected across all 86 regions as another family, and the regional sex-difference test for the total indirect effect was corrected across all 86 regions as its own family. Overall, this hierarchical framework reflects a balance between rigor and interpretability.

### S3.4 Puberty-moderated multi-group SEM

In the mediation analytic sample (*N* = 1,835), we compared raw Pubertal Development Scale (PDS) scores between FH+ and FH*−* youth separately by sex using Welch two-sample *t*-tests (unequal variances). Among females, mean PDS was significantly higher in FH+ (*M* = 2.23, s.d. = 0.93, *n* = 277) than in FH*−* (*M* = 2.09, s.d. = 0.90, *n* = 721; *t*(485.0) = 2.16, *p* = 0.031). Among males, mean PDS was slightly higher in FH+ (*M* = 1.34, s.d. = 0.59, *n* = 214) than in FH*−* (*M* = 1.29, s.d. = 0.54, *n* = 623), but the difference was not significant (*t*(341.9) = 1.21, *p* = 0.226). After stratifying by pubertal status (not started vs. started), cell sizes were: FH*−* males (*n* = 470 pre-pubertal, *n* = 153 pubertal), FH+ males (*n* = 154, *n* = 60), FH*−* females (*n* = 242, *n* = 479), and FH+ females (*n* = 83, *n* = 194).

In order to test whether pubertal status moderated the relationship between family history, environmental dimensions, and TE, we conducted a parallel mediation analysis in which binary pubertal status (*not started* vs. *started*) moderated all paths in the mediation model (family history *→* environmental PCs *→* TE), while retaining sex stratification via a two-group (female/male) specification as in the main analyses. Models were estimated with maximum likelihood and robust standard errors, with family ID as a clustering variable for cluster-robust SEs. Path coefficients (including moderation terms) were assessed with Wald tests from the fitted model; conditional indirect, direct (*c^′^*), and total effects, and female–male contrasts of those conditional quantities, used Monte Carlo 95% confidence intervals and two-sided *p*-values based on the asymptotic multivariate sampling distribution of the estimated parameters.

Interpretation of these puberty-moderated models is limited by the uneven distribution of pubertal status across groups, which reduced sample size after stratification and increased uncertainty in the estimated paths. In females, the positive total effect of family history on global TE remained directionally positive in both pre-pubertal and pubertal youth, but neither stratum was statistically significant. In males, by contrast, the negative total effect of family history on global TE appeared to be attenuated under pubertal conditions, with positive indirect effects through environmental variables offsetting more of the negative direct effect than in the pre-pubertal condition. This pattern should be interpreted cautiously, as the number of pubertal males—particularly FH+ pubertal males—was limited. Although underpowered, these exploratory results suggest that pubertal status may shape the expression of family-history and environmental associations with global TE.

**Figure S16:**
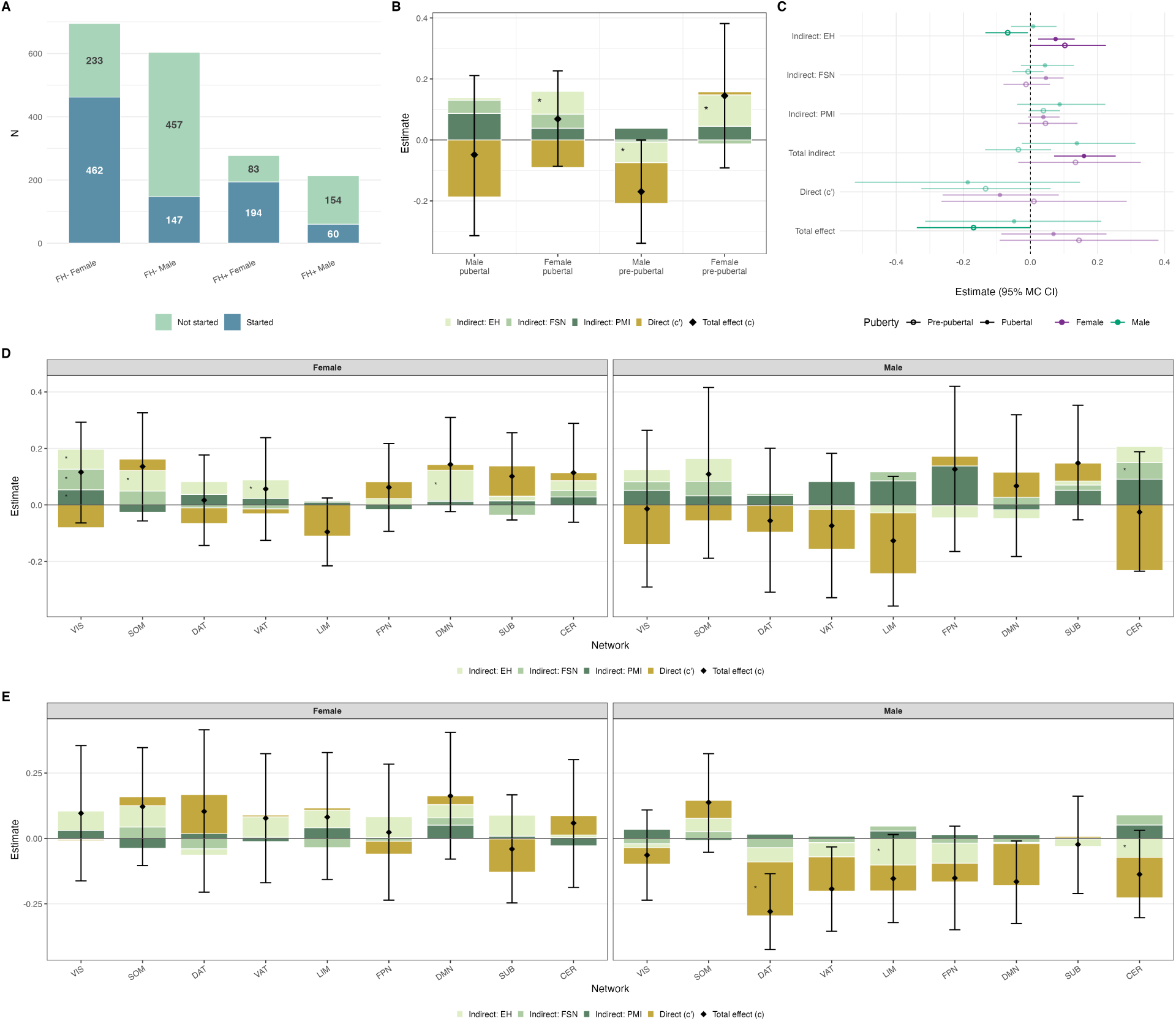
Puberty-moderated parallel mediation for global and network TE. **(A)** Sample distribution of puberty status (not started vs. started) within each family-history-by-sex group.**(B)** Decomposition of the conditional effect of family history on global TE separately for four sex-by-puberty groups (male pubertal, female pubertal, male pre-pubertal, female pre-pubertal): stacked bars show the three specific indirect pathways (economic hardship, family–school neglect, parental mental illness), the direct path, and the total effect (diamond with Monte Carlo 95% confidence interval). Asterisks indicate components for which the Monte Carlo interval excludes zero. **(C)** Forest plot of conditional effects on global TE across sex and puberty status (pre-pubertal vs. pubertal), including specific indirect, total indirect, direct, and total effects; non-significant estimates are shown with reduced opacity. **(D)** Same decomposition as panel (B) for each Yeo network outcome in the pubertal subgroup, faceted by sex. **(E)** Same network-level decomposition as panel (D) for the pre-pubertal subgroup.

### S3.5 Sensitivity Analysis: Complete-case mediation

The mediation analyses used environmental PCA scores (PCs 1–3) merged for all subjects in the subsample, some of whom had PCA inputs imputed (site-level imputation) prior to PCA when subject-level data were missing on individual input variables. As a sensitivity check, we repeated the parallel mediation model restricted to participants with complete data on all variables entering the environmental PCA. After applying this restriction, the mediation sample was *N* = 1,250 (FH+ male *n* = 144; FH+ female *n* = 161; FH*−* male *n* = 445; FH*−* female *n* = 500). The overall direction and pattern of effects were broadly consistent with the primary analysis, although some estimates were attenuated, likely reflecting reduced power due to a reduced N in each subgroup. Missingness is likely not random here and subjects with higher missingness may represent the most at-risk youth.

### S3.6 Policy-linked projection: supplementary results

Supplementary Table S10. reports coefficients from the joint reduced-domain policy → EH model (full ABCD cohort with environmental PC scores, *N* = 9,157). Of the five FH*×*Policy interaction terms, three survived Benjamini–Hochberg FDR correction: Good Samaritan law, recreational cannabis legalization, and PDMP must-access requirements. Each was associated with a smaller FH+ versus FH*−* EH gap in states where the policy was present (negative *β*_FH_*_×_*_Policy_). ACA Medicaid expansion showed a positive, non-significant interaction (larger EH gap where present), and naloxone access showed a small negative, non-significant interaction.

Supplementary Table S11. reports the projected global TE consequences of these EH-gap associations, obtained by propagating each FH*×*Policy coefficient through the sex-specific EH*→*TE *b*-path from the mediation models (Monte Carlo median across *N*_MC_ = 20,000 draws). Because the observed FH+ minus FH*−* global TE gap was positive in females (+0.077 SD) but negative in males (*−*0.168 SD), a policy-associated reduction in the EH gap is projected to move the TE gap toward zero from opposite directions in each sex: downward in females and upward (i.e., less negative) in males. The three FDR-significant policy domains individually projected partial gap closure in both sexes, ranging from 19.5% to 25.8% in females and from 5.5% to 7.2% in males. Naloxone access showed a smaller projected closure but did not show an FDR-significant FH*×*Policy interaction and was not included in the combined scenario. In the additive combined scenario summing all three significant FH*×*Policy coefficients, the projected global TE gap closed by 72.4% in females and 20.7% in males relative to the observed gap. ACA expansion, whose interaction effect was directionally opposite to the three significant policies, projected a small widening of the TE gap in both sexes and was not included in the combined scenario.

Projected effects at the network and regional level, based on the same policy-linked projection outputs, were more heterogeneous in both direction and magnitude than the global estimate, and several outcomes had observed FH TE gaps near zero (|gap*| <* 0.02 SD), for which percent gap closure was not interpreted (see Methods). These network- and regional-level projections should be interpreted as exploratory and are not corrected for multiple comparisons across outcomes.

**Table S10:**
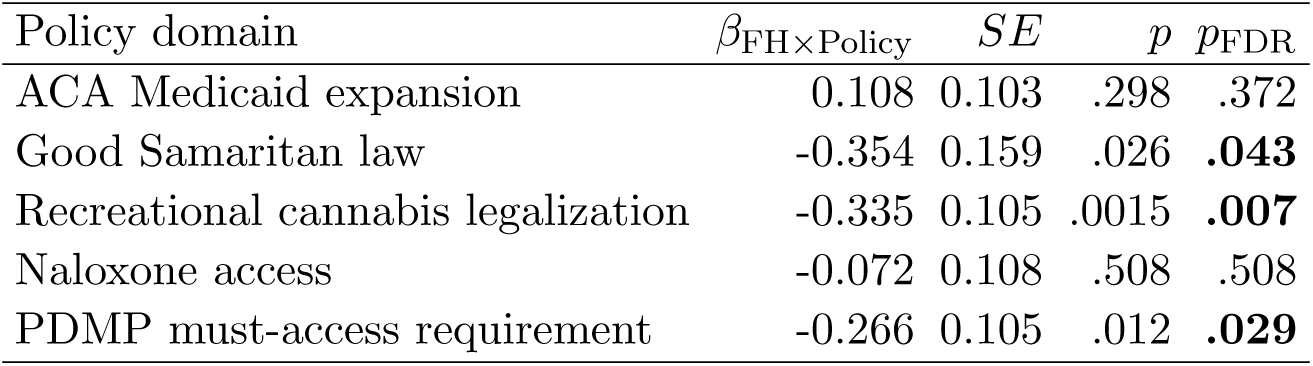
Policy *×* FH interaction effects on economic hardship (EH; joint reduced model, *N* = 9,157).

**Figure S17:**
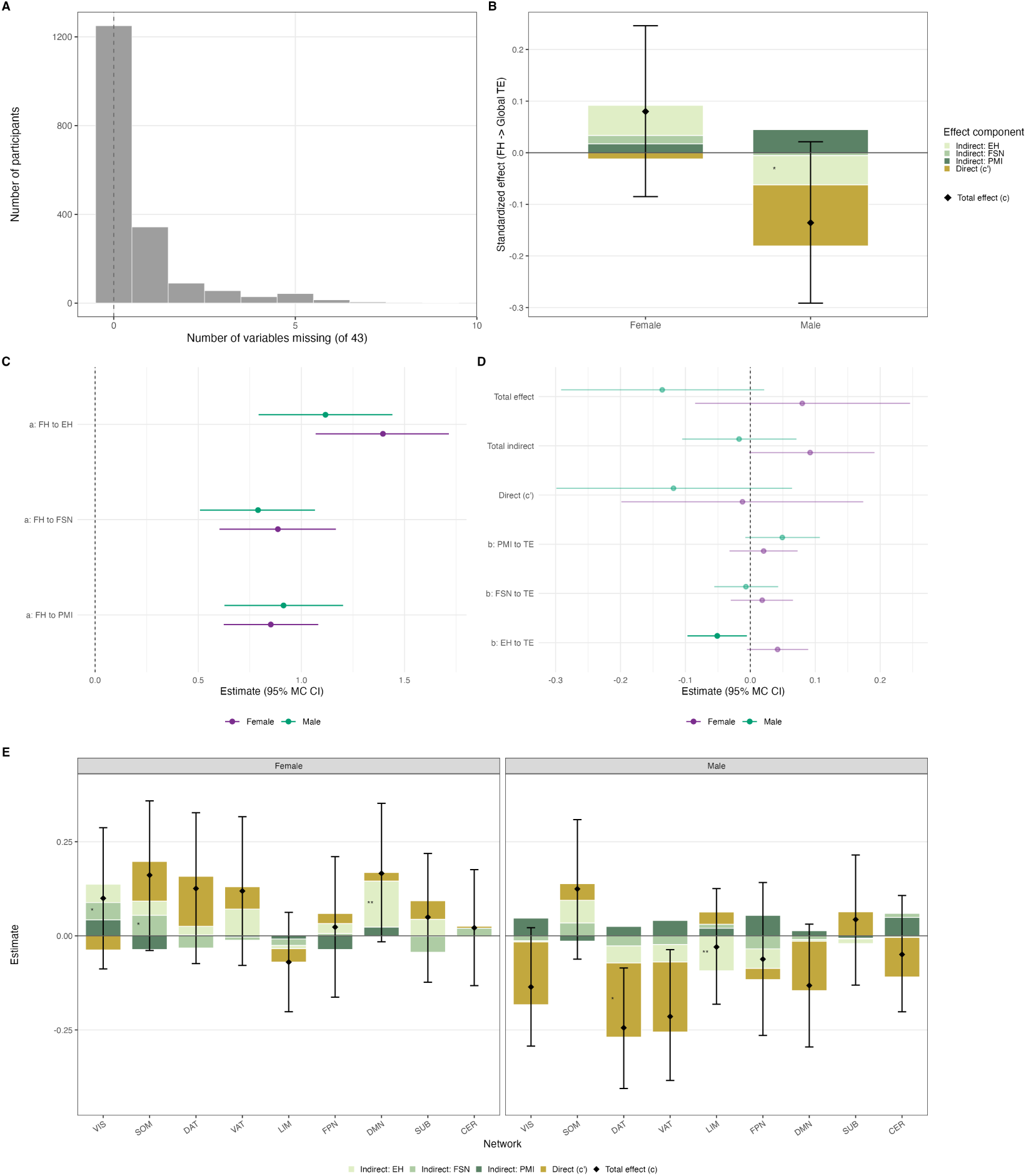
Sensitivity mediation analysis restricted to participants with complete data on all environment variables used in the PCA. **(A)** Histogram of how many of the 43 PCA input variables were missing per participant in the full mediation analytic cohort before applying the complete-case filter. **(B)** Decomposition of the family-history effect on global TE: stacked direct (*c^′^*) and indirect effects via PC1–3 (EH, FSN, PMI), with total effect (black diamond) and 95% Monte Carlo confidence intervals, by sex. **(C)** Forest plot of *a*-paths (family history *→* each environmental mediator). **(D)** Forest plot of *b*-paths (each mediator *→* global TE), total indirect effect, direct effect (*c^′^*), and total effect. **(E)** Network-level decomposition of the family-history effect on TE (Yeo 7-networks plus subcortical and cerebellar networks), with the same component coding as in panel B, faceted by sex. *^∗^*Monte Carlo *p <* 0.05; *^∗∗^*Benjamini–Hochberg FDR *p <* 0.05 where applicable.

**Table S11:**
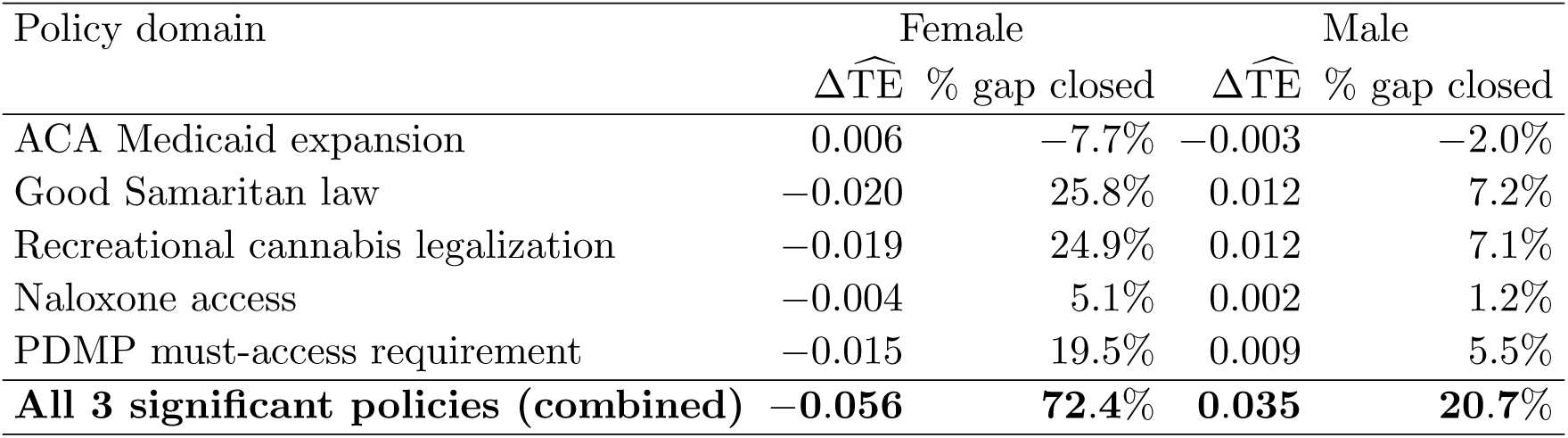
Projected global TE gap change by policy domain (Monte Carlo median, *N*_MC_ = 20,000 draws). Observed FH+ minus FH*−* global TE gap in the imaging cohort was +0.077 SD (female) and *−*0.168 SD (male).

| Policy domain | Female |  | Male |  |
| --- | --- | --- | --- | --- |
| | $\Delta\widehat{TE}$ | % gap closed | $\Delta\widehat{TE}$ | % gap closed |
| ACA Medicaid expansion | 0.006 | −7.7% | −0.003 | −2.0% |
| Good Samaritan law | −0.020 | 25.8% | 0.012 | 7.2% |
| Recreational cannabis legalization | −0.019 | 24.9% | 0.012 | 7.1% |
| Naloxone access | −0.004 | 5.1% | 0.002 | 1.2% |
| PDMP must-access requirement | −0.015 | 19.5% | 0.009 | 5.5% |
| **All 3 significant policies (combined)** | **−0.056** | **72.4%** | **0.035** | **20.7%** |

